# Upstream regulator of genomic imprinting in rice endosperm is a small RNA-associated chromatin remodeler

**DOI:** 10.1101/2023.08.31.555833

**Authors:** Avik Kumar Pal, Vivek Hari Sundar G, Amruta N, P.V. Shivaprasad

## Abstract

Genomic imprinting is observed in endosperm, a placenta-like seed tissue, where transposable elements (TEs) and repeat-derived small(s)RNAs mediate epigenetic changes in plants. In imprinting, uniparental gene expression arises due to parent-specific epigenetic marks on one allele but not on the other. The importance of sRNAs and their regulation in endosperm development or in imprinting is poorly understood in crops. Here we show that a previously uncharacterized CLASSY (CLSY)-family chromatin remodeler named *OsCLSY3* is essential for rice endosperm development and imprinting, acting as an upstream player in sRNA pathway. Comparative transcriptome and genetic analysis indicated its endosperm-preferred expression and its paternally imprinted nature. These important features were modulated by RNA-directed DNA methylation (RdDM) of tandemly arranged TEs in its promoter. Upon perturbation of *OsCLSY3* in transgenic lines we observed defects in endosperm development and loss of around 70% of all sRNAs. Interestingly, well-conserved endosperm-specific sRNAs (siren) that are vital for reproductive fitness in angiosperms were dependent on *OsCLSY3*. We also observed many imprinted genes and seed development-associated genes under the control of CLSY3-dependent RdDM. These results support an essential role of *OsCLSY3* in rice endosperm development and imprinting, and propose similar regulatory strategies involving *CLSY3* homologs among other cereals.

**Highlights:**

1. Unlike among dicots, in rice and maize, *CLSY3* is a maternally expressed imprinted gene majorly expressed in endosperm.
2. Endosperm-preferred expression of *OsCLSY3* is regulated by RNA-directed DNA methylation at two tandem transposon elements present in its promoter.
3. *OsCLSY3* is crucial for endosperm development and grain filling. It regulates expression of key seed development and endosperm-specific imprinted genes through RNA directed DNA methylation.

## Introduction

Seed formation is a major stage in the life cycle of gymnosperms and angiosperms. The embryo is embedded in a nourishment tissue and protected by a seed coat in a typical seed, and such an arrangement protects the embryo against unfavorable environmental conditions and mechanical damages. Unlike gymnosperms, angiosperms seeds evolved a unique nourishment tissue called endosperm which is also a fertilization product as embryo. Among the two haploid sperm nuclei in the pollen, one fertilizes the egg and the other fertilizes two haploid central cells nuclei, in a process called as double fertilization. The egg cell fertilization results in a diploid embryo and the fertilization of the two haploid central cell nuclei by the sperm nuclei results in a triploid endosperm (Pires, 2014; Baroux et al., 2002).

Apart from the nourishment of the developing embryo, endosperm also senses the environmental changes such as temperature, nutrient availability, biotic factors etc. to control seed germination (Chahtane et al., 2018; Iwasaki et al., 2019; De Giorgi et al., 2021; Iwasaki et al., 2022). Endosperm derived signals such as small molecules, hormones, proteins, peptides, RNAs etc. play crucial roles in the development of the embryo (Doll et al., 2020). Besides, endosperm is the major source for carbohydrates, proteins and fat for most of the animals, including humans.

Despite several landmark studies, mechanism of endosperm development is still an obscure area of research (Li and Berger, 2012). Multiple studies in model plant *Arabidopsis* indicates that genetic, hormonal and epigenetic pathways play vital roles in endosperm development. Unique ploidy, genomic imprinting and chromatin organization in endosperm distinguishes it from other plant tissues (Baroux et al., 2002; Gehring, 2013). Imprinting is important since it regulates endosperm development, providing reproductive fitness to the embryo. A gene is imprinted when one of the parental alleles is partially or completely silenced due to allele-specific epigenetic marks in chromatin and the other allele gets expressed. Among the imprinted genes which are paternally silent but maternally expressed are called maternally expressed imprinted genes (MEGs) and which are maternally silent but paternally expressed are referred as paternally expressed imprinted genes (PEGs) (Gehring, 2013; Kiyosue et al., 1999; Kradolfer et al., 2013; Gutierrez-Marcos et al., 2003; Kinoshita, 2007; Yuan et al., 2017).

Epigenetic players such as DNA methylation, demethylation and polycomb group complexes (PcG), processes regulate genomic imprinting through chromatin modifications in *Arabidopsis* (Gehring et al., 2006; Kinoshita et al., 2004; Batista and Köhler, 2020). In agreement with this, mutations in several epigenetic-associated imprinted genes led to defective endosperm and embryos (Ishikawa et al., 2011; Huang et al., 2017; Yuan et al., 2017; Zhu et al., 2018; Cheng et al., 2021). Surprisingly, across plants, most of the major epigenetic regulators of imprinting themselves were imprinted in endosperm, including *AtFIS2, AtMEA, AtVIM5, AtNRPD, OsFIE1, OsEMF2a, ZmFIE1, ZmFIE2* genes. Often, tissue- and allele-specific expression of these major players were also tightly regulated by epigenetic pathways (Luo et al., 1999; Grossniklaus et al., 1998; Ohad et al., 1996; Kiyosue et al., 1999; Waters et al., 2013; Anderson et al., 2021; Chen et al., 2018; Luo et al., 2009; Cheng et al., 2020). Epigenetic pathways play a crucial role in endosperm development by regulating expression of imprinted and development related genes.

DNA methylation, considered as the primary epigenetic mark, is a major contributor to genomic imprinting in plants (Köhler and Weinhofer-Molisch, 2010; Satyaki and Gehring, 2017). The cytosine methylation of DNA is peculiar in plants as it is observed in CG, CHG and CHH contexts (where H corresponds to A, T, or C). Multiple DNA methyltransferases encoded by plants establish and maintain methylation marks. There is a division of labor to maintain these marks, the CG methylation is maintained by METHYLTRANSFERASE1 (MET1); CHG methylation requires CHROMOMETHYLASE3 (CMT3). The CHH methylation is regulated by a *de novo* DNA methyltransferase named DOMAINS REARRANGED METHYLTRANSFERASE2 (DRM2) which is guided by RdDM.

A plant specific DNA-dependent RNA polymerase IV (PolIV) generates small (∼30-40 nt) transcripts predominantly from TEs and repeats to initiate RdDM. These RNAs get converted into double-stranded (ds) forms by RNA DEPENDENT RNA POLYMERASE2 (RDR2). The ds substrates are processed into 23-24 nt sRNA duplexes by DICER-LIKE3 (DCL3). The 24 nt sRNAs are preferentially loaded into ARGONAUTE4/6/9 (AGO4/6/9) to guide DRM2 to methylate TEs and repeats by associating with long non-coding transcripts (150-200 nt) generated by another plant-specific polymerase named PolV. PolIV is specifically recruited to TEs and repeats by a family of SNF2 chromatin remodelers named CLASSY (CLSY) along with SAWADEE HOMEODOMAIN HOMOLOG1 (SHH1) proteins. The tissue-specific expression of CLSYs regulate tissue-specific DNA methylation in *Arabidopsis* (Smith et al., 2007; Zhang et al., 2013; Law et al., 2011; Zhou et al., 2018, 2022; Martins and Law, 2023).

Although RdDM is known to regulate imprinting in *Arabidopsis* (Vu et al., 2013), there seems to be some variations in the way it contributes to genome imprinting across plants. For example, RdDM pathway derived 24 nt sRNAs are majorly maternally biased in the young seeds of *Arabidopsis* (Mosher et al., 2009; Mosher, 2010) and maternally biased sRNAs regulate imprinting and seed development (Lu et al., 2012; Kirkbride et al., 2019). Also, in maize and *Brassica rapa*, 24 nt sRNAs were majorly maternally biased in endosperm (Xin et al., 2014; Grover et al., 2020) whereas in rice, 24 nt sRNAs were derived from both the parental genomes (Rodrigues et al., 2013; Yuan et al., 2017; Chen et al., 2018). Unlike other tissues, a major portion of 24 nt sRNAs were majorly derived from genic regions and not from TEs in rice endosperm (Rodrigues et al., 2013). The same study found a unique class of highly expressing sRNA loci in rice endosperm named <u>siR</u>NA <u>en</u>dosperm-specific (siren) loci. Those loci overlapped with genic and inter-genic regions (Rodrigues et al., 2013). Multiple studies have also revealed the presence of siren loci in *B. rapa* and *Arabidopsis,* both in endosperm and ovule tissues. The siren derived sRNAs not just induced DNA methylation (Grover et al., 2020; Burgess et al., 2022), but also regulates expression of adjacent genes. Siren loci derived sRNAs can also trigger DNA methylation *in trans* at specific protein-coding genes (Burgess et al., 2022; Long et al., 2021), indicating their versatile nature, function of which is not well understood. In rice, maize and *Arabidopsis*, many imprinted sRNA loci are located proximal to the imprinted genes. Surprisingly, silenced alleles of many imprinted genes generates imprinted sRNAs (Rodrigues et al., 2013; Xin et al., 2014; Yuan et al., 2017; Erdmann et al., 2017; Chen et al., 2018).

The early stages of endosperm development are very similar between cereals like rice and dicots like *Arabidopsis*. However, differences are observed in the later stages of development. After the cellularization, cereal endosperm differentiated into two major tissues: the outer aleurone layer and the inner starchy endosperm. The cereal endosperm rapidly accumulates large amounts of storage material which nourishes the embryo during embryogenesis and seed germination. In *Arabidopsis*, endosperm provides nourishment only during embryogenesis. How such variations in development are imposed and maintained in plants are unknown (Olsen, 2020; Liu et al., 2022a) and this variation holds huge commercial and agronomic value. Few imprinting-associated genes were found conserved between dicots and monocots (Rodrigues et al., 2013, 2021; Chen et al., 2018; Luo et al., 2011; Yuan et al., 2017; Waters et al., 2011, 2013), while, the number of conserved imprinted genes were comparatively higher among monocots (Waters et al., 2011, 2013; Anderson et al., 2021). Cereals seem to have many different regulators for endosperm development, and their contributions are unknown.

Here, we used a transcriptome screen to identify epigenetic regulators of endosperm development in rice. A previously unannotated ortholog of *Arabidopsis CLSY3* was identified, that majorly expressed in endosperm and regulated key TE-derived sRNAs, siren loci and imprinted sRNA loci. Using genetic and molecular methods including knockdown (kd) and knockout (*KO*) lines, we show that rice CLSY3-dependent loci regulate several imprinted genes and seed development-related genes. Mutation in *OsCLSY3* negatively affected endosperm formation, whereas its overexpression (OE) led to larger seeds with defective cellularization. We also identified that *OsCLSY3* itself was a maternally expressed imprinted gene (MEG) and its silencing in vegetative tissues was maintained through an RdDM loop operating at TEs in its promoter. These results indicate the presence of tissue-specific roles of paternally imprinted *OsCLSY3* in rice endosperm and its regulation of other genes through RdDM.

## Results

### Os*CLSY3* is an endosperm-preferred imprinted gene

To understand key regulators of endosperm development in rice, we performed a tissue-specific transcriptome analysis using rice seeds. We isolated rice embryo as well as endosperm (20-25 days after pollination-DAP), and performed transcriptomic analysis with biological replicates using *indica* rice variety Pusa Basmati-1 (PB1) as described in methods (Fig. 1A, Supplemental Fig. S1A). More than 90% of the intact reads of more than 30 million reads per library were mapped to rice genome (Supplemental Table S1). Well known tissue-specific marker genes that express either in embryo, such as *OsH1*, *OsARF2*, *OsRS2*, *OsGRF4,* or in endosperm, such as *OsFIE1*, *OsNF-YC11,* showed expected patterns, indicating that the tissues taken for analysis were free from cross-contamination (Supplemental Fig. S1B and C). The green non-seed tissue contamination, considered as a major problem in seed transcriptomes, was not observed in our embryo and endosperm transcriptomes, since green tissue specific genes such a Os12g0516000, Os09g0537700 were not detected (Supplemental Fig. S1D) (Li et al., 2018). To find the endosperm and embryo-preferred genes, we considered log_2_ two-fold upregulated and downregulated genes in endosperm when compared to embryo for further analysis. Around 3607 transcripts showed embryo-preferred expression, while transcripts that expressed highly in endosperm were around 3686 (Supplemental dataset S1 and Supplemental dataset S2). From the gene ontology (GO) analysis, we observed embryo-preferred genes were majorly hormonal pathways, transcription-related functions, development-related while endosperm-preferred genes were related to carbohydrate, protein metabolism and transport-related pathways (Supplemental Fig. S1E and F). The genes that showed endosperm-preferred expression in dicots such as *AP2, WRKY, bHLH, MADS* box, *NAC* transcription factors behaved similarly in rice (Hsieh et al., 2011; Le et al., 2010; Mahto et al., 2017). We also observed multiple carbohydrate storage, starch branching and sugar metabolism related genes such as *OsSGL, OsGBP, SBDCP1*, and *OsFLO2* were uniquely expressed in cereal endosperm (Supplemental dataset S1, Supplemental dataset S2).

**Figure 1.**
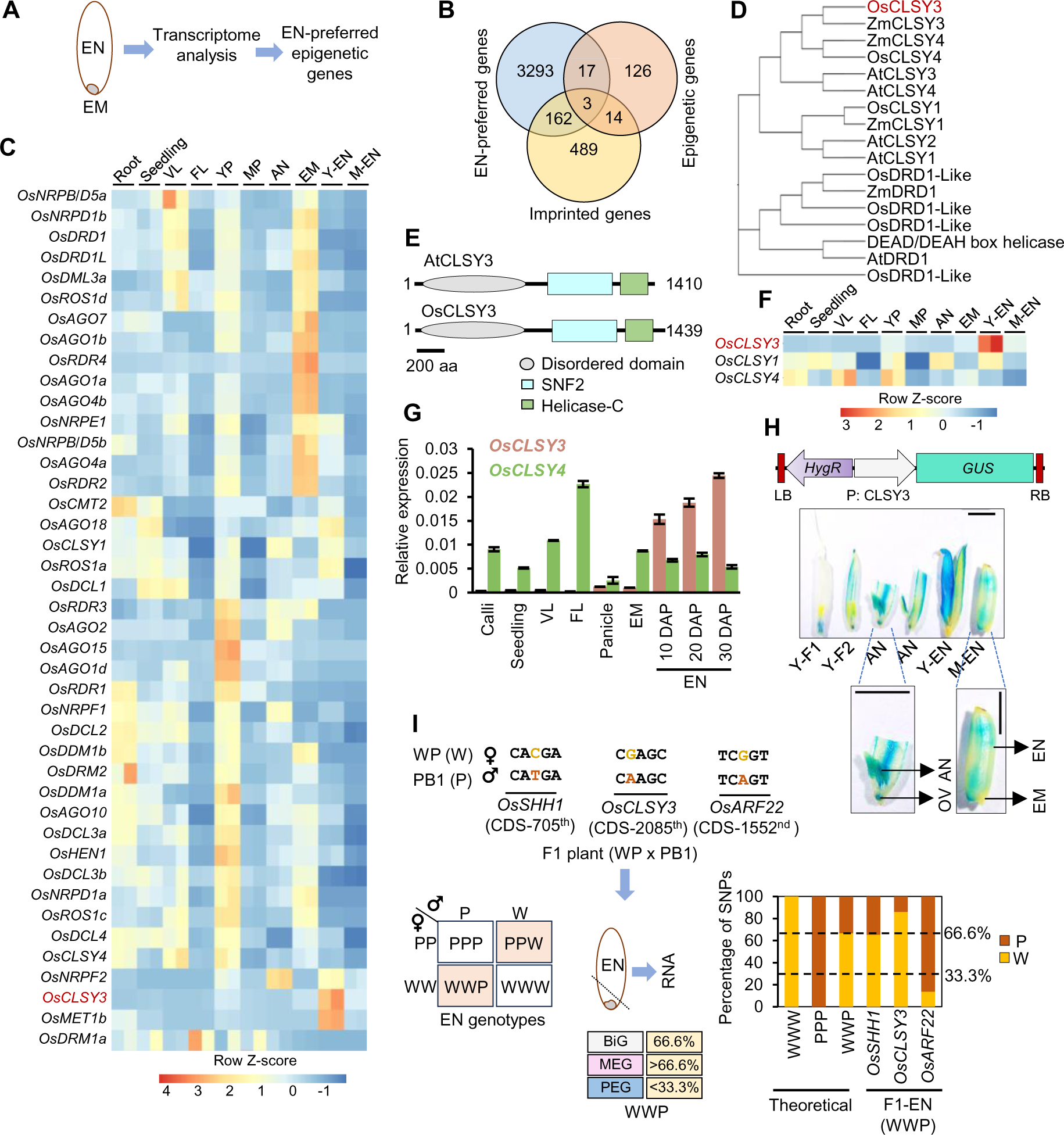
*OsCLSY3* is an endosperm-preferred imprinted gene. (A) Schematic showing EM (embryo) and EN (endosperm) tissues taken for transcriptome analysis. (B) Venn diagram showing overlap between epigenetic genes, EN-preferred genes and all known imprinted genes. (C) Heatmap depicting tissue-specific expression of epigenetic genes. Root (GSE166669), seedling (GSE229604), VL-vegetative leaf (GSE229604), FL-flag leaf (GSE111472), YP-young panicle (GSE180457), MP-mature panicle (GSE107903), AN-anther (GSE180457), Y-EN-young EN (15 days after pollination-DAP), M-EN-mature EN (25 DAP) (GSEXXXXXX). Row Z-score was plotted. (D) Phylogenetic tree for DRD1 family proteins in *Arabidopsis*, rice and maize. 1000 bootstrap replications. (E) Domain architecture of CLSY3 proteins (predicted). (F) Expression of rice CLSYs across tissues (RNA-seq). (G) RT-qPCR analysis of *OsCLSY3* and *OsCLSY4* across different tissues. *OsActin* served as internal control. Error bar-Standard Error (SE). (H) Vector map of P:CLSY3:GUS, and GUS expression pattern across reproductive tissues. Y-F1-spikelet before anthesis, Y-F2-spikelet 1 d before pollination, OV-unfertilized ovule. Scale bar (SB)-1 mm. (I) Scheme depicting WP and PB1 cross, and the SNPs identified in CDS of *OsARF22* (PEG), *OsCLSY3* (putative MEG) and *OsSHH1* (BiG). Punnett square showing possible EN genotypes. Transcript contribution from the WWP genotype seeds is shown for imprinted genes. Stacked barplots showing theoretical and observed transcript contribution of three genes in WWP EN.

We explored if the differential expression of genes were due to epigenetic players, chromatin modifiers, transcription factors and associated co-factors. To find possible endosperm-preferred epigenetic-regulators, we compared expression of 160 epigenetic genes in endosperm (Chen and Zhou, 2013; Shi et al., 2014; Higo et al., 2020; Wang et al., 2022; Deng et al., 2016). Among these, 20 were highly expressed in endosperm (Fig. 1B). We observed that candidates such as *SDG714, OsAGO12, OsMORC6, ENL-1 like* chromatin remodelers, several methyltransferases, JMJ genes and previously studied PcG complex related genes showed endosperm-preferred expression (Supplemental dataset S1 and Supplemental dataset S2). Many epigenetic genes themselves were imprinted and they can regulate endosperm development and imprinting of other genes (Luo et al., 1999; Kiyosue et al., 1999; Köhler et al., 2005; Mosher et al., 2009; Zhang et al., 2012; Dhatt et al., 2021; Cheng et al., 2021). To find endosperm-preferred imprinted epigenetic genes, we overlapped endosperm-preferred genes with 668 known imprinted genes (Luo et al., 2011; Yuan et al., 2017; Chen et al., 2018; Rodrigues et al., 2021) and 160 epigenetic genes (Supplemental dataset S3). We found 3 epigenetic genes, i.e., *OsFIE1*, *OsJMJ706* and an unannotated gene Os02g0650800 that were imprinted as well as highly expressed in rice endosperm (Fig. 1B). The roles of *OsFIE1* in endosperm development (Luo et al., 2009; Zhang et al., 2012; Huang et al., 2016; Dhatt et al., 2021; Cheng et al., 2020), and *OsJMJ706* in flower development are well known (Sun and Zhou, 2008; Liu et al., 2014). However, identity and function of Os02g0650800, which is a majorly endosperm-preferred gene that is listed as an imprinted gene in two published datasets was unknown (Fig. 1C). The gene was grouped in DRD1 family of SNF2 domain containing chromatin remodeler in rice (CHR740) (Hu et al., 2013b). A phylogenetic analysis indicated that CHR740 was close to *AtCLSY3* (Fig. 1D) and matched previous phylogenetic studies (Yang et al., 2018a) (Supplemental Table S4). The CHR740 had 29.3% amino acid identity, 44.5% amino acid similarity with AtCLSY3 and both having N-terminal Intrinsically disordered region (IDR), C-terminal SNF2 domains as predicted in InterProScan and Prosite tools (Quevillon et al., 2005; Sigrist et al., 2002) (Fig. 1E). IDRs are crucial for many chromatin associated epigenetic regulators (Musselman and Kutateladze, 2021). Monocots such as rice and maize seem to have three *CLSY* genes unlike four *CLSY*s found in *Arabidopsis* (Fig. 1D) (Hale et al., 2007; Hu et al., 2013b; Yang et al., 2018a). We found that maize ortholog of *OsCLSY3* GRMZM2G178435, is also a MEG (Waters et al., 2013) and it is closely related to OsCLSY3 (45.3% amino acid identity and 57.5% amino acid similarity). We noted that rice CHR722 (Os07g0692600) and CHR742 (Os05g0392400) were similar to *AtCLSY1* and *AtCLSY4*, respectively (Fig.1D). The CHR722 was already denoted as *OsCLSY1*, where it was implicated in anaerobic germination and seedling growth (Castano-Duque et al., 2021). In *Arabidopsis*, *CLSYs* have tissue-specific expression, while *AtCLSY1*, *AtCLSY2* are majorly expressed in leaf and flower buds, whereas *AtCLSY3*, *AtCLSY4* are majorly expressed in unfertilized ovules (Zhou et al., 2022). We observed that *OsCLSY3* and *OsCLSY1* are majorly expressed in endosperm and embryos, respectively. However, unlike *Arabidopsis*, *OsCLSY4* is expressed ubiquitously (Fig.1F, Supplemental Fig. S2A and B). The expression patterns of *CLSY* genes in maize also matched the same pattern as in rice, indicating that monocot *CLSYs* have similar expression patterns (Supplemental Fig. S2C). RT-qPCR analysis across different rice tissues further confirmed spatiotemporal expression of *CLSYs* (Fig.1G and Supplemental Fig. S2D). In order to further clarify the tissue-specific expression of *OsCLSY3*, we generated a β-glucuronidase (GUS) reporter construct driven by 1.1 kb promoter of *OsCLSY3* (Supplemental Fig. S2E). We observed that its expression was restricted in the unfertilized ovule, specific-anther stages and all tissues within the endosperm, but not in embryo (Fig.1H).

To investigate imprinting status of *OsCLSY3* further, we identified single nucleotide polymorphisms (SNPs) in the *OsCLSY3* CDS of different rice varieties. We performed crossing with *indica* rice variety Whiteponni (WP) as maternal and PB1 as paternal parents. Endosperm of F1 plants were genotyped and imprinting status of *OsCLSY3* (10 DAP) was tested (Supplemental Fig. S2F). We used *OsSHH1* as a biallelic gene (BiG) and a published paternally expressed gene (PEG) *OsARF22* as controls (Chen et al., 2018). We observed that for *OsCLSY3*, 86.1% of transcripts came from the maternal genome when compared to *OsSHH1* (BiG) which showed 65.3%. As expected in the case for *OsARF22* (a PEG), 86.4% of transcripts were from the paternal genome (Fig.1I, Supplemental Fig. S2F). These analyses demonstrated that *OsCLSY3* is a maternally expressed imprinted (MEG) in rice.

### DNA methylation at MITE TEs control expression of *OsCLSY3*

Since MEGs in rice are majorly regulated by DNA methylation at their proximal regions, we probed DNA methylation status of *OsCLSY3* promoter to understand the basis for its endosperm-preferred expression. Surprisingly, two heavily methylated Miniature Inverted TE repeats (MITE)-like regions, one similar to Ditto element, and another to Tourist element, were observed 600 bp upstream of *OsCLSY3* transcription start site. To investigate the role of methylation in *OsCLSY3* regulation, PB1 seeds were germinated in MS media and transferred to DNA methyltransferase blocker 5-Aza-2′-Deoxycytidine (AZA) containing media, as described (Chen et al., 2018) (Fig. 2A). We observed ectopic expression of *OsFIE1*, a well-known endosperm-specific MEG as well as *OsCLSY3* in a dose-dependent manner (Fig. 2B). Ubiquitously expressed *OsCLSY4* was found unaltered upon AZA treatment, indicating that DNA methylation regulated expression of *OsCLSY3* (Fig. 2B). In leaf, the *OsCLSY3* promoter (2 kb) was heavily methylated majorly at CHH sites when compared to the *OsCLSY4* promoter (Fig. 2C). As TEs and repeats at the gene promoter can regulate tissue-specific DNA methylation (Kinoshita et al., 2004; Satyaki and Gehring, 2017), we measured DNA methylation level of those MITEs by targeted bisulfite (BS)-PCR. DNA methylation at MITE TEs was higher in vegetative leaf and young panicle when compared to endosperm (Fig. 2D). This result was in agreement with published embryo and endosperm DNA methylation datasets (Rodrigues et al., 2021), specific to *OsCLSY3* promoter and not *OsCLSY4* promoter (Fig. 2E). These results supported the antagonistic relation between DNA methylation at *OsCLSY3* promoter and its expression across tissues.

**Figure 2.**
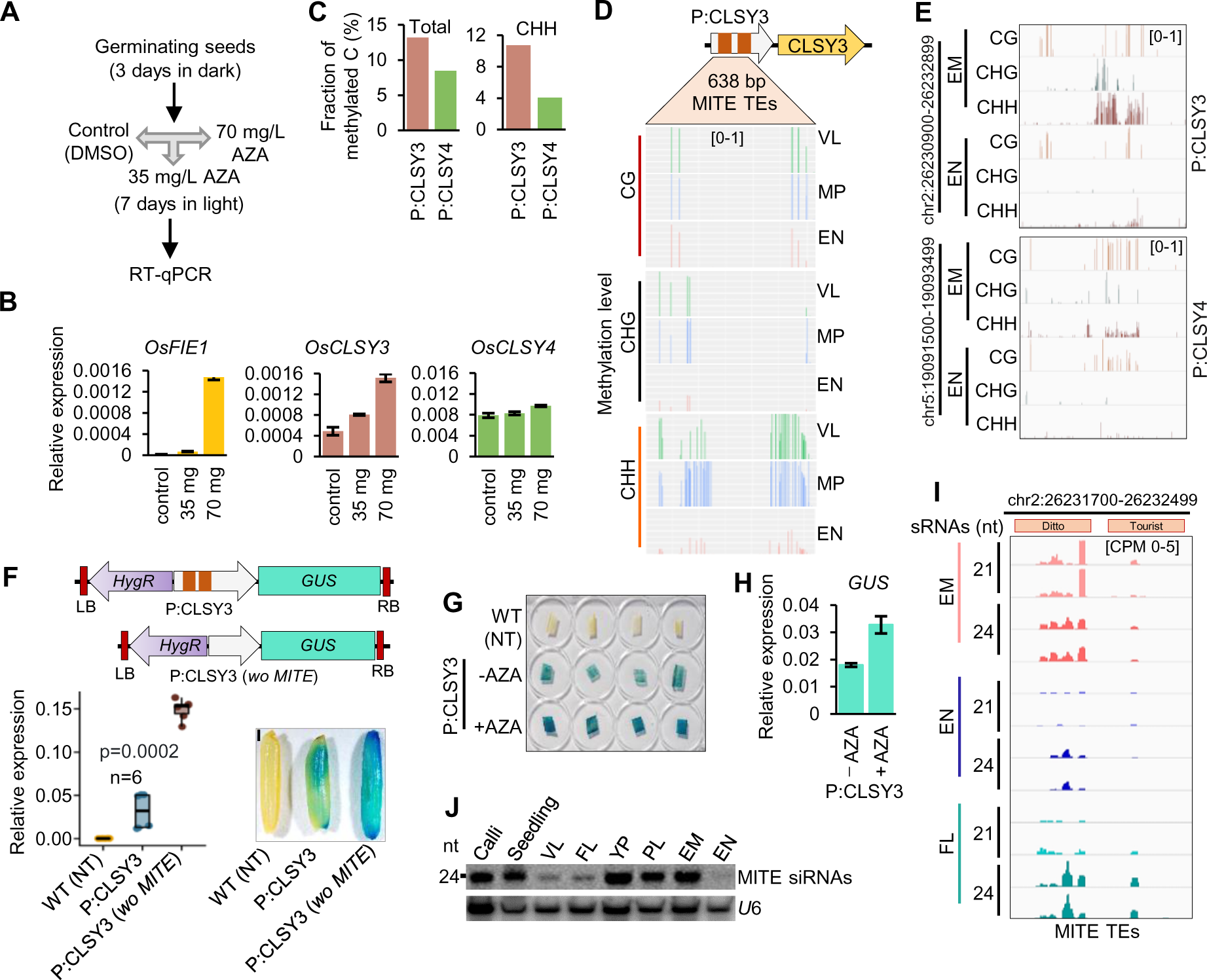
DNA methylation at MITE TEs controls expression of *OsCLSY3*. (A) Scheme used for AZA treatment. (B) RT-qPCR showing relative expression of *OsCLSY*s and *OsFIE1*, normalised to *OsActin*. Error bar-SE. (C) Barplots showing total (left) and only CHH (right) DNA methylation status of 2 kb promoters of *OsCLSY*s in leaf (GSE130168). (D) Analysis of DNA methylation level of MITE TEs by targeted BS-PCR sequencing across tissues. (E) IGV screenshots showing DNA methylation status of promoters of *OsCLSY*s in EM and EN (GSE130122). (F) Linear map of GUS reporter constructs with 1.1 kb native *OsCLSY3* promoter or 600 bp deletion of MITE TEs in its promoter (*wo MITE*). NT-non-transgenic. Boxplots showing relative expression of *GUS* in transgenic panicles (two-tailed Student’s *t*-test). Mature seeds showing levels of GUS (SB-1 mm). Same P:CLSY3 seed was also used in Fig.1H. (G) Image showing GUS expression in 10 d-old AZA treated seedlings. (H) Barplot showing *GUS* level upon AZA treatment. (I) IGV screenshots showing MITE derived 21 nt and 24 nt sRNAs across different tissues. (J) Northern blot showing MITE derived sRNA levels across tissues. *U*6 served as loading control (PL-palea-lemma).

Further to study the role of MITE TEs in context of transcriptional regulation of *OsCLSY3*, we generated transgenic GUS reporter plants, with GUS gene driven by wild type (WT) *OsCLSY3* promoter (1.1 kb) (Fig. 1H) or by *OsCLSY3* promoter with deleted MITE TEs. Removal of TEs significantly elevated GUS expression which further suggested that DNA methylation repressed *OsCLSY3* expression *via* MITE TEs (Fig. 2F). Further, the GUS expression was increased upon AZA treatment, supporting our previous observations (Fig. 2G and H). Short TEs near to gene are regulated by the RdDM pathway when cytosine methylation marks are majorly at CHH contexts (Matzke et al., 2015). Since MITEs were methylated densely at CHH sites, we quantified the sRNA levels across different tissues from previously published datasets (Hari Sundar G et al., 2023; Swetha et al., 2018). We found that 24 nt sRNAs at the MITEs TEs were more abundant in embryo and flag leaf when compared to endosperm in our sRNA datasets (Fig. 2I) and in northern analysis (Fig. 2J). The sRNA level variation across tissues also suggests a clear correlation between sRNAs and DNA methylation at the *OsCLSY3* promoter.

### DNA methylation at MITE TEs is regulated through RdDM

If the RdDM pathway regulated DNA methylation of MITE TEs, signatures of perturbation of this regulation must be seen across mutants in the RdDM pathway. Abundance of MITE derived sRNAs in poliv-kd panicle (Hari Sundar G et al., 2023) were drastically reduced in comparison to WT (Fig. 3A and B). Since sRNAs that establish DNA methylation are loaded into AGO4 in *Arabidopsis* (Zilberman et al., 2003), we investigated available rice AGO RNA Immunoprecipitation (RIP) datasets (Wu et al., 2009) and found that MITE derived sRNAs were enriched in OsAGO4a and OsAGO4b (Fig. 3C). Correspondingly, *OsCLSY3* mRNA expression was high in poliv-kd young panicle (Fig. 3D). We also measured sRNA and DNA methylation levels using published datasets of several RdDM mutants (Wang et al., 2022; Hu et al., 2022; Chakraborty et al., 2022). We found that RdDM mutants such as *rdr2* and *nrp(d/e)2* show reduction of MITE derived sRNAs in the *OsCLSY3* promoter, whereas, as expected, MITE derived sRNAs were unaltered in *polv* (Fig. 3E). Since *Arabidopsis* CLSYs recruit PolIV at specific TEs and repeats (Zhou et al., 2018), we hypothesized that MITE TEs might be regulated by another member of the CLSY family. *OsCLSY4* was found majorly expressed in vegetative tissues unlike *OsCLSY3* (Fig. 1F and G). We hypothesized that OsCLSY4 might be recruiting PolIV to the MITE regions in the *OsCLSY3* promoter in vegetative tissues. To check this, we generated clsy4-kd plants by artificial miRNA (amiR) strategy (Ossowski et al., 2008; Warthmann et al., 2008; Narjala et al., 2020) and compared silencing of MITEs through sRNAs. In clsy4-kd leaf, we observed dose-dependent reduction of MITE sRNAs (Fig. 3F). Also, we observed drastic reduction in MITE derived sRNAs in clsy4-kd similar to poliv-kd panicle, indicating OsCLSY4 is likely recruiting PolIV into *OsCLSY3* promoter (Fig. 3F). Using bisulfite sequencing (BS-PCR), we found a reduction of DNA methylation at the *OsCLSY3* promoter in leaves of clsy4-kd, which indicated that OsCLSY4 controls expression of *OsCLSY3 via* RdDM (Fig. 3F). We also quantified DNA methylation levels in the MITE region in several known RdDM mutant datasets (Zheng et al., 2021; Hu et al., 2022). We observed, similar to sRNAs, methylation was also reduced in *rdr2* mutant (Fig. 3G). Among datasets derived from several DNA methyltransferase mutants (Hu et al., 2022), only in *drm2* and its other combinations, a total abolishment of DNA methylation at *OsCLSY3* promoter was observed (Fig. 3H). All these results conclusively demonstrate that the RdDM pathway regulates expression of *OsCLSY3*.

**Figure 3.**
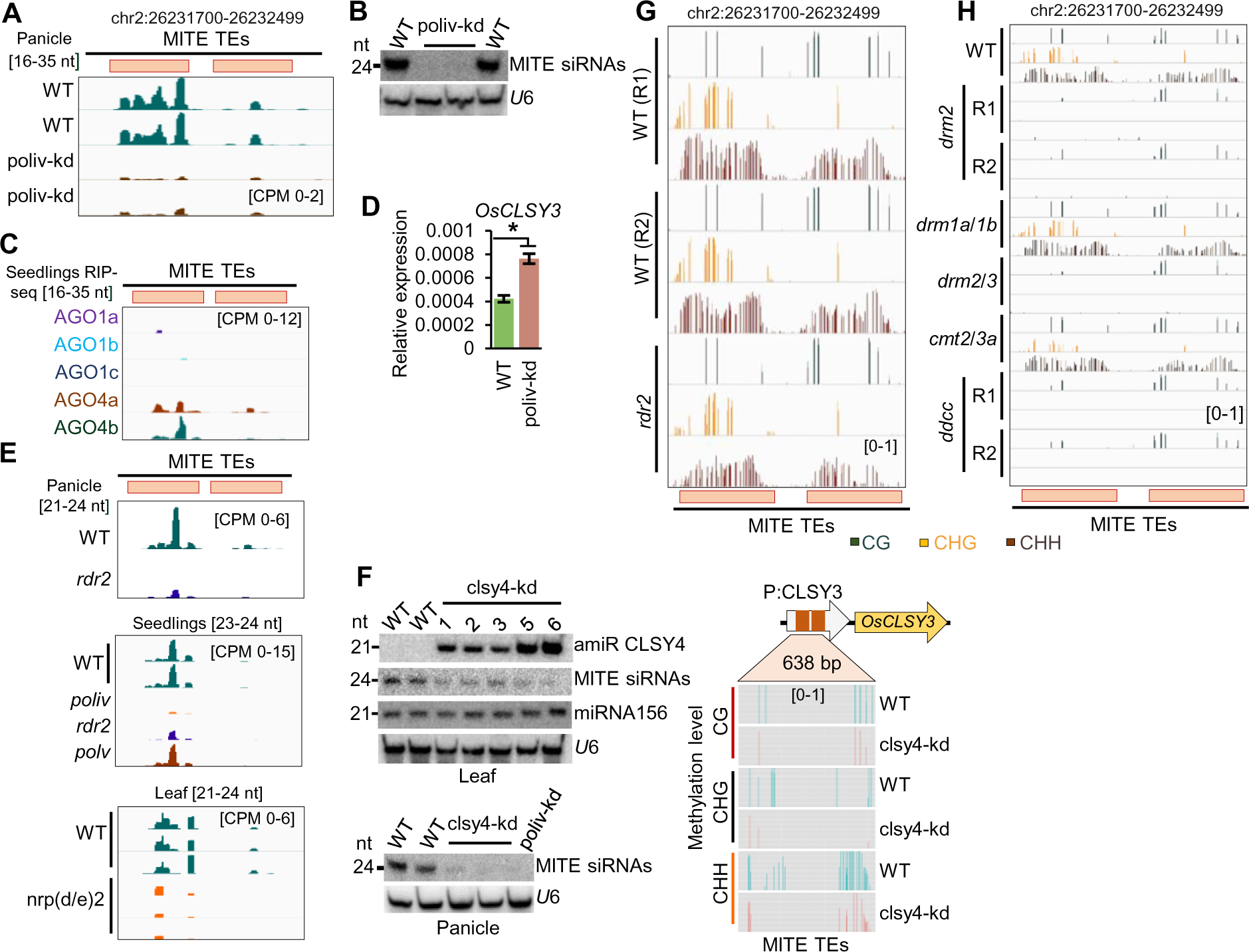
DNA methylation in *OsCLSY3* promoter is regulated by the RdDM. (A) IGV screenshots showing *OsCLSY3* promoter derived sRNA levels (16-35 nt) in poliv-kd panicle. (B) Northern blot showing sRNA levels in poliv-kd panicle. (C) IGV screenshots showing enrichment of *OsCLSY3* promoter derived sRNAs (16-35 nt) in RIP of different AGOs in rice (GSE18250 and GSE20748). (D) Barplot showing *OsCLSY3* level in poliv-kd panicles. *OsActin* served as internal control. Error bar-SE. (E) IGV screenshots showing status of MITE derived sRNAs among RdDM mutants of rice (GSE130166, GSE158709, PRJNA758109). (F) Northern blots showing MITE derived sRNA level in clsy4-kd and poliv-kd (left panels) and analysis of DNA methylation in *OsCLSY3* promoter in clsy4-kd leaf by BS-PCR (right panel)*. U*6 served as loading control. (G) IGV screenshots showing methylation status of 800 bp MITE TE region in *OsCLSY3* promoter in *rdr2* seedlings (GSE130168). (H) IGV screenshot showing methylation status in leaves of *ddccc* mutant (GSE138705). *ddccc-drm2/drm1/cmt2a/cmt2b/cmt3*.

### *OsCLSY3* knockout (*KO)* plants exhibited sterility

While *OsCLSY3* is an endosperm-preferred imprinted gene in monocots such as rice and maize (Luo et al., 2011; Chen et al., 2018; Waters et al., 2013), in *Arabidopsis* its homolog is not an imprinted gene (Hsieh et al., 2011; Wolff et al., 2015). However, the RdDM pathway derived sRNAs regulate endosperm development in *Arabidopsis* (Kirkbride et al., 2019). Since the role of *CLSY3* in the context of rice endosperm development is not explored, to understand that we targeted *OsCLSY3* with CRISPR-Cas9 gRNA against a unique region of 1^st^ exon of *OsCLSY3* and generated *KO* plants. Presence of intact T-DNA was confirmed by junction-fragment Southern blot analysis (Supplemental Fig. S3A). In all these plants, the *OsCLSY3* gene was edited as desired (Supplemental Fig. S3B). Vegetative growth of *KO* plants was unaffected (Supplemental Fig. S3C), however, three mutants (*KO* #1, #4, #5) showed strong phenotypes in panicles including complete sterility, indicating that *OsCLSY3* plays an essential role in reproduction (Supplemental Fig. S3D). Rest of the genome edited mutants where defects were not observed had either 1 or 2 amino acids deletions without altering the protein significantly. The *poliv* mutant of rice and *Capsella* showed very low yield due to drastic pollen defects (Hari Sundar G et al., 2023; Wang et al., 2020). In order to check if *KO* plants show pollen sterility, we performed pollen viability assay and found non-viable pollen grains in *KO* plants (Supplemental Fig. S3E). A drastic reduction in number and morphologically defective, prematurely dead pollens were observed in *KO* lines (Supplemental Fig. S3F and G). We found few endosperms in *KO,* but they were either dead or severely defected (Supplemental Fig. S3H). However, none of the *KO* plants displayed growth defects in vegetative stages, indicating that the role of *OsCLSY3* is restricted to specific tissues such as pollens and endosperm (Supplemental Fig. S3I). All these results collectively suggest that *OsCLSY3* is also crucial for overall reproductive development.

### Knockdown (kd) and overexpression of *OsCLSY3* (OE) transgenic plants displayed yield-related phenotypes

Since *KO* plants were sterile, we planned to knockdown the gene through amiRs strategy (Ossowski et al., 2008; Warthmann et al., 2008; Narjala et al., 2020). We generated transgenic plants with two amiRs driven by constitutive promoters in one T-DNA which was named as clsy3-kd2 (double amiR) (Supplemental Fig. S4A and B). To silence *OsCLSY3* in a dose-dependent manner, we also generated transgenics expressing single amiR (Supplemental Fig. S4A and B). Integration of T-DNA in the genome was verified by Southern blot (Supplemental Fig. S4C). We obtained a total of 7 transgenic plants with double amiRs (clsy3-kd2) (Fig. 4A, Supplemental Fig. S4D) and 8 plants with single amiR (clsy3-kd1) (Fig. 4A). We selected two different clsy3-kd lines for phenotyping and qRT PCR (clsy3-kd1 plant #1 was denoted as clsy3-kd1, and clsy3-kd2 plant #5 as clsy3-kd2, respectively). About 76% and 60% reduction in Os*CLSY3* transcripts were observed in clsy3-kd2 and clsy3-kd1 endosperm tissues, respectively (Fig. 4B). Both clsy3-kd lines did not show abnormalities in vegetative tissues (Fig. 4C and E), but they had smaller seeds when compared to WT (Fig. 4D). The clsy3-kd had reduced seed length, seed width and weight, indicating a crucial role of *OsCLSY3* in endosperm development (Fig. 4F). We obtained one transgenic line (clsy3-kd2 #3) in which T-DNA was intact but amiR was completely silenced (Supplemental Fig. S4C and D). We observed that seeds of this line were similar to WT (Supplemental Fig. S4E). The clsy3-kd plants displayed minor alterations in primary root length in the second generation, probably due to the change in germination time (Supplemental Fig. S4F). To investigate the tissue-specific role of *OsCLSY3* further, we generated OE plants driven by a constitutive promoter (Supplemental Fig. S4H). The single copy OE plants (Fig. 4G), showed robust vegetative growth and had bigger seeds when compared to WT (Fig. 4H and I). We observed that grain length and grain width were also increased in OE plants (Fig. 4J-L). However, the grain filling rate was significantly reduced probably due to trade-off phenotype (Fig. 4M). To study the internal morphology of clsy3-kd and OE seeds, we sectioned endosperm and observed chalkiness, a signature of altered cellularization, in OE endosperms (Supplemental Fig. S4I). The chalkiness is considered as a poor agronomic quality of the grains (Wada et al., 2019; An et al., 2020). In OE seedlings, we observed smaller roots which was exactly the opposite of that we observed in clsy3-kd seedlings (Supplemental Fig. S4J and F). These phenotypes were in agreement with perturbed endosperm development, since this tissue plays important roles in germination and seedling development (Yan et al., 2014; Iwasaki et al., 2019). Our germination assay revealed slower seed germination in OE when compared to WT and clsy3-kd (Supplemental Fig. S4K). These results collectively suggest that proper spatiotemporal expression of *OsCLSY3* is crucial for normal development in rice.

**Figure 4.**
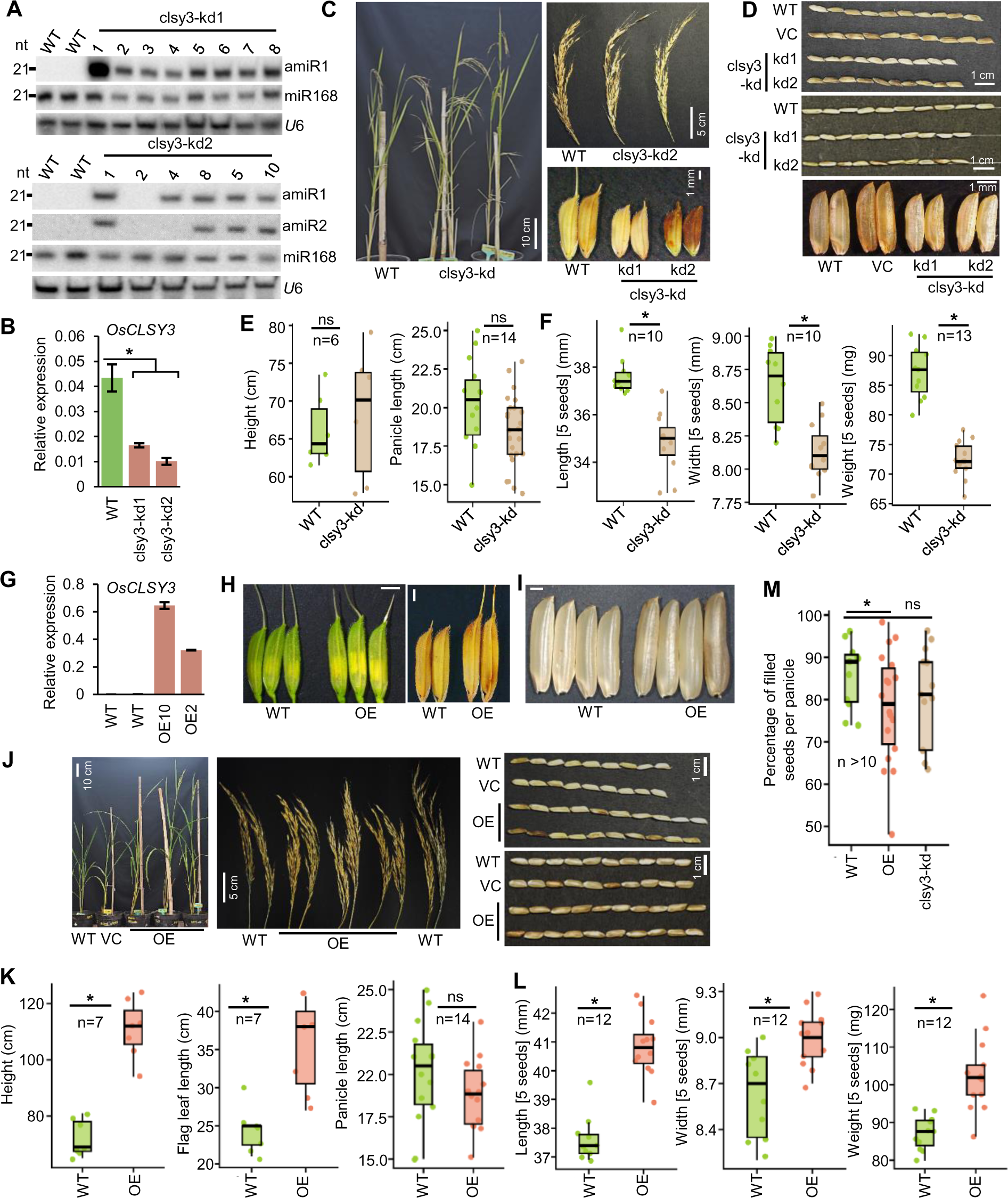
Phenotypes of clsy3-kd and OE lines. (A) Northern blots showing expression of amiRs. (B) Barplot showing levels of *OsCLSY3* expression in clsy3-kd EN. *OsActin* served as internal control. Error bar-SE. *-significant. Two-tailed Student’s *t*-test. (C) Images showing morphology of clsy3-kd plants, panicles and seeds when compared to equally grown WT. (D) Seed morphology of clsy3-kd with and without de-husking. (E) Boxplots showing height and panicle length of clsy3-kd plants. (F) Boxplots showing 5 seeds length, width and weight in clsy3-kd. (G) Barplot showing expression of *OsCLSY3* in OE plant leaves. *OsActin* served as internal control. Error bar-SE. (H) Images showing spikelet and seed morphology in OE. SB-1 mm. (I) Images showing de-husked OE seeds SB-1 mm. (J) Images showing morphology of OE plants, panicles and seeds (with and without de-husking) compared to WT. (K) Boxplots showing height, flag leaf and panicle length of OE plants. (L) Boxplots showing 5 seeds length, width and weight in OE. (M) Boxplot showing percentage of seed filling in OE and clsy3-kd plants. In box plots, *-significant, ns-non significant (two-tailed Student’s *t*-test).

### Nearly 70% of all endosperm-specific sRNAs were *OsCLSY3* dependent

CLSYs in *Arabidopsis* recruit PolIV into repeats and transposons, hallmark of which include production of 24 nt sRNAs derived from these regions (Zhou et al., 2018; Yang et al., 2018a). To identify endosperm-specific sRNA loci, sRNA sequencing in clsy3-kd was carried out (clsy3-kd2). Nearly 90% of an average of 20 million reads mapped to rice genome (Supplemental Table S1). The 23-24 nt sRNA profile was different in endosperm when compared to other tissues as observed previously (Rodrigues et al., 2013; Grover et al., 2020). Unlike flag leaf and embryo tissues, fewer sRNA loci contributed to the 23-24 nt sRNA pool in endosperm (Fig. 5A). A clear reduction in different size classes of sRNAs was observed in clsy3-kd endosperm (Fig. 5B and Supplemental Fig. S5A). Principal component Analysis (PCA) indicated that the identical pool of 23-24 nt endosperm-specific sRNAs were downregulated in clsy3-kd and poliv-kd endosperm tissues (Fig. 5C). Reduction in 23-24 nt sRNAs were observed chromosome-wide (Fig. 5D). These sRNAs were derived from genic regions as well as from, class I and class II TEs (Fig. 5E). Majority of the 23-24 nt sRNAs which had 5’A were drastically reduced in clsy3-kd (Supplemental Fig. S5B). To identify CLSY3-dependent sRNA loci, we quantified number of loci present in WT and clsy3-kd endosperm by Shortstack analysis (Supplemental dataset S3 and S4). We observed around 70% of sRNA loci lost sRNAs in clsy3-kd compared to WT which were further called CLSY3-dependent sRNA loci (Fig. 5F and Supplemental dataset S6). Since clsy3-kd plants displayed a global reduction of sRNAs, we checked pools of 21-22 nt sRNAs mapped to miRNA encoding loci (Griffiths-Jones et al., 2008; Anushree and Shivaprasad, 2017), and they were not significantly reduced in clsy3-kd (Fig. 5G). The 23-24 nt CLSY3-dependent sRNAs were derived from both class I (Gypsy, LINE1, SINE, LTR) and class II (MITE, En-Spm) TEs (Fig. 5H and Supplemental Fig. S5C). Since CLSY-dependent sRNAs can induce DNA methylation, we quantified DNA methylation of some TEs (MITE, 5S-rDNA repeats) which were generally hypermethylated in endosperm and overlapped with CLSY3-dependent sRNA loci. We found a clear reduction in DNA methylation at MITE and 5S-rDNA repeats, correlating with their reduced sRNA levels (Fig. 5I). As a control, we quantified DNA methylation in a few TEs (LINE1, CACTA), which were hypomethylated in the endosperm and did not overlap with CLSY3-dependent sRNA loci. We found unaltered DNA methylation and sRNAs at those TEs in clsy3-kd lines (Fig. 5I, Supplemental Fig. S5D and E). These results collectively suggest that *OsCLSY3* is a major regulator of sRNAs in endosperm and CLSY3-dependent sRNAs are crucial for silencing of specific classes of TEs and repeats through RdDM.

**Figure 5.**
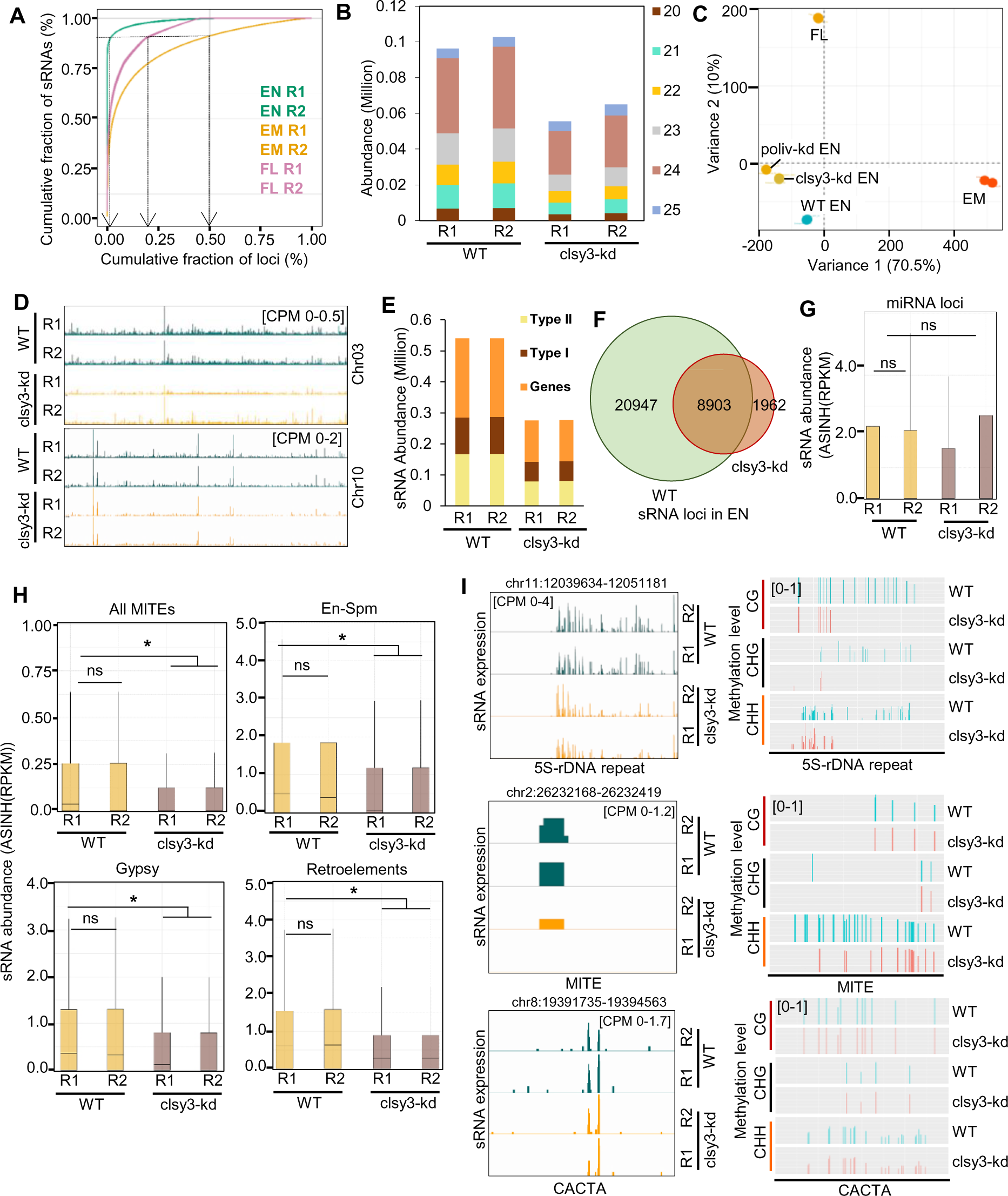
*OsCLSY3* regulates repeat and TE-derived sRNAs and DNA methylation in EN tissue. (A) Percentage cumulative sum plots for sRNAs (23-24 nt) in rice tissues. The arrows indicate cumulative percentage of loci which generates 90% of all sRNAs. (B) Stacked barplot showing abundance of mapped sRNAs (20-25 nt) in WT and clsy3-kd EN. (C) PCA plot comparing the mapped sRNAs (23-24 nt) across different tissues and genotypes. poliv-kd EN (16-20 DAP) dataset is GSE180456. (D) IGV screenshots showing levels of sRNAs (23-24 nt) across different chromosomes. (E) Stacked barplot showing sRNAs (23-24 nt) across class I and class II TEs. (F) Venn diagram showing sRNA loci across WT and clsy3-kd EN. (G) Boxplot showing expression of miRNA transcripts in WT and clsy3-kd EN. *-significant, ns-non-significant (Wilcoxon test p<0.01). (H) Boxplots showing expression of different sRNAs across class I and class II TEs in WT and clsy3-kd EN. (I) IGV screenshots showing sRNAs abundance in 5S-rDNA repeat, MITE, CACTA TEs (left side) and targeted BS-PCRs showing methylation status of those TEs (right side). Replicates (R1 and R2) are identically coloured.

### *OsCLSY3* regulates expression of endosperm-specific siren loci and adjoining genes in endosperm

Since in *Arabidopsis* ovules, *CLSY3* and *CLSY4* generate sRNAs from siren loci (Zhou et al., 2022; Martins and Law, 2023), we investigated if *OsCLSY3* is involved any role in the regulation of siren loci (Supplemental dataset S7). We measured 23-24 nt sRNAs derived from known rice siren loci (Rodrigues et al., 2021), and found sRNAs from these loci were reduced in clsy3-kd and poliv-kd endosperm tissues (Fig. 6A-C, Supplemental Fig. S6A and B). As observed previously (Rodrigues et al., 2021, 2013), siren loci were less expressed in embryo and flag leaf (Fig. 6D-E and Supplemental Fig. S6B). In a northern blot analysis, we found siren sRNAs were reduced in clsy3-kd and poliv-kd endosperm tissues, but not in clsy4-kd endosperm, indicating a direct role of *OsCLSY3* (Fig. 6F). Since siren loci derived sRNAs have the potential to guide DNA methylation at *cis* or *trans* loci (Rodrigues et al., 2013; Grover et al., 2020; Long et al., 2021; Burgess et al., 2022), we performed a targeted BS-PCR and found a clear reduction in DNA methylation (Fig. 6G). The observed reduction of methylation was comparatively more drastic in poliv-kd than in clsy3-kd endosperm, probably due to compensatory effect, likely involving other *CLSYs* (Fig. 6G). These results collectively suggested that *OsCLSY3* also regulated sRNAs production from the siren loci and their regulation of DNA methylation. Although endosperm-development related phenotypes were not reported in *Arabidopsis* clsy mutants, we found smaller endosperm in clsy3-kd seeds when compared to WT (Fig. 4D). To understand the mechanism that might be operating, a transcriptomic analysis was carried out in clsy3-kd endosperm (Supplemental Table S8 and S9). We shortlisted 2869 differentially expressed genes (DEGs) with log_2_ 1.5-fold upregulation or downregulation (Fig. 6H). To understand the correlation between levels of CLSY3-dependent sRNAs and DEGs, we compared 23-24 nt sRNA levels +/- 2 kb windows near DEGs and observed a reduction of sRNAs near the DEGs (Fig. 6I). A strong antagonistic correlation between sRNAs and gene expression was observed in around 1000 loci, indicating that CLSY3-dependent sRNAs might be regulating gene expression (Fig. 6J). Few genes which were upregulated in clsy3-kd, were hypermethylated in WT endosperm and the hypermethylated sites overlapped with CLSY3-dependent sRNA loci (Fig. 6K). This observation suggested that the genes were likely regulated by CLSY3*-*dependent DNA methylation, specific to endosperm. We also found mis-expression of well-known endosperm development and yield-related genes (Li et al., 2022, 2019) having siren loci adjacent to them (+/- 2 kb) in clsy3-kd lines (Fig. 6L). The adjacent genes next to CLSY3-dependent sRNA loci were mis-expressed in clsy3-kd (Supplemental Fig. S7A). Many MADS box genes previously identified as crucial regulators of endosperm development (Zhang et al., 2010; Paul et al., 2020; Chen et al., 2016; Zhang et al., 2018; Cheng et al., 2021) were mis-expressed in clsy3-kd (Supplemental Fig. S7A). As a control, we analyzed few genes that overlapped with CLSY3-independent sRNA loci. As expected, we observed these genes were unaltered in clsy3-kd (Supplemental Fig. S7B). In addition, many published seed-development related genes such *as OsMKKK10, OsFAD2, OsTAR1, OsNF-YB1* known for their role in seed development related to signaling and fat / carbohydrate metabolism were mis-expressed in clsy3-kd endosperm (Supplemental Fig. S7C). Among these few genes were overlapped with CLSY3-dependent sRNA loci (Supplemental Fig. S7C) indicating a direct role of *OsCLSY3* in their regulation. These results collectively demonstrate that CLSY3-dependent sRNAs direct endosperm development by regulating expression of multiple development, hormone and metabolism related genes.

**Figure 6.**
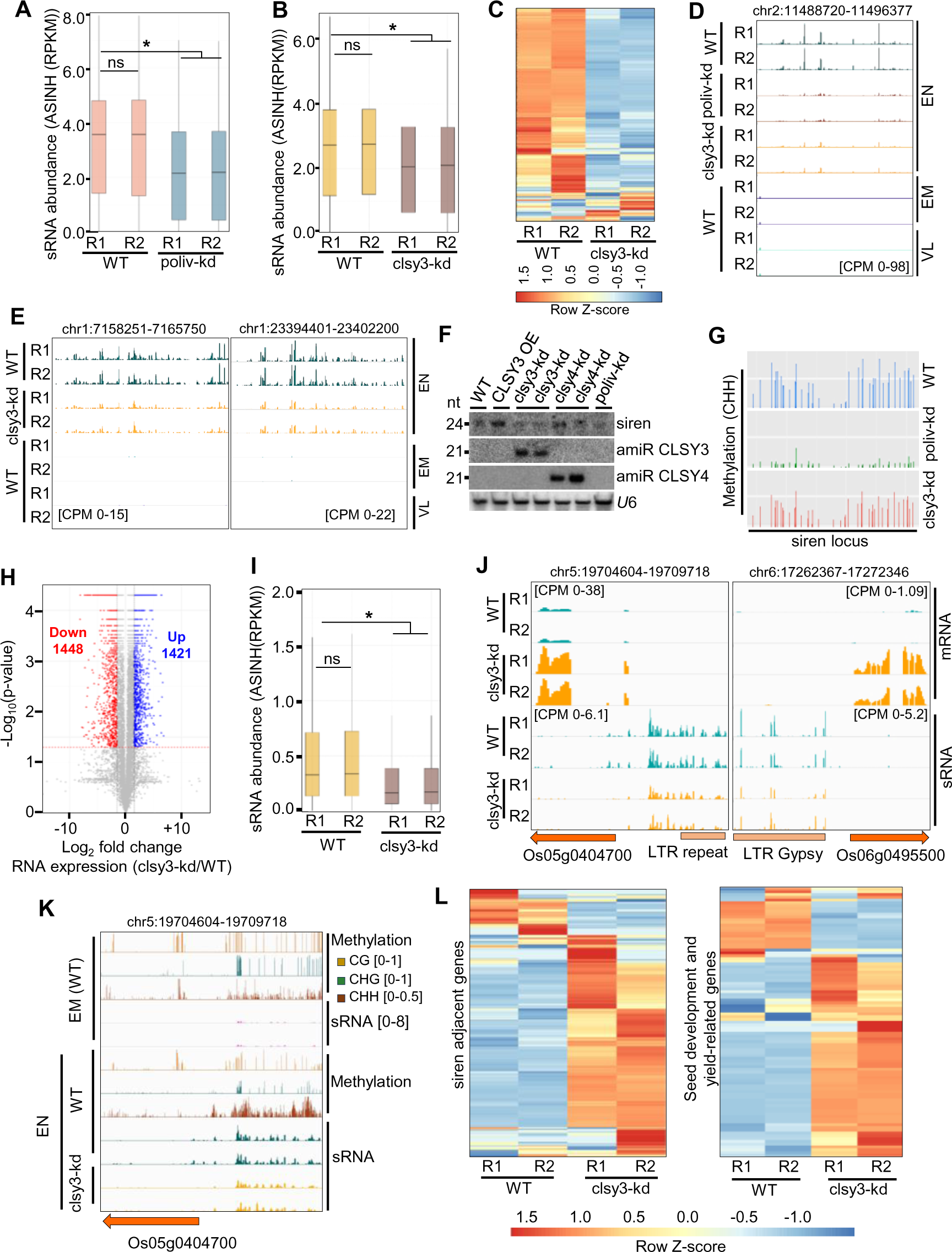
*OsCLSY3* controls expression of siren loci and development-related genes. (A) Boxplot showing sRNAs (23-24 nt) derived from siren loci in poliv-kd EN (797 loci). ASINH converted RPKM values were used for boxplot. (B) Boxplot representing status of sRNAs (23-24 nt) from siren loci in clsy3-kd EN. (C) Heatmap showing siren sRNAs in clsy3-kd EN. Row Z-score was plotted. (D) and (E) IGV screenshot showing expression of siren loci in clsy3-kd and poliv-kd EN. VL-vegetative leaf. (F) Northern blot showing siren sRNAs in RdDM mutants. (G) BS-PCR showing methylation status of a siren locus. (H) Volcano plot representing all DEGs of clsy3-kd. (I) Box plot showing sRNA levels near DEGs. (J) IGV screenshots showing expression of two selected DEGs and levels of adjacent sRNA loci in clsy3-kd EN. (K) IGV screenshots showing methylation and sRNA levels between EM and EN in a selected DEG. (L) Heatmap showing expression of published siren loci adjacent genes (510) and seed-development (102) genes. Row Z-score was plotted. In (B) and (I), *-significant and ns-non-significant (Wilcoxon test, p < 0.01).

### *OsCLSY3* regulates expression of imprinted genes through imprinted sRNA loci

It was observed that RdDM dependent sRNAs regulated expression of imprinted genes in *Arabidopsis* (Erdmann et al., 2017; Kirkbride et al., 2019), however, it was not known if *CLSYs* played any role in genomic imprinting. In the rice clsy3-kd transcriptome, we observed mis-expression of well-established imprinted genes, suggesting a possible role for *OsCLSY3* in imprinting (Fig. 7A and B). Quantification of total sRNAs derived from gene bodies and 5’ and 3’ regions of 2 kb of all imprinted genes showed a drastic reduction in clsy3-kd (Fig. 7C). In *Arabidopsis*, imprinted sRNAs regulated expression of proximal imprinted genes (Erdmann et al., 2017). In rice, imprinted sRNA loci were also found adjacent to many imprinted genes, and it was hypothesized previously that these sRNAs might be regulating imprinted genes (Rodrigues et al., 2013; Yuan et al., 2017; Chen et al., 2018; Rodrigues et al., 2021). We found among the 15 maternally expressed sRNA loci, around 9 loci were downregulated, whereas, all 16 paternally expressed sRNA loci were downregulated in clsy3-kd endosperm (Fig. 7D). As expected, described imprinted sRNA loci were also downregulated in poliv-kd similar to clsy3-kd (Supplemental Fig. S8A and B). However, most of the upregulated maternally expressed sRNA loci in clsy3-kd were downregulated in poliv-kd, probably due to a compensatory effect likely involving other *CLSYs* (Supplemental Fig. S8A and B). All the 20 imprinted genes identified in rice and which have antagonistic relationship with imprinted sRNAs showed upregulation in clsy3-kd lines (Fig. 7E and F). Methylation analysis revealed that imprinted sRNA loci were hypermethylated majorly in CHH context in WT endosperm when compared to WT embryo (Rodrigues et al., 2021) (Supplemental Fig. S9). Measurement of DNA methylation level by targeted BS-PCR indicated a reduction of methylation in clsy3-kd as well as in poliv-kd endosperm (Fig. 7G). Our further quantification of DNA methylation by quantitative chop-PCR showed decrease in methylation in clsy3-kd (Fig. 7H). In the experiment, *OsActin* and another locus did not show any significant change, indicating unaltered DNA methylation of these control loci in clsy3-kd (Fig. 7H). These results collectively suggested expression of many imprinted sRNAs loci and imprinted genes were regulated by *OsCLSY3*.

**Figure 7.**
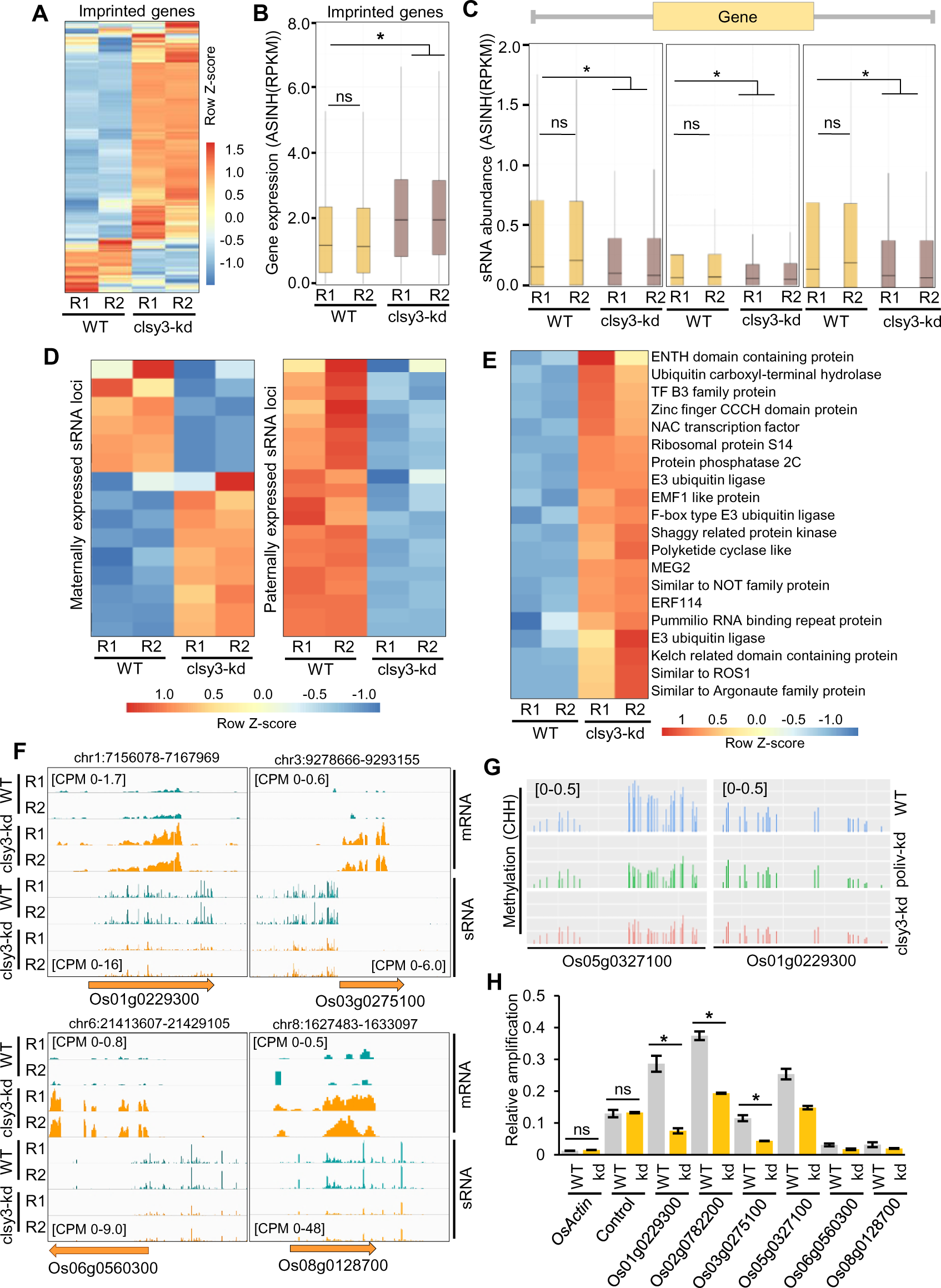
*OsCLSY3* controls expression of imprinted genes by regulating proximal imprinted sRNA loci. (A) Heatmap showing expression of all imprinted genes in clsy3-kd EN (668 genes). (B) Boxplot representing expression of imprinted genes in clsy3-kd EN. ASINH converted RPKM values were used for boxplot. (C) Boxplots showing abundance of CLSY3-dependent sRNAs in the promoter, terminator and gene body of imprinted genes. *-significant and ns-non-significant (Wilcoxon test, p < 0.01). (D) Heatmaps showing expression of sRNAs in imprinted sRNA loci. Row Z-score was plotted. (E) Heatmap representing expression of imprinted genes proximal to imprinted sRNA loci. (F) IGV screenshots showing expression of four imprinted genes in clsy3-kd EN. (G) BS-PCRs showing methylation status of two imprinted genes. (H) Barplot showing methylation status of six imprinted genes.

## Discussion

### *OsCLSY3* is essential for endosperm development in rice

Endosperm is unusual because it is a triploid tissue with more accessible chromatin where TEs and gene regulation is vastly different from other tissues including embryo (Li and Berger, 2012; Jiang and Köhler, 2012). Development of endosperm is unique and regulated by multiple genes that are further regulated by epigenetic pathways (Gehring, 2013; Köhler and Weinhofer-Molisch, 2010). Only in endosperm tissue, genomic imprinting, a key process for its development is observed. In agreement with all these, several DNA methylation and histone modification pathway players such as - *FIE, MEA, NRPD* in *Arabidopsis* and *OsFIE1, OsEMF2a, ZmFIE1* in monocots, were identified as crucial players for endosperm development and imprinting (Grossniklaus et al., 1998; Köhler et al., 2005; Luo et al., 2009; Zhang et al., 2012; Huang et al., 2016; Cheng et al., 2020; Tonosaki et al., 2021; Cheng et al., 2021). Interestingly, many of these genes were also found imprinted in endosperm where they were preferably expressed. Along with DNA methylation, DNA demethylation pathway players, such as demethylase DME in *Arabidopsis* and ROS1a in rice central cells, were also implicated in imprinting across plants (Xiao et al., 2003; Gehring et al., 2006; Kim et al., 2019). One of the crucial players in plant epigenetics are sRNAs and their expression drastically increases in reproductive tissues (Chow and Mosher, 2023; Hari Sundar G et al., 2023). Unlike *Arabidopsis*, other dicots like *B. rapa* and *Capsella* RdDM mutants showed pollen and seed abnormalities (Mosher et al., 2009; Grover et al., 2018; Wang et al., 2020). In rice too, RdDM pathway genes such as *PolIV, RDR2, PolV* regulate important agronomic traits such as panicle development, seed setting, pollen development etc. (Hari Sundar G et al., 2023; Wang et al., 2022; Higo et al., 2020; Zheng et al., 2021; Chakraborty et al., 2022). The guides for sRNA production in plants are a family of chromatin remodelers named *CLSYs* that regulate DNA methylation in a tissue- and locus-specific manner (Zhou et al., 2018, 2022). Using genetic, genomic and molecular approaches, we identified *OsCLSY3*, a previously unannotated, imprinted gene as a critical player in these processes. An ortholog of this gene is imprinted and majorly expressed in maize endosperm, indicating that monocot endosperms that are preserved during seed development, also have atypical regulatory layers. The strong phenotypes observed in the seeds of *OsCLSY3* mis-expression lines clearly indicate the importance of CLSY3 in seed development. (Fig.4D and H, Supplemental Fig. S3D and H). Our results conclusively suggest that *OsCLSY3* is an upstream regulator, essential for reproduction and endosperm development.

### The RdDM pathway regulates tissue-specific expression of *OsCLSY3 via* TEs

One of the most striking features of epigenetic regulators of imprinting, operating in endosperm, is their imprinted nature. Imprinting and expression of such players were tightly regulated by epigenetic pathways, usually involving TEs or repeats. To maintain the genome integrity, plants were thought to have evolved silencing involving various epigenetic layers (Quesneville, 2020), including silencing mechanisms that have the ability to spread or influence neighboring genes (Deng et al., 2016). Several such examples exist, for example, in *Arabidopsis,* reproductive-tissue-specific and paternally imprinted *FWA* gene was regulated by SINE retroelements located in its promoter. Here, DNA methylation marks silence the gene in vegetative tissue and paternal allele in endosperm (Kinoshita et al., 2004, 2007). In *Arabidopsis*, a balance between DNA methylation and demethylation was regulated by a TE located in the 5’ flanking sequences of demethylase gene *ROS1* (Williams et al., 2015). Also, DNA binding sites of imprinted gene *PHE1* have TEs of RC/Helitron type (Batista et al., 2019). The paternally imprinted gene *ALLANTOINASE (ALN)* is a negative regulator of seed dormancy in *Arabidopsis*, and it is regulated by a DNA transposon named POGO present in its promoter (Iwasaki et al., 2019). In rice too, a CACTA DNA transposon derived miRNA820 negatively regulated DNA methyltransferase *OsDRM2* by PTGS (Nosaka et al., 2012). Rice tillering is regulated by the RdDM pathway through MITEs elements present at the OsmiRNA156j and D14 gene (Xu et al., 2020). A stowaway-like MITE embedded in the 3’ UTR of the Ghd2 (*CONSTANS*-like gene) regulates its expression through RdDM (Shen et al., 2017). A PcG complex MEG gene *MEA* in *Arabidopsis* is itself silenced by PcG complex in vegetative stages (Jullien et al., 2006). All these studies suggest a strong feedback loop for effective control of a critical process such as endosperm development, acting as a negative deterrent. Our finding of imprinted nature of *OsCLSY3* through two tandem MITE TEs, a Ditto element and a Tourist element, supports this idea. Those TEs were methylated in vegetative tissues by the RdDM pathway where another CLSY namely OsCLSY4 recruited PolIV to regulate its expression. Presence of MITEs as regulatory elements of a crucial gene such as *OsCLSY3* is not entirely surprising. After all, MITEs are the largest family of TEs in the rice genome, located close to more than 23000 genes (nearly 58% of all genes) (Yuan et al., 2017). Most of these MITEs are targets of RdDM and are methylated in most tissues (Zemach et al., 2010). In endosperm, methylation at TEs occupying the *OsCLSY3* promoter was likely abolished due to ROS1a-dependent hypomethylation, similar to other genes under its control (Kim et al., 2019). The expression level of *OsCLSY3* appears to be vital in determining endosperm size, its morphology and functions. OE of *OsCLSY3* produced a larger endosperm with chalkiness (Supplemental Fig. S4I) implicating its role in cellularization. The chalkiness is an indication of starch and protein synthesis and storage defect in endosperm, suggesting that *OsCLSY3* is important for maintaining nutrient quality. In monocot seeds, endosperm plays an important role in seed germination as seen in mis-expression lines of *OsCLSY3* where seed germination rate and initial root development were affected (Supplemental Fig. S4J and K).

**Figure 8.**
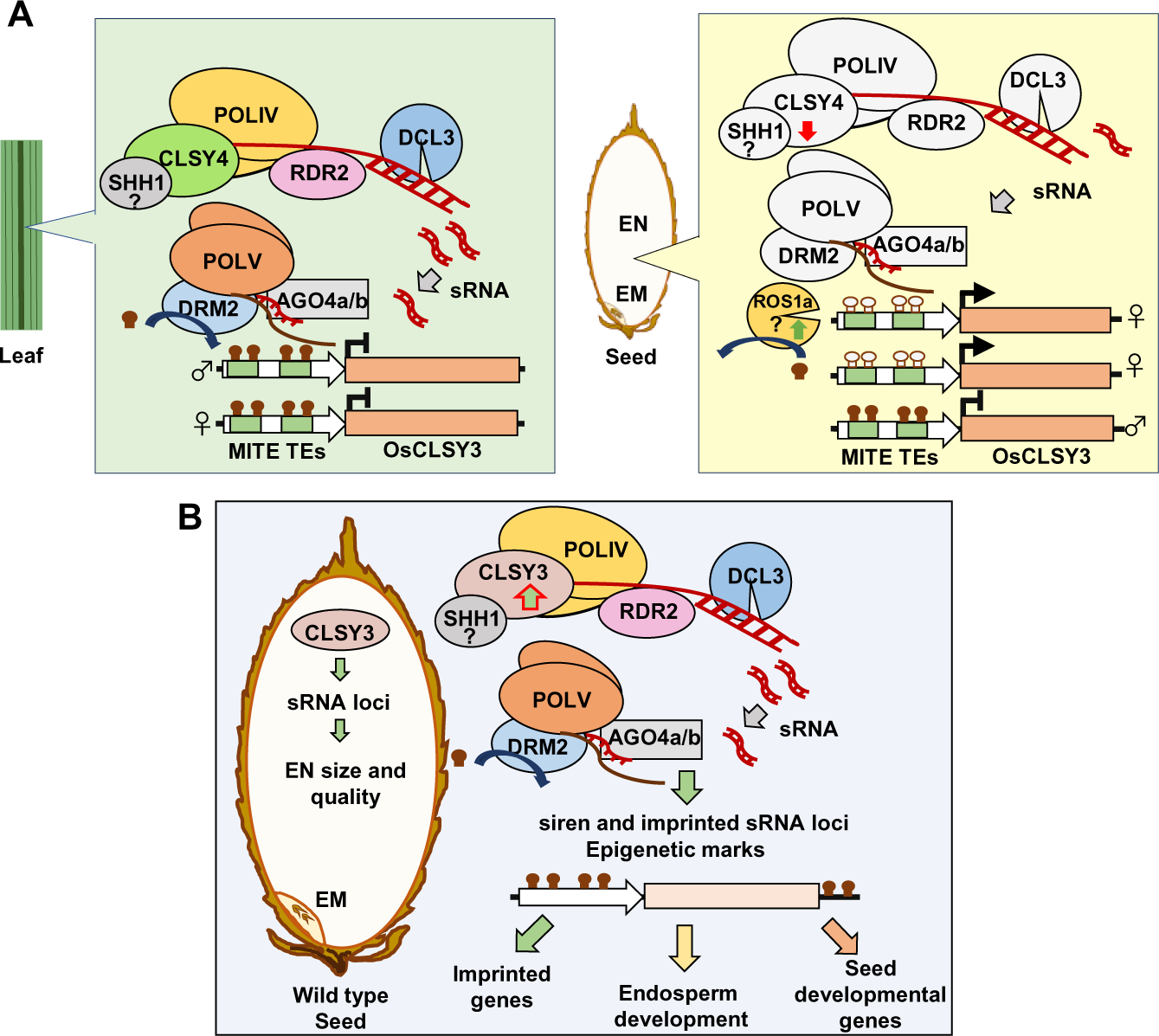
Tissue-preferred expression and functions of *OsCLSY3* in rice endosperm. (A) Tissue-preferred expression of *OsCLSY3* regulated by the RdDM pathway through MITE TEs present in its promoter. In vegetative-tissues, OsCLSY4 recruits PolIV into *OsCLSY3* promoter and methylates MITE TEs *via* 24 nt sRNAs (left panel). In endosperm, OsROS1a induces demethylation of the maternal alleles of *OsCLSY3* promoter leading to its expression (right panel). Here, CLSY4 is not very active to induce silencing in TEs. (B) *OsCLSY3* regulated expression of TE and repeat derived sRNAs, siren loci and imprinted sRNAs in endosperm. Those sRNAs directed DNA methylation and induced associated epigenetic regulations that are crucial for proper development. CLSY3-dependent sRNAs regulated expression of imprinted and seed-development related genes, thereby contributing to endosperm size, cellularization and its quality. Roles of all indicated genes on OsCLSY3-related regulation, except those with question marks, were delineated in this study. Filled and unfilled lollypops indicated methylated and unmethylated DNA, respectively. EM-embryo. EN-endosperm.

### *OsCLSY3* regulates siren loci

In the endosperm of flowering plants, fewer sRNA loci closer to genes contribute a large number of sRNAs in endosperm tissues (Rodrigues et al., 2013; Grover et al., 2020). Such sRNAs, named siren, are a conserved feature of Angiosperms, but they are not identical sequence-wise (Rodrigues et al., 2013; Xin et al., 2014; Grover et al., 2020; Burgess et al., 2022). The siren loci derived sRNAs direct DNA methylation of genes by RdDM (Grover et al., 2020). Recently, studies showed that siren sRNAs can also methylate protein coding genes *in trans* (Burgess et al., 2022). In our study, we showed *OsCLSY3* is the major regulator of siren loci in rice. We also observed siren loci derived sRNAs guide DNA methylation which regulates expression of proximal genes (Fig. 6L). The involvement of *OsCLSY3* in the generation of siren loci is in complete agreement with its larger role as a key regulator of endosperm development.

### *OsCLSY3* regulates expression of many imprinted genes and seed development related genes

In *Arabidopsis*, rice and maize, previous studies indicated that imprinted sRNA loci were located near the silenced allele of few imprinted genes (Mosher et al., 2009; Rodrigues et al., 2013; Xin et al., 2014; Yuan et al., 2017; Erdmann et al., 2017; Chen et al., 2018; Rodrigues et al., 2021). Due to these observations, it was suspected that imprinted sRNAs might be regulating expression of imprinted genes (Erdmann et al., 2017). Same study also documented perturbation of maternal and paternal sRNA ratios in endosperm that can shift the mRNA ratio generated from parental genomes (Erdmann et al., 2017; Satyaki and Gehring, 2022). Unlike *Arabidopsis*, where *PolIV* appears to be a PEG, *PolIV* is not an imprinted gene in rice endosperm probably because *OsCLSY3* is a MEG acting as upstream player. We observed that *OsCLSY3* regulates expression of many imprinted sRNA loci, and adjoining imprinted genes (Fig. 7D and E). In rice, *OsCLSY3* might be regulating transcription of genes via sRNAs as observed in the case of poliv mutant in *Arabidopsis*.

In several examples, CLSY3-dependend sRNAs directly regulated them *via* DNA methylation. Several genes also had expression changes independent of DNA methylation or sRNAs. Such variations might be due to alterations in other epigenetics marks, altered chromatin state or due to mis-regulation of regulatory modules. We observed several auxin, brassinosteroid signaling pathway genes such as *TUD1,OsDWARF/BRD1, BRD2, OsGSK3* which were well documented as important players in grain size (Hu et al., 2013a; Gao et al., 2019; Qin et al., 2018; Huang et al., 2022), were mis-expressed in clsy3-kd lines. Grain development related MAPK pathway genes (Liu et al., 2021; Xu et al., 2018) and transcription factors (TF) such as *OsMADS6, OsMADS29*, *OsWRKY53* which were known for starch synthesis, proper cellularization, cell size regulation and programmed cell death also mis-regulated in clsy3-kd lines (Yin and Xue, 2012; Nayar et al., 2013, 2014; Zhang et al., 2010; Tang et al., 2022). Multiple starch, protein, lipid metabolism related genes such as *OsSGL, RSS, OsSWEET11, OsFAD2, OsLHT1, OsGAD3* that were important for synthesis and their transport, were mis-regulated in clsy3-kd, further indicating a critical role of *OsCLSY3* (Ogawa et al., 2011; Tiwari et al., 2016; Yang et al., 2018b; Peng et al., 2022; Liu et al., 2022b). Since size, quality and nutritive aspects of rice endosperm, a major staple food for the majority, is under *OsCLSY3* control, this gene can be an important candidate to improve nutritional benefits of rice grains.

## Materials and Methods

### Rice transformation and plant growth

For generating rice transgenic plants, Agrobacterium mediated transformation was performed as described previously (Hiei et al., 1997; Sridevi et al., 2003). Briefly, 21-day old embryo-derived calli were obtained from *indica* variety rice (*O. sativa indica*) PB1. Embryogenic calli were infected with freshly grown *A. tumefaciens* strain LBA4404 with *vir* helper plasmid pSB1 (pSB1 carries extra copies of *vir* genes) carrying the binary plasmids of interest was grown until A600 nm = 1.0 OD and was used for infection as described. The regenerated transgenic seedlings were maintained in a growth chamber at 23°C with 16 h/8 h light/dark cycle at 70% RH. The plants were subsequently transferred to a greenhouse.

### Vector design and construction

To generate OE plants, the *OsCLSY3* gene (Os02g0650800) (5.4 kb) was amplified using appropriate primers (Supplemental Table S3) from genomic DNA. The full-length gene was cloned under maize Ubiquitin promoter (P: ZmUbi) in pCAMBIA1300 with *hygromycin phosphotransferase (hph)* gene as a selection marker. The 3xFLAG epitope tag was added to N-terminal of *OsCLSY3*. Two clsy3-kd lines were generated using artificial miRNA (amiR) strategy. The amiRs were designed using the WMD3 web tool (Ossowski et al., 2008; Warthmann et al., 2008), with the stringent criteria for robust amiR generation as described (Narjala et al., 2020). The amiR-precursors were synthesized by GeneArt (Thermofisher). The amiR sequences were cloned into pCAMBIA1300 under P: ZmUbi and into pRT100 vector under CaMV 35S (P:35S) promoter. For constructing a double amiR containing binary vector, an amiR cassette containing pRT100 was cloned into a binary vector that already had P: ZmUbi driven amiR. The KO lines were generated using CRISPR-Cas9 strategy. Guide RNA was designed using CRISPR-PLANT (http://omap.org/crispr/index.html). Guide RNA targeting a unique region of *OsCLSY3* gene at 1st exon (UCCUCUCGGCCCUCCAACAG) was cloned into pRGEB32 vector (Addgene Plasmid #63142) (Xie and Yang, 2013). All the constructs were verified by restriction enzyme-based analysis and Sanger sequencing. Constructs were mobilized into *Agrobacterium* strain LBA4404 (pSB1), and mobilization was verified by polymerase chain reaction (PCR) analysis.

### Genetic crosses

To understand the imprinting status of the *OsCLSY3* gene, two rice varieties - WP and PB1 were crossed. The hybrid was confirmed by Sanger sequencing of known SNP containing regions. Endosperm from the hybrid plant was collected and dissected into embryo and endosperm. Embryo region was used for the DNA isolation and endosperm was used for RNA isolation. After validation of the genotype, cDNA was synthesized from 1.5 µg of total RNA using Thermo Scientific RevertAid First Strand cDNA Synthesis Kit as per manufacturer’s instructions, and regions of interest were amplified. The amplicons were purified and deep-sequenced on NovaSeq 6000 (2 X 100 bp mode). The obtained reads were adapter trimmed using cutadapt (Martin, 2011) and aligned to genes by using CRISPResso2 (Clement et al., 2019).

### 5-Aza-2′-Deoxycytidine treatment

After 3 days of germination in half-strength MS media, seedlings were placed in MS media containing 35 mg/L and 70 mg/L Aza (Chen et al., 2018). DMSO was used as control. After 7 days, seedlings were collected for RNA extraction or GUS staining.

### Phenotyping of transgenic plants

Phenotypes of transgenic plants such as plant height, leaf length, panicle length were measured using (n>6) mature plants grown for about 4 months. Other details including replicates were mentioned in the figure legends. Images of rice spikelets and seeds from WT, clsy3-kd and OE lines were obtained using Lecia S8APO stereomicroscope and Nikon camera. For statistical analysis, paired *t*-test was used.

### RNA extraction and RT-qPCR

Total RNA extraction from rice tissues were performed using TRIzol® Reagent (Invitrogen) as per manufacturer instructions. For endosperm tissue, RNA isolation was performed as described earlier (Wang et al., 2012). RT-qPCR was performed for expression of *CLSY*s and other RdDM pathway related genes. First-strand cDNA was synthesized from 1.5 µg of total RNA using Thermo Scientific RevertAid First Strand cDNA Synthesis Kit as per manufacturer instructions, and the qPCR was carried out with Solis Biodyne - 5x HOT Firepol Evagreen qPCR Master Mix. *OsActin* (Os03g0718100) and *Glyceraldehyde-3-phosphate dehydrogenase* (*OsGAPDH)* (Os04g0486600) were used as internal control. RT-qPCRs were performed at least three times using the BioRad CFX system. Primers used for expression analysis are provided in Supplemental Table S3.

### Northern hybridization

About 8-15 µg of total RNA was used for sRNA northern as described earlier (Shivaprasad et al., 2012; Tirumalai et al., 2020). Membranes were stripped and used for multiple hybridizations. Hybridization was done at 35°C. U6 and miRNA168 were used as loading controls. The blot was exposed to a phosphorimager screen and re-probed with controls. Typhoon scanner (GE healthcare) was used to detect the hybridization signal. Details of the DNA oligonucleotide probes that were used in this study are provided in Supplemental Table S3.

### Southern hybridization

Junction fragment Southern blot analysis was performed as described earlier (Ramanathan and Veluthambi, 1995; Vivek Hari Sundar and Shivaprasad, 2022). Total DNA was extracted from equally grown control and transgenic plants using CTAB method (Rogers and Bendich, 1994). For the probe, *hph* gene was amplified from binary plasmid. The probe was labelled with [α-32P] dCTP (BRIT, India) using the Rediprime random DNA labelling system (GE healthcare). The prehybridization, hybridization and washes were performed at 65°C. The blots were exposed to a phosphorimager screen. Typhoon scanner (GE healthcare) was used to detect the hybridization signal. Details of the probe that was used in this study are provided in Supplemental Table S3.

### DNA methylation analyses

Total DNA extracted by using the CTAB method from the tissues mentioned (Rogers and Bendich, 1994). About 200-400 ng of DNA was bisulfite treated using EZ DNA Methylation-Gold Kit. The bisulfite treated DNA was used as the template for PCR. Amplification of targeted regions performed by JumpStart™ Taq DNA Polymerase (Sigma). The PCR products were deep sequenced in paired end mode (100 bp) on a Hiseq2500 platform. The obtained reads were quality checked and trimmed using cutadapt (Martin, 2011) and aligned to create genome (Target sites) using Bismark aligner tool with default parameters (Krueger and Andrews, 2011). DNA methylation status of those targeted sites was extracted and coverage reports were generated using the Bismark. The obtained results are analyzed using methylation package ViewBS (Huang et al., 2018). Primers used for analysis are provided in Supplemental Table S3.

### sRNA differential expression analyses

The obtained reads were quality checked and trimmed by UEA sRNA Workbench (Stocks et al., 2018). The filtered sRNAs were classified into 21-22 nt and 23-24 nt, aligned and sRNAs loci were identified using Shortstack (Axtell, 2013) with following parameters: --nohp --mmap f --mismatches 1 -mincov 2rpmm. The counts obtained were used for differential expression analyses using DESeq2 (Love et al., 2014). For quantifying the sRNA expression from transposons, siren loci etc. bedtools multicov (Quinlan and Hall, 2010) was used to obtain raw abundance and then normalised to RPKM values. These values are plotted as box plots using custom R scripts in ggplot2 (Wickham, 2016)

### RNA-seq and analyses

RNA seq was performed using 20 days old endosperm tissues. The total RNA was extracted using previously described methods. Poly(A) enrichment was done before library preparation. Library preparation was done with NEBNext® Ultra™ II Directional RNA Library Prep kit (E7765L) as per manufacturer’s instructions. The obtained libraries were sequenced in paired end mode (100 bp) on the Illumina Hiseq2500 platform. The obtained reads were adapter trimmed using Trimmomatic (Bolger et al., 2014). The reads were aligned to IRGSP1.0 genome using HISAT2 (Kim et al., 2015) with default parameters. Cufflinks was used to perform differential gene expression (DEGs) analyses and statistical testing (Trapnell et al., 2012). The volcano plots were generated for DEGs using custom R scripts with the p-value cut-off of less than 0.05 and absolute log2 (fold change) expression cut-off of more than 1.5. For quantifying the expression of genes, bedtools multicov (Quinlan and Hall, 2010; Quinlan, 2014; Patwardhan et al., 2019) was used to obtain raw abundance and then normalized to RPKM values. These values are plotted as box plots and heatmaps using custom R scripts in ggplot2 (Wickham, 2016).

### AGO-IP data analyses

Previously published, AGO IP datasets of various AGOs from rice were processed in the same way as the sRNA datasets, mapping to the IRGSP1.0 genome (Quinlan and Hall, 2010; Quinlan, 2014).

### Chop-qPCR

The chop qPCR performed with 20 days endosperm DNA of clsy3-kd and WT as described (Chakraborty et al., 2022). Total DNA was isolated by CTAB method and 500 ng DNA digested with *NlaIII* (10 K units/ml) restriction enzyme for 2 h. The digested DNA was used as template for qPCR. Equally treated DNA, without any enzyme was used as mock. Target loci as well as ACTIN with *NlaIII* restriction site, was used as a control. qPCRs were performed at least twice using BioRad CFX system. Primers used for chop qPCR are provided in Supplemental Table S3.

### GUS histochemical assay

Histochemical assay was performed as described (Jefferson et al., 1987). Plant tissues were collected in 50 mM phosphate buffer that contained 1% Triton-X 100 and incubated for 3 h at 37°C. Explants were transferred into X-Gluc staining solution (1 mM) and vacuum infiltrated for 15 min. Next, the explants were incubated at 37°C for 16 h. Further, explants were washed once with double distilled water followed by 70% ethanol and then transferred to acetone: methanol mix (1:3) in rotospin at 4°C for 1 h to remove chlorophyll. After chlorophyll removal, explants were imaged by Nikon camera or Leica S8APO stereomicroscope.

### SEM imaging

For scanning electron microscope (SEM) imaging, rice spikelets were collected just before flowering. Samples were fixed in 16% Formaldehyde, 25% Glutaraldehyde and 0.2 M cacodylate buffer for 12-16 h. The samples were rinsed with double distilled water and dehydrated in series of ethanol and dried in CPD (Leica EM CPD300), gold coated, and the images were obtained using a Carl Zeiss scanning electron microscope at an accelerating voltage of 2 kV-4 kV as described before (Pachamuthu et al., 2021; Hari Sundar G et al., 2023).

### Germination assay

The germination assay was performed as described previously (He et al., 2020). Total 20 seeds of different genotypes (five seeds per replicate) were imbibed on the wet filter paper in 5 cm diameter petri-plates. In all the plates, 4 ml of single distilled water was used to wet the filter papers and seeds were germinated at 25°C for 6 days in the dark. Experiments were repeated twice independently.

### Pollen staining

Pollen viability test was performed using I2-KI staining solution containing 0.2% (w/v) I2 and 2% (w/v) KI as described earlier (Khatun and Flowers, 1995; Yao et al., 2018; Pachamuthu et al., 2021). Anthers from six spikelets of mature panicles were collected in 200 μl of solution one day before the fertilization. Pollen grains were released in the solution by mechanical shearing. After 10 min, viable pollen grains were counted under the bright-field microscope. Round and dark blue stained pollen grains were considered as viable, while very light blue and distorted pollens were considered as non-viable (Das et al., 2020; Hari Sundar G et al., 2023).

## Data access

All raw and processed sequencing data generated in this study have been submitted to the NCBI Gene Expression Omnibus (GEO; https://www.ncbi.nlm.nih.gov/geo/) under accession number GSEXXXXXX.

## Acknowledgements

We thank Prof. K. Veluthambi for *Agrobacterium* strains, WP, PB1 seeds and binary plasmids. We thank genomics, electron microscopy, CIFF, IT, radiation, greenhouse, and lab-kitchen facilities at the NCBS. We acknowledge the guidance of Anushree Narjala in bioinformatics analysis. We acknowledge the guidance of Dhandapani Murugesan and his team in rice crossing. We thank all the lab members for discussions and comments.

This work was supported by NCBS-TIFR core funding and a grant (BT/PR25767/GET/ 119/151/2017) from Department of Biotechnology (DBT), Government of India. This study was also supported by the Department of Atomic Energy, Government of India, under Project Identification No. RTI 4006 (1303/3/2019/R&D-II/DAE/4749 dated 16.7.2020). These funding agencies did not participate in the designing of experiments, analysis, or interpretation of data, or in writing of the manuscript.

## Author contributions

PVS and AKP designed all experiments and discussed results and wrote the manuscript. AKP performed most of the experiments and bioinformatics analyses. VHS generated poliv-kd datasets and performed bioinformatics analysis. AN performed embryo and endosperm transcriptome analysis. All authors have read and approved the manuscript.

## Supplemental information

This manuscript has 9 supplemental figures, 4 supplemental tables and 9 supplemental datasets.

## Supplemental Figures

Supplemental Figure S1. Transcriptome analysis of endosperm and embryo tissues.

Supplemental Figure S2. OsCLSY3 is an endosperm-preferred gene among monocots.

Supplemental Figure S3. KO transgenic plants show reproductive abnormalities.

Supplemental Figure S4. Validation of OsCLSY3 transgenic lines.

Supplemental Figure S5. Endosperm sRNAs are globally reduced in clsy3-kd.

Supplemental Figure S6. The RdDM pathway regulates expression of siren loci in rice.

Supplemental Figure S7. CLSY3-dependent sRNAs regulate protein coding genes.

Supplemental Figure S8. The RdDM pathway controls imprinted sRNA loci in rice endosperm.

Supplemental Figure S9. DNA methylation levels in imprinted genes of embryo and endosperm.

## Supplemental Tables

Supplemental Table S1. Details of high-throughput genomics data generated in this study.

Supplemental Table S2. Details of high-throughput genomics data obtained from publicly available datasets.

Supplemental Table S3. List of oligos and probes used in this study.

Supplemental Table S4. Sequences of DRD1 family proteins used in this study.

## Supplemental datasets

Supplemental dataset S1. List of Log_2_ 2.0-fold change upregulated genes mature endosperm and embryo (endosperm-preferred genes).

Supplemental dataset S2. List of Log_2_ 2.0-fold change downregulated genes mature endosperm and embryo (embryo-preferred genes).

Supplemental dataset S3. List of epigenetic genes and all published imprinted genes in rice.

Supplemental dataset S4. List of all Shortstack 23-24 nt sRNA loci in WT endosperm.

Supplemental dataset S5. List of all Shortstack 23-24 nt sRNA loci in clsy3-kd endosperm.

Supplemental dataset S6. List of CLSY3-dependent 23-24 nt sRNA loci.

Supplemental dataset S7. List of published rice siren loci with their normalized expression.

Supplemental dataset S8. List of log_2_1.5-fold change upregulated genes in clsy3-kd endosperm.

Supplemental dataset S9. List of log_2_1.5-fold change downregulated genes in clsy3-kd endosperm.

## References

An, L., Tao, Y., Chen, H., He, M., Xiao, F., Li, G., Ding, Y., and Liu, Z. (2020). Embryo-endosperm interaction and its agronomic relevance to rice quality. Front. Plant Sci. 11: 587641.

Anderson, S.N., Zhou, P., Higgins, K., Brandvain, Y., and Springer, N.M. (2021). Widespread imprinting of transposable elements and variable genes in the maize endosperm. PLoS Genet. 17: e1009491.

Anushree, N. and Shivaprasad, P.V. (2017). Regulation of Plant miRNA Biogenesis. Proc. Indian Natl. Sci. Acad. (A Phys. Sci.) 95.

Axtell, M.J. (2013). ShortStack: comprehensive annotation and quantification of small RNA genes. RNA 19: 740–751.

Baroux, C., Spillane, C., and Grossniklaus, U. (2002). Evolutionary origins of the endosperm in flowering plants. Genome Biol. 3: reviews1026.

Batista, R.A. and Köhler, C. (2020). Genomic imprinting in plants-revisiting existing models. Genes Dev. 34: 24–36.

Batista, R.A., Moreno-Romero, J., Qiu, Y., van Boven, J., Santos-González, J., Figueiredo, D.D., and Köhler, C. (2019). The MADS-box transcription factor PHERES1 controls imprinting in the endosperm by binding to domesticated transposons. Elife 8.

Bolger, A.M., Lohse, M., and Usadel, B. (2014). Trimmomatic: a flexible trimmer for Illumina sequence data. Bioinformatics 30: 2114–2120.

Burgess, D., Chow, H.T., Grover, J.W., Freeling, M., and Mosher, R.A. (2022). Ovule siRNAs methylate protein-coding genes in trans. Plant Cell 34: 3647–3664.

Castano-Duque, L., Ghosal, S., Quilloy, F.A., Mitchell-Olds, T., and Dixit, S. (2021). An epigenetic pathway in rice connects genetic variation to anaerobic germination and seedling establishment. Plant Physiol. 186: 1042–1059.

Chahtane, H., Nogueira Füller, T., Allard, P.-M., Marcourt, L., Ferreira Queiroz, E., Shanmugabalaji, V., Falquet, J., Wolfender, J.-L., and Lopez-Molina, L. (2018). The plant pathogen Pseudomonas aeruginosa triggers a DELLA-dependent seed germination arrest in Arabidopsis. Elife 7.

Chakraborty, T., Trujillo, J.T., Kendall, T., and Mosher, R.A. (2022). A null allele of the pol IV second subunit impacts stature and reproductive development in Oryza sativa. Plant J. 111: 748–755.

Chen, C., Begcy, K., Liu, K., Folsom, J.J., Wang, Z., Zhang, C., and Walia, H. (2016). Heat stress yields a unique MADS box transcription factor in determining seed size and thermal sensitivity. Plant Physiol. 171: 606–622.

Chen, C., Li, T., Zhu, S.[.]., Liu, Z., Shi, Z., Zheng, X., Chen, R., Huang, J., Shen, Y., Luo, S., and Wang, L. (2018). Characterization of imprinted genes in rice reveals conservation of regulation and imprinting with other plant species. Plant Physiology 177: 1754–1771.

Chen, X. and Zhou, D.-X. (2013). Rice epigenomics and epigenetics: challenges and opportunities. Curr. Opin. Plant Biol. 16: 164–169.

Cheng, X., Pan, M. E Z., Zhou, Y., Niu, B., and Chen, C. (2020). Functional divergence of two duplicated Fertilization Independent Endosperm genes in rice with respect to seed development. Plant J. 104: 124–137.

Cheng, X., Pan, M. E. Z., Zhou, Y., Niu, B., and Chen, C. (2021). The maternally expressed polycomb group gene OsEMF2a is essential for endosperm cellularization and imprinting in rice. Plant Commun. 2: 100092.

Chow, H.T. and Mosher, R.A. (2023). Small RNA-mediated DNA methylation during plant reproduction. Plant Cell 35: 1787–1800.

Clement, K., Rees, H., Canver, M.C., Gehrke, J.M., Farouni, R., Hsu, J.Y., Cole, M.A., Liu, D.R., Joung, J.K., Bauer, D.E., and Pinello, L. (2019). CRISPResso2 provides accurate and rapid genome editing sequence analysis. Nat. Biotechnol. 37: 224–226.

Das, S., Swetha, C., Pachamuthu, K., Nair, A., and Shivaprasad, P.V. (2020). Loss of function of Oryza sativa Argonaute 18 induces male sterility and reduction in phased small RNAs. Plant Reprod. 33: 59–73.

De Giorgi, J. et al. (2021). The Arabidopsis mature endosperm promotes seedling cuticle formation via release of sulfated peptides. Dev. Cell 56: 3066–3081.e5.

Deng, X., Song, X., Wei, L., Liu, C., and Cao, X. (2016). Epigenetic regulation and epigenomic landscape in rice. Natl. Sci. Rev. 3: 309–327.

Dhatt, B.K., Paul, P., Sandhu, J., Hussain, W., Irvin, L., Zhu, F., Adviento-Borbe, M.A., Lorence, A., Staswick, P., Yu, H., Morota, G., and Walia, H. (2021). Allelic variation in rice Fertilization Independent Endosperm 1 contributes to grain width under high night temperature stress. New Phytol. 229: 335–350.

Doll, N.M., Royek, S., Fujita, S., Okuda, S., Chamot, S., Stintzi, A., Widiez, T., Hothorn, M., Schaller, A., Geldner, N., and Ingram, G. (2020). A two-way molecular dialogue between embryo and endosperm is required for seed development. Science 367: 431–435.

Erdmann, R.M., Satyaki, P.R.V., Klosinska, M., and Gehring, M. (2017). A small RNA pathway mediates Allelic dosage in endosperm. Cell Rep. 21: 3364–3372.

Gao, X. et al. (2019). Rice qGL3/OsPPKL1 functions with the GSK3/SHAGGY-like kinase OsGSK3 to modulate brassinosteroid signaling. Plant Cell 31: 1077–1093.

Gehring, M. (2013). Genomic imprinting: insights from plants. Annu. Rev. Genet. 47: 187– 208.

Gehring, M., Hsieh, T.F., Penterman, J., Choi, Y., Harada, J.J., and Fischer, R.L. (2006). DEMETER DNA glycosylase establishes MEDEA Polycomb gene self-imprinting by allele-specific demethylation Cell. Cell 124: 495–506.

Griffiths-Jones, S., Saini, H.K., van Dongen, S., and Enright, A.J. (2008). miRBase: tools for microRNA genomics. Nucleic Acids Res. 36: D154–8.

Grossniklaus, U., Vielle-Calzada, J.P., Hoeppner, M.A., and Gagliano, W.B. (1998). Maternal control of embryogenesis by MEDEA, a polycomb group gene in Arabidopsis. Science 280: 446–450.

Grover, J.W., Burgess, D., Kendall, T., Baten, A., Pokhrel, S., King, G.J., Meyers, B.C., Freeling, M., and Mosher, R.A. (2020). Abundant expression of maternal siRNAs is a conserved feature of seed development. Proc. Natl. Acad. Sci. U. S. A. 117: 15305–15315.

Grover, J.W., Kendall, T., Baten, A., Burgess, D., Freeling, M., King, G.J., and Mosher, R.A. (2018). Maternal components of RNA-directed DNA methylation are required for seed development in Brassica rapa. Plant J. 94: 575–582.

Gutierrez-Marcos, J.F., Pennington, P.D., Costa, L.M., and Dickinson, H.G. (2003). Imprinting in the endosperm: a possible role in preventing wide hybridization. Philos. Trans. R. Soc. Lond. B Biol. Sci. 358: 1105–1111.

Hale, C.J., Stonaker, J.L., Gross, S.M., and Hollick, J.B. (2007). A novel Snf2 protein maintains trans-generational regulatory states established by paramutation in maize. PLoS Biol. 5: e275.

Hari Sundar G, V., Swetha, C., Basu, D., Pachamuthu, K., Raju, S., Chakraborty, T., Mosher, R.A., and Shivaprasad, P.V. (2023). Plant polymerase IV sensitizes chromatin through histone modifications to preclude spread of silencing into protein-coding domains. Genome Res.

He, Y., Zhao, J., Yang, B., Sun, S., Peng, L., and Wang, Z. (2020). Indole-3-acetate beta-glucosyltransferase OsIAGLU regulates seed vigour through mediating crosstalk between auxin and abscisic acid in rice. Plant Biotechnol. J. 18: 1933–1945.

Hiei, Y., Komari, T., and Kubo, T. (1997). Transformation of rice mediated by Agrobacterium tumefaciens. Plant Mol. Biol. 35: 205–218.

Higo, A. et al. (2020). DNA methylation is reconfigured at the onset of reproduction in rice shoot apical meristem. Nat. Commun. 11: 4079.

Hsieh, T.-F., Shin, J., Uzawa, R., Silva, P., Cohen, S., Bauer, M.J., Hashimoto, M., Kirkbride, R.C., Harada, J.J., Zilberman, D., and Fischer, R.L. (2011). Regulation of imprinted gene expression in Arabidopsis endosperm. Proc. Natl. Acad. Sci. U. S. A. 108: 1755–1762.

Hu, D. et al. (2022). Erratum for: Multiplex CRISPR-Cas9 editing of DNA methyltransferases in rice uncovers a class of non-CG methylation specific for GC-rich regions. Plant Cell 34: 1416.

Hu, X., Qian, Q., Xu, T., Zhang, Y., Dong, G., Gao, T., Xie, Q., and Xue, Y. (2013a). The U-box E3 ubiquitin ligase TUD1 functions with a heterotrimeric G α subunit to regulate Brassinosteroid-mediated growth in rice. PLoS Genet. 9: e1003391.

Hu, Y., Zhu, N., Wang, X., Yi, Q., Zhu, D., Lai, Y., and Zhao, Y. (2013b). Analysis of rice Snf2 family proteins and their potential roles in epigenetic regulation. Plant Physiol. Biochem. 70: 33–42.

Huang, F., Zhu, Q.-H., Zhu, A., Wu, X., Xie, L., Wu, X., Helliwell, C., Chaudhury, A., Finnegan, E.J., and Luo, M. (2017). Mutants in the imprinted PICKLE RELATED 2 gene suppress seed abortion of fertilization independent seed class mutants and paternal excess interploidy crosses in Arabidopsis. Plant J. 90: 383–395.

Huang, J. et al. (2022). Natural variation of the BRD2 allele affects plant height and grain size in rice. Planta 256: 27.

Huang, X., Lu, Z., Wang, X., Ouyang, Y., Chen, W., Xie, K., Wang, D., Luo, M., Luo, J., and Yao, J. (2016). Imprinted gene OsFIE1 modulates rice seed development by influencing nutrient metabolism and modifying genome H3K27me3. Plant J. 87: 305– 317.

Huang, X., Zhang, S., Li, K., Thimmapuram, J., and Xie, S. (2018). ViewBS: a powerful toolkit for visualization of high-throughput bisulfite sequencing data. Bioinformatics 34: 708–709.

Ishikawa, R., Ohnishi, T., Kinoshita, Y., Eiguchi, M., Kurata, N., and Kinoshita, T. (2011). Rice interspecies hybrids show precocious or delayed developmental transitions in the endosperm without change to the rate of syncytial nuclear division. Plant J. 65: 798–806.

Iwasaki, M., Hyvärinen, L., Piskurewicz, U., and Lopez-Molina, L. (2019). Non-canonical RNA-directed DNA methylation participates in maternal and environmental control of seed dormancy. Elife 8.

Iwasaki, M., Penfield, S., and Lopez-Molina, L. (2022). Parental and Environmental Control of Seed Dormancy in Arabidopsis thaliana. Annu. Rev. Plant Biol. 73: 355– 378.

Jefferson, R.A., Kavanagh, T.A., and Bevan, M.W. (1987). GUS fusions: beta-glucuronidase as a sensitive and versatile gene fusion marker in higher plants. EMBO J. 6: 3901–3907.

Jiang, H. and Köhler, C. (2012). Evolution, function, and regulation of genomic imprinting in plant seed development. J. Exp. Bot. 63: 4713–4722.

Jullien, P.E., Katz, A., Oliva, M., Ohad, N., and Berger, F. (2006). Polycomb group complexes self-regulate imprinting of the Polycomb group gene MEDEA in Arabidopsis. Curr. Biol. 16: 486–492.

Khatun, S. and Flowers, T.J. (1995). The estimation of pollen viability in rice. J. Exp. Bot. 46: 151–154.

Kim, D., Langmead, B., and Salzberg, S.L. (2015). HISAT: a fast spliced aligner with low memory requirements. Nat. Methods 12: 357–360.

Kim, M.Y., Ono, A., Scholten, S., Kinoshita, T., Zilberman, D., Okamoto, T., and Fischer, R.L. (2019). DNA demethylation by ROS1a in rice vegetative cells promotes methylation in sperm. Proc. Natl. Acad. Sci. U. S. A. 116: 9652–9657.

Kinoshita, T. (2007). Reproductive barrier and genomic imprinting in the endosperm of flowering plants. Genes Genet. Syst. 82: 177–186.

Kinoshita, T., Choi, Y., Kinoshita, Y., Cao, X., Jacobsen, S.E., and Fischer, R.L. (2004). One-way control of FWA imprinting in Arabidopsis endosperm by DNA methylation Science. Science 303: 521–523.

Kinoshita, Y., Saze, H., Kinoshita, T., Miura, A., Soppe, W.J.J., Koornneef, M., and Kakutani, T. (2007). Control of FWA gene silencing in Arabidopsis thaliana by SINE-related direct repeats. Plant J. 49: 38–45.

Kirkbride, R.C., Lu, J., Zhang, C., Mosher, R.A., Baulcombe, D.C., and Chen, Z.J. (2019). Maternal small RNAs mediate spatial-temporal regulation of gene expression, imprinting, and seed development in Arabidopsis. Proc. Natl. Acad. Sci. U. S. A. 116: 2761–2766.

Kiyosue, T., Ohad, N., Yadegari, R., Hannon, M., Dinneny, J., Wells, D., Katz, A., Margossian, L., Harada, J.J., Goldberg, R.B., and Fischer, R.L. (1999). Control of fertilization-independent endosperm development by the MEDEA polycomb gene in Arabidopsis. Proc. Natl. Acad. Sci. U. S. A. 96: 4186–4191.

Köhler, C., Page, D.R., Gagliardini, V., and Grossniklaus, U. (2005). The Arabidopsis thaliana MEDEA Polycomb group protein controls expression of PHERES1 by parental imprinting. Nat. Genet. 37: 28–30.

Köhler, C. and Weinhofer-Molisch, I. (2010). Mechanisms and evolution of genomic imprinting in plants. Heredity (Edinb.) 105: 57–63.

Kradolfer, D., Wolff, P., Jiang, H., Siretskiy, A., and Köhler, C. (2013). An imprinted gene underlies postzygotic reproductive isolation in Arabidopsis thaliana. Dev. Cell 26: 525–535.

Krueger, F. and Andrews, S.R. (2011). Bismark: a flexible aligner and methylation caller for Bisulfite-Seq applications. Bioinformatics 27: 1571–1572.

Law, J.A., Vashisht, A.A., Wohlschlegel, J.A., and Jacobsen, S.E. (2011). SHH1, a homeodomain protein required for DNA methylation, as well as RDR2, RDM4, and chromatin remodeling factors, associate with RNA polymerase IV. PLoS Genet. 7: e1002195.

Le, B.H. et al. (2010). Global analysis of gene activity during Arabidopsis seed development and identification of seed-specific transcription factors. Proc. Natl. Acad. Sci. U. S. A. 107: 8063–8070.

Li, H., Li, J., Xu, R., Qin, R., Song, F., Li, L., Wei, P., and Yang, J. (2018). Isolation of five rice nonendosperm tissue-expressed promoters and evaluation of their activities in transgenic rice. Plant Biotechnol. J. 16: 1138–1147.

Li, J. and Berger, F. (2012). Endosperm: food for humankind and fodder for scientific discoveries. New Phytol. 195: 290–305.

Li, N., Xu, R., and Li, Y. (2019). Molecular networks of seed size control in plants. Annu. Rev. Plant Biol. 70: 435–463.

Li, P., Chen, Y.-H., Lu, J., Zhang, C.-Q., Liu, Q.-Q., and Li, Q.-F. (2022). Genes and their molecular functions determining seed structure, components, and quality of rice. Rice (N. Y.) 15: 18.

Liu, H., Guo, S., Xu, Y., Li, C., Zhang, Z., Zhang, D., Xu, S., Zhang, C., and Chong, K. (2014). OsmiR396d-regulated OsGRFs function in floral organogenesis in rice through binding to their targets OsJMJ706 and OsCR4. Plant Physiol. 165: 160–174.

Liu, J., Wu, M.-W., and Liu, C.-M. (2022a). Cereal endosperms: Development and storage product accumulation. Annu. Rev. Plant Biol. 73: 255–291.

Liu, Z. et al. (2021). OsMKKK70 regulates grain size and leaf angle in rice through the OsMKK4-OsMAPK6-OsWRKY53 signaling pathway. J. Integr. Plant Biol. 63: 2043– 2057.

Liu, Z., Jiang, S., Jiang, L., Li, W., Tang, Y., He, W., Wang, M., Xing, J., Cui, Y., Lin, Q., Yu, F., and Wang, L. (2022b). Transcription factor OsSGL is a regulator of starch synthesis and grain quality in rice. J. Exp. Bot. 73: 3417–3430.

Long, J., Walker, J., She, W., Aldridge, B., Gao, H., Deans, S., Vickers, M., and Feng, X. (2021). Nurse cell--derived small RNAs define paternal epigenetic inheritance in Arabidopsis. Science 373: eabh0556.

Love, M.I., Huber, W., and Anders, S. (2014). Moderated estimation of fold change and dispersion for RNA-seq data with DESeq2. bioRxiv.

Lu, J., Zhang, C., Baulcombe, D.C., and Chen, Z.J. (2012). Maternal siRNAs as regulators of parental genome imbalance and gene expression in endosperm of Arabidopsis seeds. Proc. Natl. Acad. Sci. U. S. A. 109: 5529–5534.

Luo, M., Bilodeau, P., Koltunow, A., Dennis, E.S., Peacock, W.J., and Chaudhury, A.M. (1999). Genes controlling fertilization-independent seed development in Arabidopsis thaliana. Proc. Natl. Acad. Sci. U. S. A. 96: 296–301.

Luo, M., Platten, D., Chaudhury, A., Peacock, W.J., and Dennis, E.S. (2009). Expression, imprinting, and evolution of rice homologs of the polycomb group genes. Mol. Plant 2: 711–723.

Luo, M., Taylor, J.M., Spriggs, A., Zhang, H., Wu, X., Russell, S., Singh, M., and Koltunow, A. (2011). A genome-wide survey of imprinted genes in rice seeds reveals imprinting primarily occurs in the endosperm. PLoS Genet. 7: e1002125.

Mahto, A., Mathew, I.E., and Agarwal, P. (2017). Decoding the transcriptome of rice seed during development. In Advances in Seed Biology (InTech).

Martin, M. (2011). Cutadapt removes adapter sequences from high-throughput sequencing reads. EMBnet J. 17: 10.

Martins, L.M. and Law, J.A. (2023). Moving targets: Mechanisms regulating siRNA production and DNA methylation during plant development. Curr. Opin. Plant Biol. 75: 102435.

Matzke, M.A., Kanno, T., and Matzke, A.J.M. (2015). RNA-directed DNA methylation: The evolution of a complex epigenetic pathway in flowering plants. Annu. Rev. Plant Biol. 66: 243–267.

Mosher, R.A. (2010). Maternal control of Pol IV-dependent siRNAs in Arabidopsis endosperm. New Phytol. 186: 358–364.

Mosher, R.A., Melnyk, C.W., Kelly, K.A., Dunn, R.M., Studholme, D.J., and Baulcombe, D.C. (2009). Uniparental expression of PolIV-dependent siRNAs in developing endosperm of Arabidopsis. Nature 460: 283–286.

Musselman, C.A. and Kutateladze, T.G. (2021). Characterization of functional disordered regions within chromatin-associated proteins. iScience 24: 102070.

Narjala, A., Nair, A., Tirumalai, V., Hari Sundar, G.V., and Shivaprasad, P.V. (2020). A conserved sequence signature is essential for robust plant miRNA biogenesis. Nucleic Acids Res. 48: 3103–3118.

Nayar, S., Kapoor, M., and Kapoor, S. (2014). Post-translational regulation of rice MADS29 function: homodimerization or binary interactions with other seed-expressed MADS proteins modulate its translocation into the nucleus. J. Exp. Bot. 65: 5339–5350.

Nayar, S., Sharma, R., Tyagi, A.K., and Kapoor, S. (2013). Functional delineation of rice MADS29 reveals its role in embryo and endosperm development by affecting hormone homeostasis. J. Exp. Bot. 64: 4239–4253.

Nosaka, M., Itoh, J.-I., Nagato, Y., Ono, A., Ishiwata, A., and Sato, Y. (2012). Role of transposon-derived small RNAs in the interplay between genomes and parasitic DNA in rice. PLoS Genet. 8: e1002953.

Ogawa, D. et al. (2011). RSS1 regulates the cell cycle and maintains meristematic activity under stress conditions in rice. Nat. Commun. 2: 278.

Ohad, N., Margossian, L., Hsu, Y.C., Williams, C., Repetti, P., and Fischer, R.L. (1996). A mutation that allows endosperm development without fertilization. Proc. Natl. Acad. Sci. U. S. A. 93: 5319–5324.

Olsen, O.-A. (2020). The modular control of cereal endosperm development. Trends Plant Sci. 25: 279–290.

Ossowski, S., Schwab, R., and Weigel, D. (2008). Gene silencing in plants using artificial microRNAs and other small RNAs. Plant J. 53: 674–690.

Pachamuthu, K., Swetha, C., Basu, D., Das, S., Singh, I., Sundar, V.H., Sujith, T.N., and Shivaprasad, P.V. (2021). Rice-specific Argonaute 17 controls reproductive growth and yield-associated phenotypes. Plant Mol. Biol. 105: 99–114.

Patwardhan, M.N., Wenger, C.D., Davis, E.S., and Phanstiel, D.H. (2019). Bedtoolsr: An R package for genomic data analysis and manipulation. J. Open Source Softw. 4: 1742.

Paul, P. et al. (2020). MADS78 and MADS79 are essential regulators of early seed development in rice. Plant Physiol. 182: 933–948.

Peng, B., Lou, A.-Q., Peng, J., Zhang, Q.-X., Sun, X.-Y., Liu, Y., Xu, X.-J., Sun, Y.-Y., Huang, Y.-Q., Song, X.-H., and Wang, Q.-X. (2022). Scanning electron microscopic observation on chalkiness of rice mutant OsLHT1 grains. J. Agric. Sci. 14: 54.

Pires, N.D. (2014). Seed evolution: parental conflicts in a multi-generational household. Biomol. Concepts 5: 71–86.

Qin, R., Zeng, D., Yang, C., Akhter, D., Alamin, M., Jin, X., and Shi, C. (2018). LTBSG1, a new allele of BRD2, regulates panicle and grain development in rice by brassinosteroid biosynthetic pathway. Genes (Basel) 9: 292.

Quesneville, H. (2020). Twenty years of transposable element analysis in the Arabidopsis thaliana genome. Mob. DNA 11: 28.

Quevillon, E., Silventoinen, V., Pillai, S., Harte, N., Mulder, N., Apweiler, R., and Lopez, R. (2005). InterProScan: protein domains identifier. Nucleic Acids Res. 33: W116–20.

Quinlan, A.R. (2014). BEDTools: The Swiss-army tool for genome feature analysis. Curr. Protoc. Bioinformatics 47: 11.12.1-34.

Quinlan, A.R. and Hall, I.M. (2010). BEDTools: a flexible suite of utilities for comparing genomic features. Bioinformatics 26: 841–842.

Ramanathan, V. and Veluthambi, K. (1995). Transfer of non-T-DNA portions of the Agrobacterium tumefaciens Ti plasmid pTiA6 from the left terminus of TL-DNA. Plant Mol. Biol. 28: 1149–1154.

Rodrigues, J.A., Hsieh, P.-H., Ruan, D., Nishimura, T., Sharma, M.K., Sharma, R., Ye, X., Nguyen, N.D., Nijjar, S., Ronald, P.C., Fischer, R.L., and Zilberman, D. (2021). Divergence among rice cultivars reveals roles for transposition and epimutation in ongoing evolution of genomic imprinting. Proc. Natl. Acad. Sci. U. S. A. 118: e2104445118.

Rodrigues, J.A., Ruan, R., Nishimura, T., Sharma, M.K., Sharma, R., Ronald, P.C., Fischer, R.L., and Zilberman, D. (2013). Imprinted expression of genes and small RNA is associated with localized hypomethylation of the maternal genome in rice endosperm. Proc. Natl. Acad. Sci. U. S. A. 110: 7934–7939.

Rogers, S.O. and Bendich, A.J. (1994). Extraction of total cellular DNA from plants, algae and fungi. In Plant Molecular Biology Manual (Springer Netherlands: Dordrecht), pp. 183–190.

Satyaki, P.R.V. and Gehring, M. (2017). DNA methylation and imprinting in plants: machinery and mechanisms. Crit. Rev. Biochem. Mol. Biol. 52: 163–175.

Satyaki, P.R.V. and Gehring, M. (2022). RNA Pol IV induces antagonistic parent-of-origin effects on Arabidopsis endosperm. PLoS Biol. 20: e3001602.

Shen, J., Liu, J., Xie, K., Xing, F., Xiong, F., Xiao, J., Li, X., and Xiong, L. (2017). Translational repression by a miniature inverted-repeat transposable element in the 3’ untranslated region. Nat. Commun. 8: 14651.

Shi, J., Dong, A., and Shen, W.-H. (2014). Epigenetic regulation of rice flowering and reproduction. Front. Plant Sci. 5: 803.

Shivaprasad, P.V., Dunn, R.M., Santos, B.A., Bassett, A., and Baulcombe, D.C. (2012). Extraordinary transgressive phenotypes of hybrid tomato are influenced by epigenetics and small silencing RNAs. EMBO J. 31: 257–266.

Sigrist, C.J.A., Cerutti, L., Hulo, N., Gattiker, A., Falquet, L., Pagni, M., Bairoch, A., and Bucher, P. (2002). PROSITE: a documented database using patterns and profiles as motif descriptors. Brief. Bioinform. 3: 265–274.

Smith, L.M., Pontes, O., Searle, I., Yelina, N., Yousafzai, F.K., Herr, A.J., Pikaard, C.S., and Baulcombe, D.C. (2007). An SNF2 protein associated with nuclear RNA silencing and the spread of a silencing signal between cells in Arabidopsis. Plant Cell 19: 1507–1521.

Sridevi, G., Sabapathi, N., Meena, P., Nandakumar, R., Samiyappan, R., Muthukrishnan, S., and Veluthambi, K. (2003). Transgenic indica rice variety Pusa basmati 1 constitutively expressing a rice chitinase gene exhibits enhanced resistance to Rhizoctonia solani. J. Plant Biochem. Biotechnol. 12: 93–101.

Stocks, M.B., Mohorianu, I., Beckers, M., Paicu, C., Moxon, S., Thody, J., Dalmay, T., and Moulton, V. (2018). The UEA sRNA Workbench (version 4.4): a comprehensive suite of tools for analyzing miRNAs and sRNAs. Bioinformatics 34: 3382–3384.

Sun, Q. and Zhou, D.-X. (2008). Rice jmjC domain-containing gene JMJ706 encodes H3K9 demethylase required for floral organ development. Proc. Natl. Acad. Sci. U. S. A. 105: 13679–13684.

Swetha, C., Basu, D., Pachamuthu, K., Tirumalai, V., Nair, A., Prasad, M., and Shivaprasad, P.V. (2018). Major domestication-related phenotypes in Indica rice are due to loss of miRNA-mediated laccase silencing. Plant Cell 30: 2649–2662.

Tang, J., Mei, E., He, M., Bu, Q., and Tian, X. (2022). Functions of OsWRKY24, OsWRKY70 and OsWRKY53 in regulating grain size in rice. Planta 255: 92.

Tirumalai, V., Prasad, M., and Shivaprasad, P.V. (2020). RNA blot analysis for the detection and quantification of plant MicroRNAs. J. Vis. Exp.

Tiwari, G.J., Liu, Q., Shreshtha, P., Li, Z., and Rahman, S. (2016). RNAi-mediated down-regulation of the expression of OsFAD2-1: effect on lipid accumulation and expression of lipid biosynthetic genes in the rice grain. BMC Plant Biol. 16.

Tonosaki, K. et al. (2021). Mutation of the imprinted gene OsEMF2a induces autonomous endosperm development and delayed cellularization in rice. Plant Cell 33: 85–103.

Trapnell, C., Roberts, A., Goff, L., Pertea, G., Kim, D., Kelley, D.R., Pimentel, H., Salzberg, S.L., Rinn, J.L., and Pachter, L. (2012). Differential gene and transcript expression analysis of RNA-seq experiments with TopHat and Cufflinks. Nat. Protoc. 7: 562–578.

Vivek Hari Sundar, G. and Shivaprasad, P.V. (2022). Investigation of transposon DNA methylation and copy number variation in plants using Southern hybridisation. Bio Protoc. 12: e4432.

Vu, T.M., Nakamura, M., Calarco, J.P., Susaki, D., Lim, P.Q., Kinoshita, T., Higashiyama, T., Martienssen, R.A., and Berger, F. (2013). RNA-directed DNA methylation regulates parental genomic imprinting at several loci in Arabidopsis. Development 140: 2953–2960.

Wada, H., Hatakeyama, Y., Onda, Y., Nonami, H., Nakashima, T., Erra-Balsells, R., Morita, S., Hiraoka, K., Tanaka, F., and Nakano, H. (2019). Multiple strategies for heat adaptation to prevent chalkiness in the rice endosperm. J. Exp. Bot. 70: 1299– 1311.

Wang, G., Wang, G., Zhang, X., Wang, F., and Song, R. (2012). Isolation of high quality RNA from cereal seeds containing high levels of starch. Phytochem. Anal. 23: 159– 163.

Wang, L., Zheng, K., Zeng, L., Xu, D., Zhu, T., Yin, Y., Zhan, H., Wu, Y., and Yang, D.-L. (2022). Reinforcement of CHH methylation through RNA-directed DNA methylation ensures sexual reproduction in rice. Plant Physiol. 188: 1189–1209.

Wang, Z., Butel, N., Santos-González, J., Borges, F., Yi, J., Martienssen, R.A., Martinez, G., and Köhler, C. (2020). Polymerase IV plays a crucial role in pollen development in Capsella. Plant Cell 32: 950–966.

Warthmann, N., Chen, H., Ossowski, S., Weigel, D., and Hervé, P. (2008). Highly specific gene silencing by artificial miRNAs in rice. PLoS One 3: e1829.

Waters, A.J., Bilinski, P., Eichten, S.R., Vaughn, M.W., Ross-Ibarra, J., Gehring, M., and Springer, N.M. (2013). Comprehensive analysis of imprinted genes in maize reveals allelic variation for imprinting and limited conservation with other species. Proc. Natl. Acad. Sci. U. S. A. 110: 19639–19644.

Waters, A.J., Makarevitch, I., Eichten, S.R., Swanson-Wagner, R.A., Yeh, C.-T., Xu, W., Schnable, P.S., Vaughn, M.W., Gehring, M., and Springer, N.M. (2011). Parent-of-origin effects on gene expression and DNA methylation in the maize endosperm. Plant Cell 23: 4221–4233.

Wickham, H. (2016). Ggplot2 2nd ed. (Springer International Publishing: Cham, Switzerland).

Williams, B.P., Pignatta, D., Henikoff, S., and Gehring, M. (2015). Methylation-sensitive expression of a DNA demethylase gene serves as an epigenetic rheostat. PLoS Genet. 11: e1005142.

Wolff, P., Jiang, H., Wang, G., Santos-González, J., and Köhler, C. (2015). Paternally expressed imprinted genes establish postzygotic hybridization barriers in Arabidopsis thaliana. Elife 4.

Wu, L., Zhang, Q., Zhou, H., Ni, F., Wu, X., and Qi, Y. (2009). Rice MicroRNA effector complexes and targets. Plant Cell 21: 3421–3435.

Xiao, W., Gehring, M., Choi, Y., Margossian, L., Pu, H., Harada, J.J., Goldberg, R.B., Pennell, R.I., and Fischer, R.L. (2003). Imprinting of the MEA Polycomb gene is controlled by antagonism between MET1 methyltransferase and DME glycosylase. Dev. Cell 5: 891–901.

Xie, K. and Yang, Y. (2013). RNA-guided genome editing in plants using a CRISPR-Cas system. Mol. Plant 6: 1975–1983.

Xin, M., Yang, R., Yao, Y., Ma, C., Peng, H., Sun, Q., Wang, X., and Ni, Z. (2014). Dynamic parent-of-origin effects on small interfering RNA expression in the developing maize endosperm. BMC Plant Biol. 14: 192.

Xu, L., Yuan, K., Yuan, M., Meng, X., Chen, M., Wu, J., Li, J., and Qi, Y. (2020). Regulation of rice tillering by RNA-directed DNA methylation at miniature inverted-repeat transposable elements. Mol. Plant 13: 851–863.

Xu, R. et al. (2018). Control of grain size and weight by the OsMKKK10-OsMKK4-OsMAPK6 signaling pathway in rice. Mol. Plant 11: 860–873.

Yan, D., Duermeyer, L., Leoveanu, C., and Nambara, E. (2014). The functions of the endosperm during seed germination. Plant Cell Physiol. 55: 1521–1533.

Yang, D.-L. et al. (2018a). Four putative SWI2/SNF2 chromatin remodelers have dual roles in regulating DNA methylation in Arabidopsis. Cell Discov. 4: 55.

Yang, J., Luo, D., Yang, B., Frommer, W.B., and Eom, J.-S. (2018b). SWEET11 and 15 as key players in seed filling in rice. New Phytol. 218: 604–615.

Yao, M., Ai, T.-B., Mao, Q., Chen, F., Li, F.-S., and Tang, L. (2018). Downregulation of OsAGO17 by artificial microRNA causes pollen abortion resulting in the reduction of grain yield in rice. Electron. J. Biotechnol.

Yin, L.-L. and Xue, H.-W. (2012). The MADS29 transcription factor regulates the degradation of the nucellus and the nucellar projection during rice seed development. Plant Cell 24: 1049–1065.

Yuan, J., Chen, S., Jiao, W., Wang, L., Wang, L., Ye, W., Lu, J., Hong, D., You, S., Cheng, Z., Yang, D., and Chen, Z.J. (2017). Both maternally and paternally imprinted genes regulate seed development in rice. New Phytol. 216: 373–387.

Zemach, A., Kim, M.Y., Silva, P., Rodrigues, J.A., Dotson, B., Brooks, M.D., and Zilberman, D. (2010). Local DNA hypomethylation activates genes in rice endosperm. Proc. Natl. Acad. Sci. U. S. A. 107: 18729–18734.

Zhang, H. et al. (2013). DTF1 is a core component of RNA-directed DNA methylation and may assist in the recruitment of Pol IV. Proc. Natl. Acad. Sci. U. S. A. 110: 8290– 8295.

Zhang, H., Xu, H., Feng, M., and Zhu, Y. (2018). Suppression of OsMADS7 in rice endosperm stabilizes amylose content under high temperature stress. Plant Biotechnol. J. 16: 18–26.

Zhang, J., Nallamilli, B.R., Mujahid, H., and Peng, Z. (2010). OsMADS6 plays an essential role in endosperm nutrient accumulation and is subject to epigenetic regulation in rice (Oryza sativa). Plant J. 64: 604–617.

Zhang, L. et al. (2012). Identification and characterization of an epi-allele of FIE1 reveals a regulatory linkage between two epigenetic marks in rice. Plant Cell 24: 4407–4421.

Zheng, K., Wang, L., Zeng, L., Xu, D., Guo, Z., Gao, X., and Yang, D.-L. (2021). The effect of RNA polymerase V on 24-nt siRNA accumulation depends on DNA methylation contexts and histone modifications in rice. Proc. Natl. Acad. Sci. U. S. A. 118: e2100709118.

Zhou, M., Coruh, C., Xu, G., Martins, L.M., Bourbousse, C., Lambolez, A., and Law, J.A. (2022). The CLASSY family controls tissue-specific DNA methylation patterns in Arabidopsis. Nat. Commun. 13: 244.

Zhou, M., Palanca, A.M.S., and Law, J.A. (2018). Locus-specific control of the de novo DNA methylation pathway in Arabidopsis by the CLASSY family. Nat. Genet. 50: 865–873.

Zhu, H., Xie, W., Xu, D., Miki, D., Tang, K., Huang, C.-F., and Zhu, J.-K. (2018). DNA demethylase ROS1 negatively regulates the imprinting of DOGL4 and seed dormancy in Arabidopsis thaliana. Proc. Natl. Acad. Sci. U. S. A. 115: E9962–E9970.

Zilberman, D., Cao, X., and Jacobsen, S.E. (2003). ARGONAUTE4 control of locus-specific siRNA accumulation and DNA and histone methylation. Science 299: 716– 719.

