## Supplemental Figures and Tables for "Upstream regulator of genomic imprinting in rice endosperm is a small RNA-associated chromatin remodeler"

### Supplemental information

This manuscript has 9 supplemental figures, 4 supplemental tables and 9 supplemental datasets.

### Supplemental Figures

Supplemental Figure S1. Transcriptome analysis of endosperm and embryo tissues.

Supplemental Figure S2. *OsCLSY3* is an endosperm-preferred gene among monocots.

Supplemental Figure S3. *KO* transgenic plants show reproductive abnormalities.

Supplemental Figure S4. Validation of *OsCLSY3* transgenic lines.

Supplemental Figure S5. Endosperm sRNAs are globally reduced in clsy3-kd.

Supplemental Figure S6. The RdDM pathway regulates expression of siren loci in rice.

Supplemental Figure S7. CLSY3-dependent sRNAs regulate protein coding genes.

Supplemental Figure S8. The RdDM pathway controls imprinted sRNA loci in rice endosperm.

Supplemental Figure S9. DNA methylation levels in imprinted genes of embryo and endosperm.

### Supplemental Tables

Supplemental Table S1. Details of high-throughput genomics data generated in this study.

Supplemental Table S2. Details of high-throughput genomics data obtained from publicly available datasets.

Supplemental Table S3. List of oligos and probes used in this study.

Supplemental Table S4. Sequences of DRD1 family proteins used in this study.

### Supplemental datasets

Supplemental dataset S1. List of $Log_2$ 2.0-fold change upregulated genes mature endosperm and embryo (endosperm-preferred genes).

Supplemental dataset S2. List of $Log_2$ 2.0-fold change downregulated genes mature endosperm and embryo (embryo-preferred genes).

Supplemental dataset S3. List of epigenetic genes and all published imprinted genes in rice.

Supplemental dataset S4. List of all Shortstack 23-24 nt sRNA loci in WT endosperm.

Supplemental dataset S5. List of all Shortstack 23-24 nt sRNA loci in clsy3-kd endosperm.

Supplemental dataset S6. List of CLSY3-dependent 23-24 nt sRNA loci.

Supplemental dataset S7. List of published rice siren loci with their normalized expression.

Supplemental dataset S8. List of $log_2$1.5-fold change upregulated genes in clsy3-kd endosperm.

Supplemental dataset S9. List of $log_2$1.5-fold change downregulated genes in clsy3-kd endosperm.

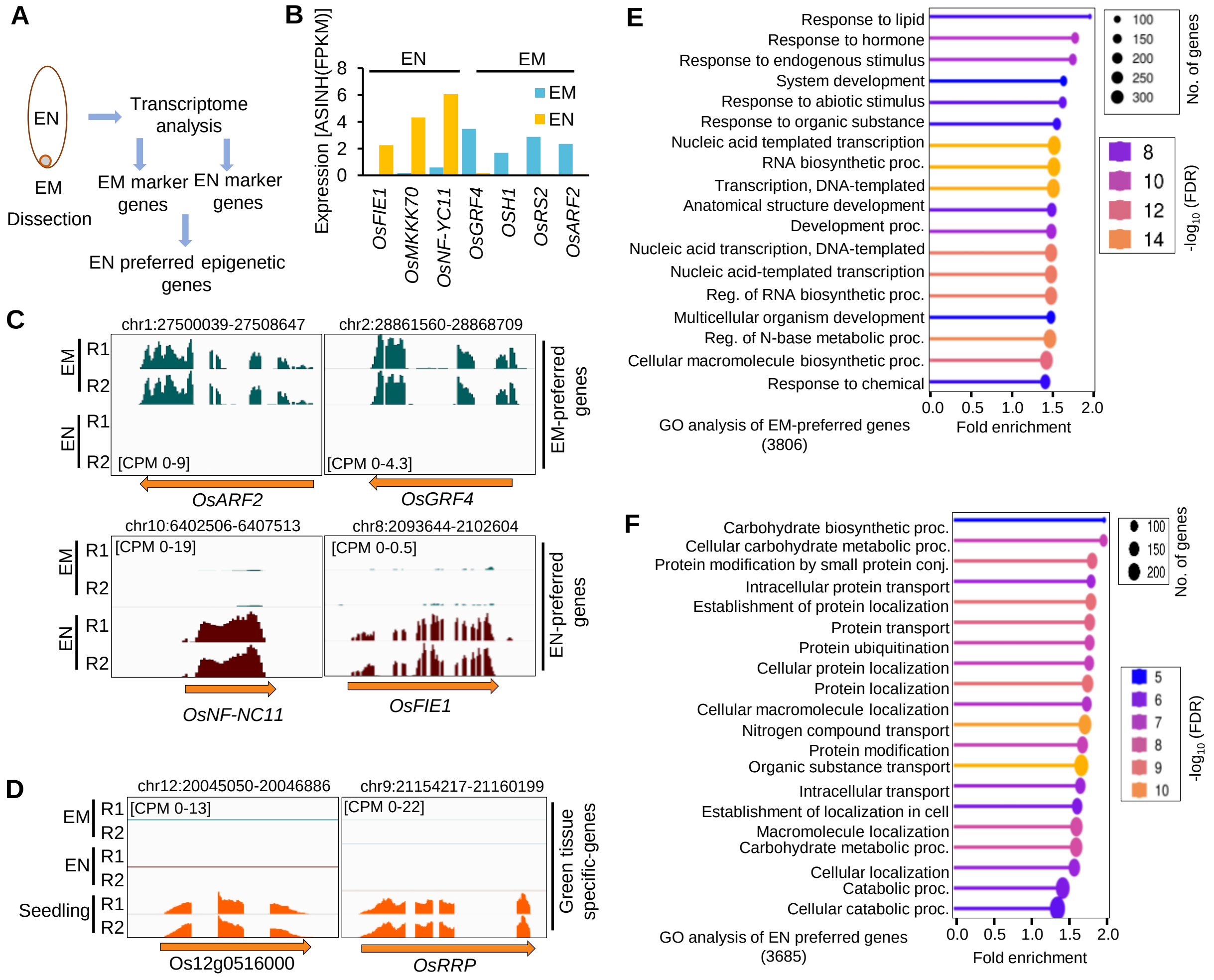


**Supplemental Figure S1. Transcriptome analysis of endosperm and embryo tissues.**
(A) Schematic showing tissue collection for transcriptome analysis. (B) Barplots showing expression of embryo (EM) and endosperm (EN)-specific marker genes in transcriptome analysis. (C) IGV screenshots showing expression of known EM- and EN-specific marker genes in transcriptome. (D) IGV screenshots showing expression of non-seed expressing genes in transcriptome. (E) GO analysis of EM-preferred genes. (F) GO analysis of EN-preferred genes.

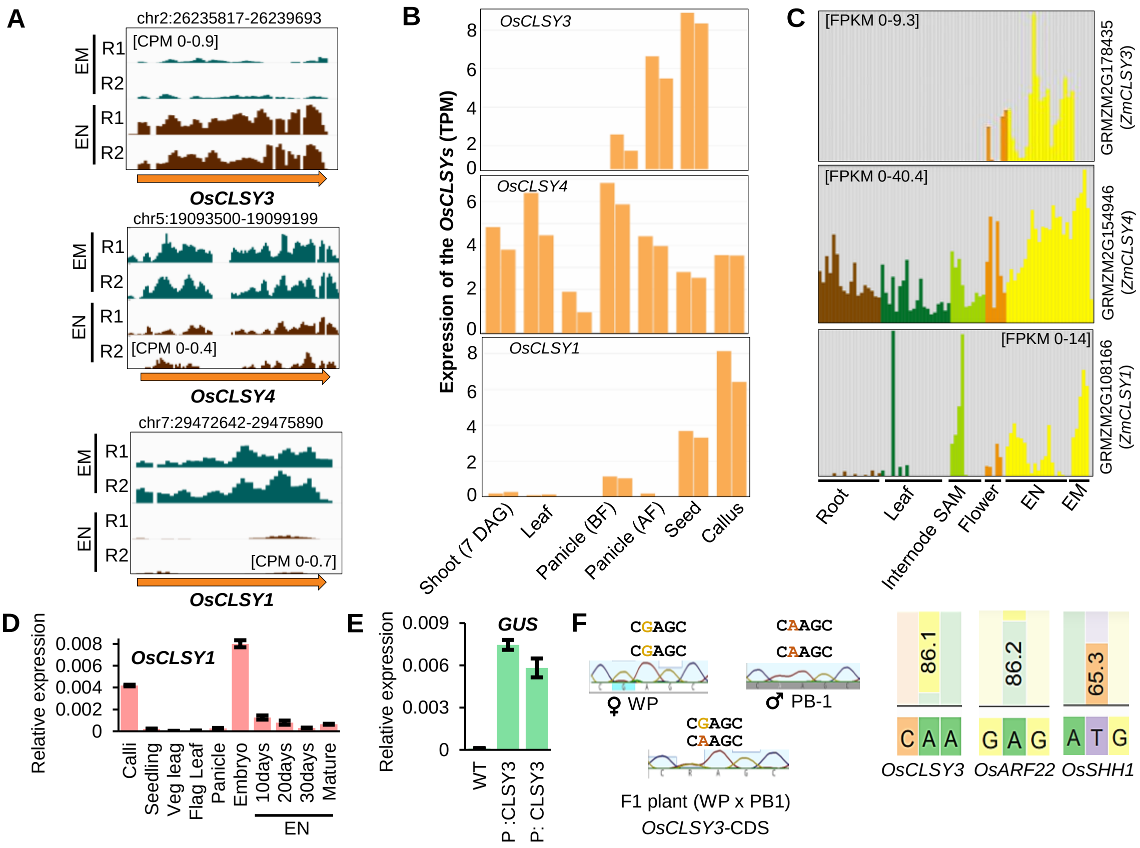


**Supplemental Figure S2. *OsCLSY3* is an endosperm-preferred gene among monocots.**
(A) IGV screenshots showing expression of *OsCLSYs* in EN and EM. (B) Barplots representing expression of *OsCLSYs* across different tissues (The Rice Annotation Project- https://rapdb.dna.affrc.go.jp/). (C) Barplots representing expression of maize *CLSYs* across different tissues (MaizeDB-https://www.maizegdb.org/). (D) Barplot showing expression of *OsCLSY1* across tissues. *OsActin* served as control. Error bar- standard error (SE). (E) Barplot showing expression of GUS in the EN of P:CLSY3:GUS transgenic lines. *OsActin* served as control. Error bar-SE. (F) Scheme depicting validation of WP and PB1 cross and the SNPs identified in CDS of *OsCLSY3*. The stacked barplots showing observed transcript contribution of three genes in WWP EN (Mismatches were calculated in crispresso2).

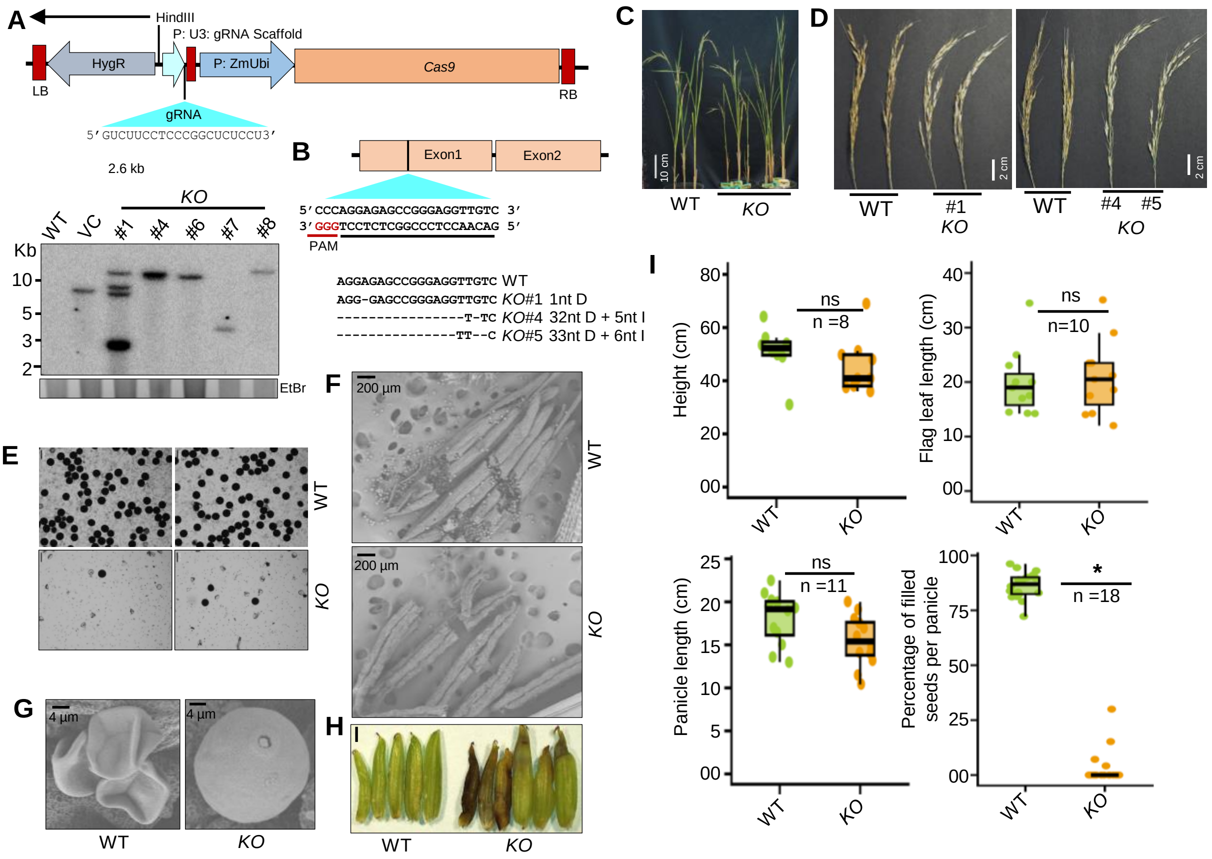


**Supplemental Figure S3. *KO* transgenic plants show reproductive abnormalities.**
(A) Vector map of *KO* construct and DNA blots showing the T-DNA junction corresponding to digestion of DNA with mentioned restriction sites. Minimum lengths of junction fragments sizes are mentioned in the map. Probe (*hph*-1kb) region is shown. (B) Schematic showing editing in guide RNA targeted region of *OsCLSY3* gene in *KO* lines. (C) Image showing *KO* plants with equally grown WT plants (WT-Wild type). (D) Panicle morphology of *KO* lines. (E) Pollen viability assay of *KO* pollen (*KO#4* and *KO#5* used for the assay, Scale bar(SB)-50µm). (F) Anther and pollen morphology of *KO* under the electron microscope (*KO#4).* (G) Detailed pollen morphology of *KO* under the electron microscope. (H) Image showing EN morphology of *KO (KO#1).* (I) Box plots representing various agronomic phenotypes measured in WT and KO plants. *-significant and (ns)-non-significant. (two-tailed Student's *t*-test).

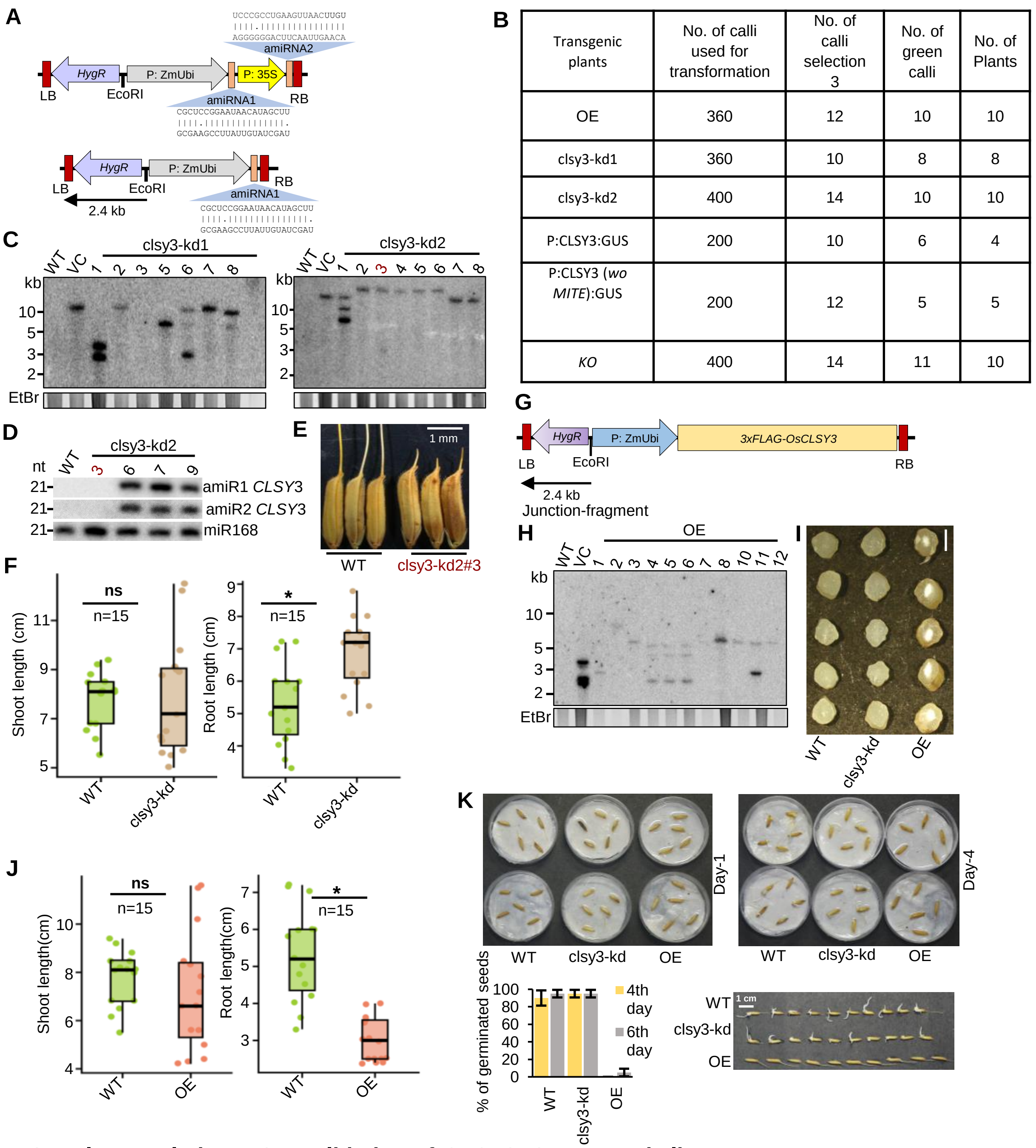


| Transgenic plants | No. of calli used for transformation | No. of calli selection 3 | No. of green calli | No. of Plants |
|---|---|---|---|---|
| OE | 360 | 12 | 10 | 10 |
| clsy3-kd1 | 360 | 10 | 8 | 8 |
| clsy3-kd2 | 400 | 14 | 10 | 10 |
| P:CLSY3:GUS | 200 | 10 | 6 | 4 |
| P:CLSY3 (*wo MITE*):GUS | 200 | 12 | 5 | 5 |
| *KO* | 400 | 14 | 11 | 10 |

**Supplemental Figure S4. Validation of *OsCLSY3* transgenic lines.**
(A) Vector maps of amiRNA constructs to target *OsCLSY3*. (B) Table showing approximate number of calli used and transgenic plants obtained in this study. (C) DNA blots showing the T-DNA junction corresponding to digestion of DNA with mentioned restriction sites. Minimum lengths of junction fragments sizes are mentioned in the map. Probe (hph-1kb) region is shown. The line clsy3-kd2 #3 (brown) had proper T-DNA insertion. (D) Northern blots showing amiR expression in clsy3-kd plants. The clsy3-kd2 #3 (marked in brown), did not express amiRs. (E) Image of clsy3-kd2 #3 seeds. (F) Box plots showing the shoot and root lengths of 12 d old clsy3-kd seedlings. (G) Vector map of OE construct. (H) DNA blots showing the T-DNA junction corresponding to digestion of DNA with mentioned restriction sites. (I) Cross sections of dry clsy3-kd and OE EN (SB-1 mm). (J) Box plots showing the shoot and root lengths of 12 d old OE seedlings. In box plots, (ns)-non-significant and *-significant (two-tailed Student's *t*-test). (K) The image showing germination of *OsCLSY3* genotypes. The barplot showing percentage of germinated seeds in *OsCLSY3* genotypes.

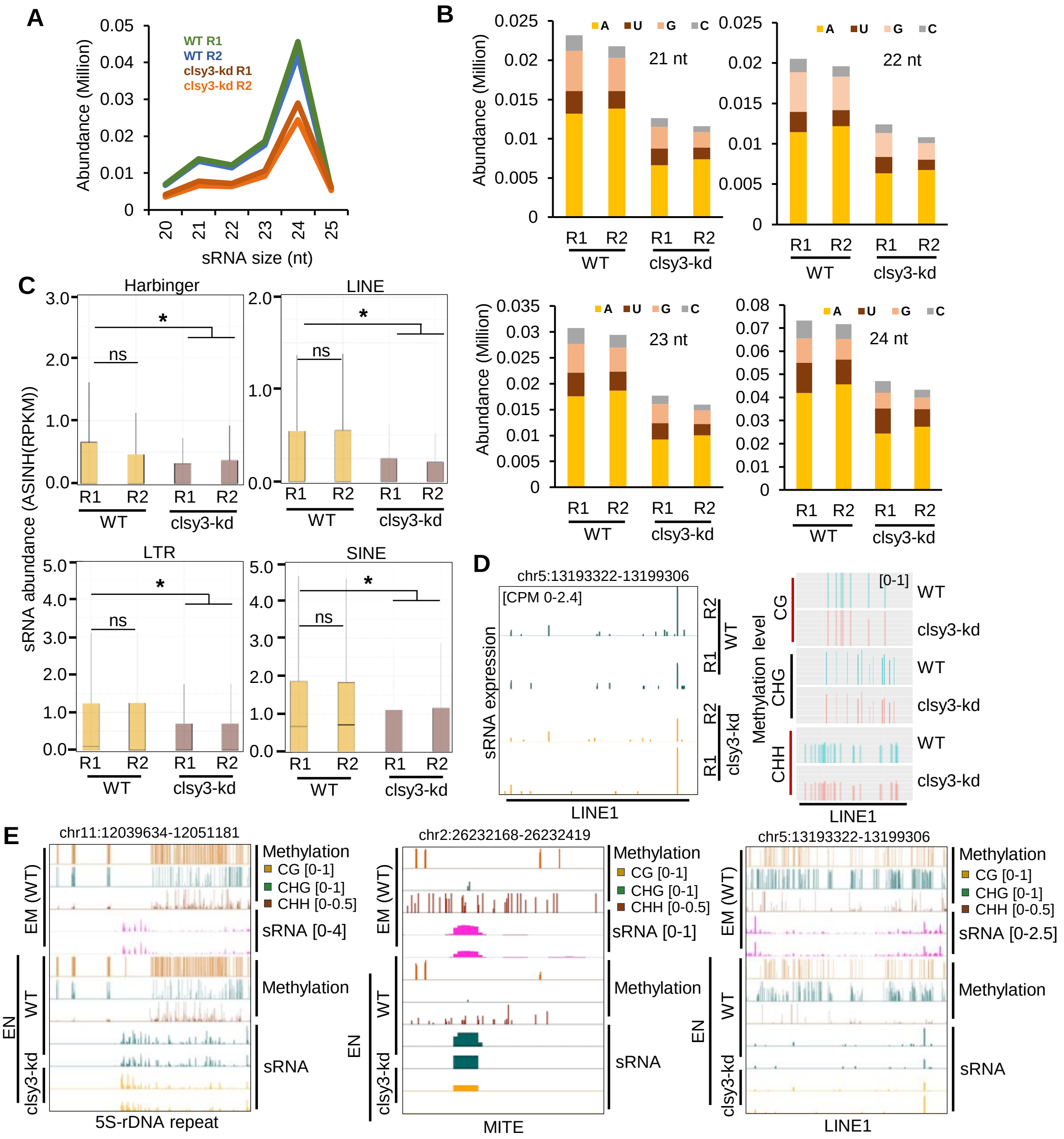


**Supplemental Figure S5. Endosperm sRNAs are globally reduced in clsy3-kd.**
(A) Plot showing mapped sRNA abundance (20-25 nt) in EN. (B) Stacked barplot showing abundance of first 5' nucleotide of mapped sRNAs in WT and clsy3-kd EN. (C) Boxplots representing normalised sRNA reads across different class II transposons in WT and clsy3-kd EN. *-significant and ns-non-significant (Wilcoxon test $p < 0.01$). (D) IGV screenshots showing sRNA and methylation status of LINE1 transposon in clsy3-kd EN. (E) IGV screenshots depicting the sRNA and methylation levels of TEs in WT EN and EM selected for BS-PCR.

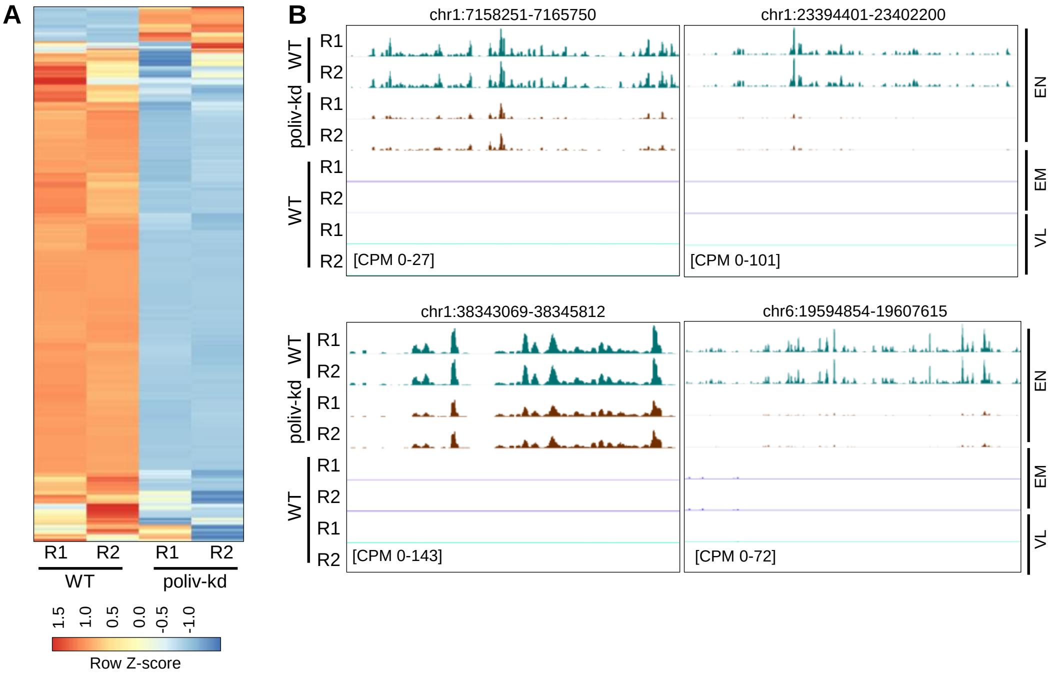


**Supplemental figure 6. The RdDM pathway regulates expression of siren loci in rice.**
(A) Heatmap showing expression of siren loci generated sRNAs (23-24 nt) in poliv-kd EN. Row Z-score was plotted (797 loci). (B) IGV screenshot representing sRNA levels (23-24 nt) in some selected siren loci in poliv-kd EN.

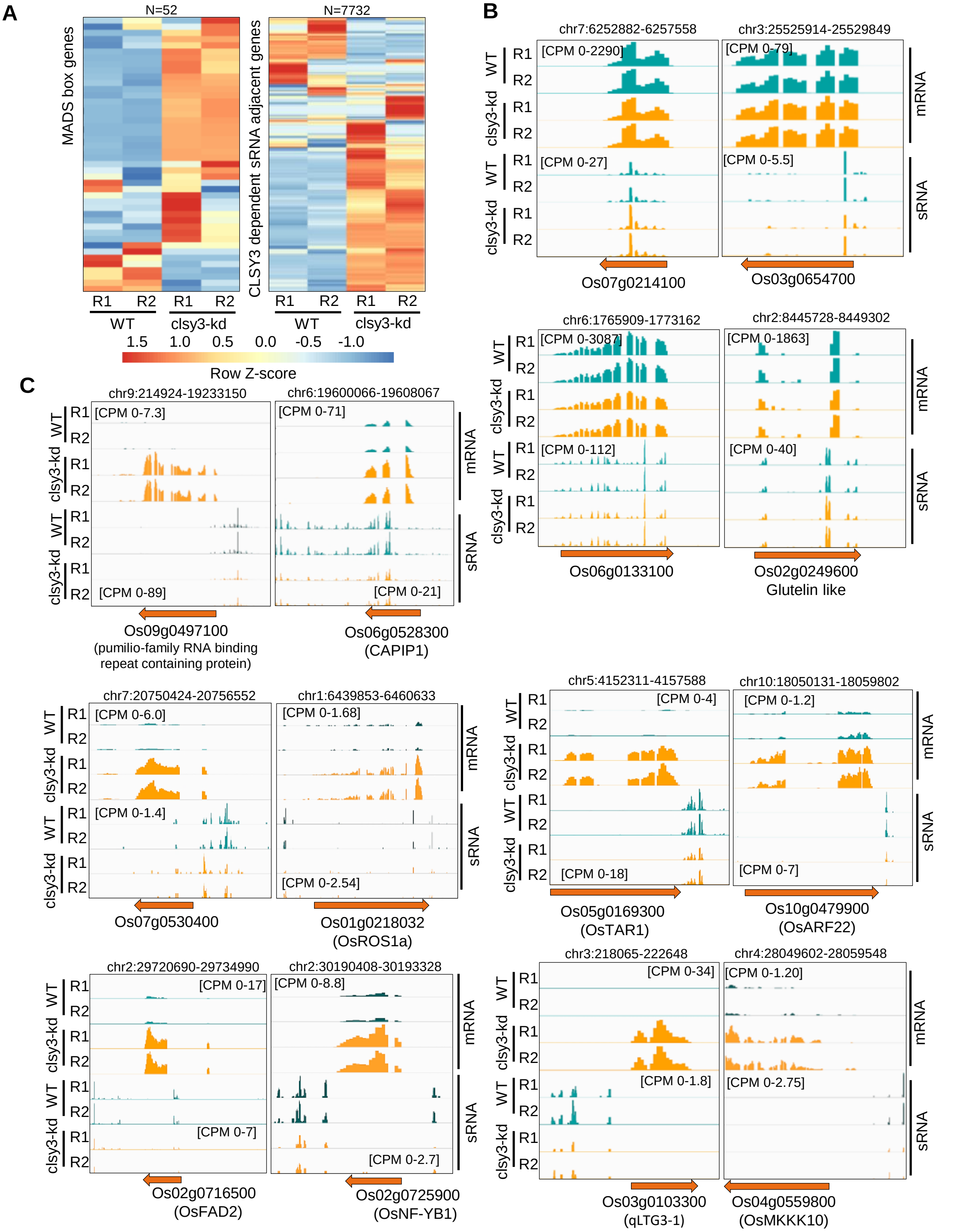


**Supplemental Figure S7. CLSY3-dependent sRNAs regulate protein coding genes.** (A) Heatmap showing MADS box and CLSY3-dependent sRNA adjacent genes in clsy3-kd EN. Row Z-score was plotted. (B) IGV screenshots showing four control genes in which sRNA and mRNA levels unaltered in clsy3-kd. (C) IGV screenshots showing expression of few selected seed development and yield related genes in clsy3-kd EN.

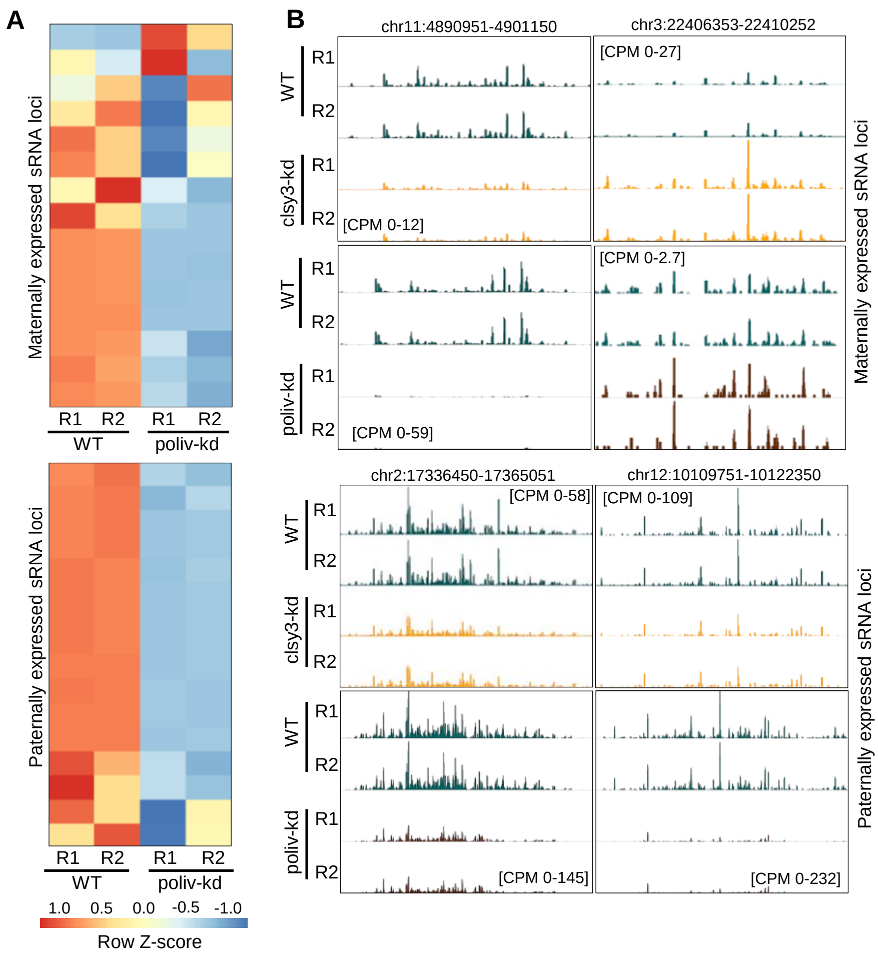


**Supplemental Figure S8. The RdDM pathway controls imprinted sRNA loci in rice endosperm.** (A) Heatmap showing expression of imprinted sRNAs in poliv-kd EN. Row Z-score was plotted. (B) IGV screenshots representing expression of imprinted sRNAs (23-24 nt) in poliv-kd and clsy3-kd EN.

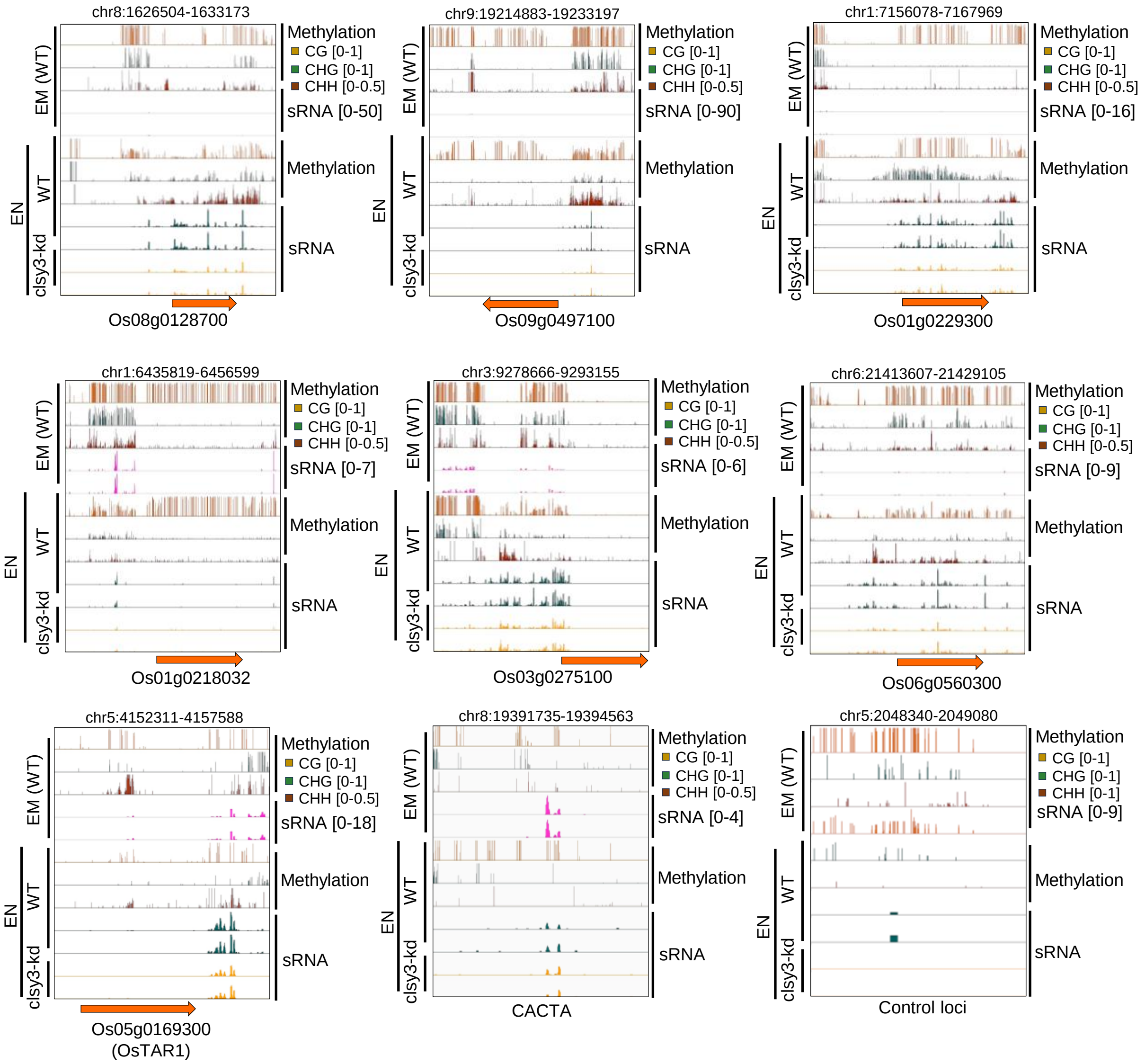


**Supplemental Figure S9. DNA methylation levels in imprinted genes of embryo and endosperm**
IGV screen shots showing CLSY3-dependent EN-specific hypermethylation at proximity to imprinted genes.

Supplemental Table S1: Details of high-throughput genomics data generated in this study

| Sl. No | Dataset type | Genotype | Replicate | Source tissue | Total number of mapped reads obtained | Sequencing mode |
|---|---|---|---|---|---|---|
| 1 | Small RNA-seq | WT | Rep1 | 20 days Endosperm | 39570183 | Single end 50bp |
| 2 | Small RNA-seq | WT | Rep2 | 20 days Endosperm | 29825291 | Single end 50bp |
| 3 | Small RNA-seq | clsy3-kd | Rep1 | 20 days Endosperm | 41731266 | Single end 50bp |
| 4 | Small RNA-seq | clsy3-kd | Rep2 | 20 days Endosperm | 26694590 | Single end 50bp |
| 5 | Small RNA-seq | WT | Rep1 | 25 days Embryo | 34720489 | Single end 50bp |
| 6 | Small RNA-seq | WT | Rep2 | 25days Embryo | 29840669 | Single end 50bp |
| 5 | RNA-seq | WT | Rep1 | 20 days Endosperm | 12941016 | Paired end 100bp |
| 6 | RNA-seq | WT | Rep2 | 20 days Endosperm | 14608494 | Paired end 100bp |
| 7 | RNA-seq | clsy3-kd | Rep1 | 20 days Endosperm | 13857932 | Paired end 100bp |
| 8 | RNA-seq | clsy3-kd | Rep2 | 20 days Endosperm | 13461363 | Paired end 100bp |
| 9 | RNA-seq | Embryo | Rep1 | 25 days Embryo | 44252442 | Paired end 100bp |
| 10 | RNA-seq | Embryo | Rep2 | 25 days Embryo | 32808865 | Paired end 100bp |
| 11 | RNA-seq | Endosperm | Rep1 | 25 days Endosperm | 35622255 | Paired end 100bp |
| 12 | RNA-seq | Endosperm | Rep2 | 25 days Endosperm | 30799873 | Paired end 100bp |
| 13 | RNA-seq | Endosperm | Rep1 | 15 days Endosperm | 37858782 | Paired end 100bp |
| 14 | RNA-seq | Endosperm | Rep2 | 15 days Endosperm | 33080996 | Paired end 100bp |
| 15 | Targeted bisulfite | WT Leaf | NA | 60 days Leaf | 1031542 paired | Paired end 100bp |
| 16 | Targeted bisulfite | WT Panicle | NA | Mature panicle (12-15 cm) | 1074742 paired | Paired end 100bp |
| 17 | Targeted bisulfite | WT Endosperm | NA | 20 days Endosperm | 1166935 paired | Paired end 100bp |

| | | | | | | |
|---|---|---|---|---|---|---|
| 18 | Targeted bisulfite | Untreated DNA | NA | Untreated Leaf DNA | 925634 paired | Paired end 100bp |
| 19 | Targeted bisulfite | WT Endosperm | NA | 20 days Endosperm | 991160 paired | Paired end 100bp |
| 20 | Targeted bisulfite | clsy3-kd Endosperm | NA | 20 days Endosperm | 961552 paired | Paired end 100bp |

Supplemental Table S2: Details of high-throughput genomics data obtained from publicly available datasets

| Sl. No | Dataset type | Genotype | Source tissue | SRA number | GSE number | Reference |
|---|---|---|---|---|---|---|
| 1 | RNAseq | WT | pre-emerged panicle | SRX11493038 | GSE180457 | (Hari Sundar G et al., 2023) |
| 2 | RNAseq | WT | pre-emerged panicle | SRX11493039 | GSE180457 | (Hari Sundar G et al., 2023) |
| 3 | RNAseq | nrpd1_kd | pre-emerged panicle | SRX11493040 | GSE180457 | (Hari Sundar G et al., 2023) |
| 4 | RNAseq | nrpd1_kd | pre-emerged panicle | SRX11493041 | GSE180457 | (Hari Sundar G et al., 2023) |
| 5 | Small RNAseq | WT | Endosperm (16-20 days after anthesis) | SRX11504321 | GSE180457 | (Hari Sundar G et al., 2023) |
| 6 | Small RNAseq | WT | Endosperm (16-20 days after anthesis) | SRX11504322 | GSE180457 | (Hari Sundar G et al., 2023) |
| 7 | Small RNAseq | nrpd1-kd | Endosperm (16-20 days after anthesis) | SRX11504323 | GSE180457 | (Hari Sundar G et al., 2023) |
| 8 | Small RNAseq | nrpd1-kd | Endosperm (16-20 days after anthesis) | SRX11504324 | GSE180457 | (Hari Sundar G et al., 2023) |
| 9 | Small RNAseq | WT | pre-emerged panicle | SRX11504325 | GSE180457 | (Hari Sundar G et al., 2023) |
| 10 | Small RNAseq | WT | pre-emerged | SRX11504326 | GSE180457 | (Hari Sundar G et al., 2023) |
| 11 | Small RNAseq | nrpd1-kd | panicle pre-emerged | SRX11504327 | GSE180457 | (Hari Sundar G et al., 2023) |
| 12 | Small RNAseq | nrpd1-kd | panicle pre-emerged panicle | SRX11504328 | GSE180457 | (Hari Sundar G et al., 2023) |

| | | | | | | |
|---|---|---|---|---|---|---|
| 13 | Small RNA-seq | WT (Nipponbare) | Seedlings | SRX5724235 | GSE130166 | (Wang et al., 2022) |
| 14 | Small RNA-seq | WT (Nipponbare) | Seedlings | SRX5724236 | GSE130166 | (Wang et al., 2022) |
| 15 | Small RNA-seq | rdr2-6 | Seedlings | SRX5724237 | GSE130166 | (Wang et al., 2022) |
| 16 | Small RNA-seq | WT (Nipponbare) | Panicle | SRX5724233 | GSE130166 | (Wang et al., 2022) |
| 17 | Small RNA-seq | rdr2-6 | Panicle | SRX5724234 | GSE130166 | (Wang et al., 2022) |
| 18 | Small RNA-seq | nrpd1 | Seedlings | SRX9211921 | GSE158709 | (Zheng et al., 2021) |
| 19 | Small RNA-seq | nrpe1 | Seedlings | SRX9211922 | GSE158709 | (Zheng et al., 2021) |
| 20 | Small RNA-seq | nrpe1 | Seedlings | SRX9211920 | GSE158709 | (Zheng et al., 2021) |
| 21 | Small RNA-seq | WT rep1 | Leaf | SRX11930857 | PRJNA758109 | (Chakraborty et al., 2022) |
| 22 | Small RNA-seq | WT rep2 | Leaf | SRX11930858 | PRJNA758109 | (Chakraborty et al., 2022) |
| 23 | Small RNA-seq | WT rep3 | Leaf | SRX11930859 | PRJNA758109 | (Chakraborty et al., 2022) |
| 24 | Small RNA-seq | nrp(d/e)2 Rep1 | Leaf | SRX11930854 | PRJNA758109 | (Chakraborty et al., 2022) |
| 25 | Small RNA-seq | nrp(d/e)2 Rep1 | Leaf | SRX11930855 | PRJNA758109 | (Chakraborty et al., 2022) |
| 26 | Small RNA-seq | nrp(d/e)2 Rep1 | Leaf | SRX11930856 | PRJNA758109 | (Chakraborty et al., 2022) |
| 27 | Bisulfite-Seq | WT (IR64) | Endosperm | SRR8937051 | GSE130122 | (Rodrigues et al., 2021) |
| 28 | Bisulfite-Seq | WT (IR64) | Embryo | SRR8937043 | GSE130122 | (Rodrigues et al., 2021) |
| 29 | Bisulfite-Seq | WT | 70-days leaf | SRX6976659 | GSE138705 | (Hu et al., 2022) |
| 30 | Bisulfite-Seq | oscmt2/cmt3a | 70-days leaf | SRX6976664 | GSE138705 | (Hu et al., 2022) |
| 31 | Bisulfite-Seq | osdrm2 | 70-days leaf | SRX6976666 | GSE138705 | (Hu et al., 2022) |
| 32 | Bisulfite-Seq | osdrm2 | 70-days leaf | SRX6976667 | GSE138705 | (Hu et al., 2022) |
| 33 | Bisulfite-Seq | osdrm1a/drm1b | 70-days leaf | SRX6976671 | GSE138705 | (Hu et al., 2022) |
| 34 | Bisulfite-Seq | osdrm2/drm3 | 70-days leaf | SRX6976672 | GSE138705 | (Hu et al., 2022) |
| 35 | Bisulfite-Seq | oscmt2/cmt3a/cmt3b/drm2/drm3 | 70-days leaf | SRX6976677 | GSE138705 | (Hu et al., 2022) |
| 36 | Bisulfite-Seq | oscmt2/cmt3a/cmt3b/drm2/drm3 | 70-days leaf | SRX6976678 | GSE138705 | (Hu et al., 2022) |
| 37 | Bisulfite-Seq | BS-seq_Nip_se_rep1 | Seedlings | SRX5724238 | GSE130168 | (Wang et al., 2022) |

| | | | | | | |
|---|---|---|---|---|---|---|
| 38 | Bisulfite-Seq | BS-seq_Nip_se_rep2 | Seedlings | SRX5724239 | GSE130168 | (Wang et al., 2022) |
| 39 | Bisulfite-Seq | BS-seq_osrdr2-6_se | Seedlings | SRX5724232 | GSE130168 | (Wang et al., 2022) |
| 40 | AGO1a-IP | WT | Seedling | SRX014804 | GSE18250 | (Wu et al., 2009) |
| 41 | AGO1b-IP | WT | Seedling | SRX014805 | GSE18250 | (Wu et al., 2009) |
| 42 | AGO1c-IP | WT | Seedling | SRX014806 | GSE18250 | (Wu et al., 2009) |
| 43 | AGO4a-IP | WT | Seedling | SRX017394 | GSE20748 | (Wu et al., 2009) |
| 44 | AGO4b-IP | WT | Seedling | SRX017395 | GSE20748 | (Wu et al., 2009) |

### References


**Chakraborty, T., Trujillo, J.T., Kendall, T., and Mosher, R.A.** (2022). A null allele of the pol IV second subunit impacts stature and reproductive development in Oryza sativa. Plant J. **111**: 748–755.

**Hari Sundar G, V., Swetha, C., Basu, D., Pachamuthu, K., Raju, S., Chakraborty, T., Mosher, R.A., and Shivaprasad, P.V.** (2023). Plant polymerase IV sensitizes chromatin through histone modifications to preclude spread of silencing into protein-coding domains. Genome Res.

**Hu, D. et al.** (2022). Erratum for: Multiplex CRISPR-Cas9 editing of DNA methyltransferases in rice uncovers a class of non-CG methylation specific for GC-rich regions. Plant Cell **34**: 1416.

**Rodrigues, J.A., Hsieh, P.-H., Ruan, D., Nishimura, T., Sharma, M.K., Sharma, R., Ye, X., Nguyen, N.D., Nijjar, S., Ronald, P.C., Fischer, R.L., and Zilberman, D.** (2021). Divergence among rice cultivars reveals roles for transposition and epimutation in ongoing evolution of genomic imprinting. Proc. Natl. Acad. Sci. U. S. A. **118**: e2104445118.

**Wang, L., Zheng, K., Zeng, L., Xu, D., Zhu, T., Yin, Y., Zhan, H., Wu, Y., and Yang, D.-L.** (2022). Reinforcement of CHH methylation through RNA-directed DNA methylation ensures sexual reproduction in rice. Plant Physiol. **188**: 1189–1209.

**Wu, L., Zhang, Q., Zhou, H., Ni, F., Wu, X., and Qi, Y.** (2009). Rice MicroRNA effector complexes and targets. Plant Cell **21**: 3421–3435.

**Zheng, K., Wang, L., Zeng, L., Xu, D., Guo, Z., Gao, X., and Yang, D.-L.** (2021). The effect of RNA polymerase V on 24-nt siRNA accumulation depends on DNA methylation contexts and histone modifications in rice. Proc. Natl. Acad. Sci. U. S. A. **118**: e2100709118.

Supplemental Table S3: List of oligos and probes used in this study

| Oligo Name | Oligo ID | Oligo sequence (5' – 3') | Purpose | Reference |
|---|---|---|---|---|
| AP_OsCLSY3_F3 | 2571 | ccatctccaacgcggagcagagtagaacacc | Amplification of *OsCLSY3* from genome | This study |
| AP_OsCLSY3_R1 | 2472 | ctgttagtgagaatggagaatcttcac | Amplification of *OsCLSY3* from genome | This study |
| AP_CLSY3_EX3_RT_F2 | 3709 | ggtgtgagctatacgcttttcagg | RT-qPCR and RTPCR of CLSY3 | This study |
| AP_CLSY3_EX3_RT_R2 | 3710 | gcgaagtgtcttgcaaactttggtc | RT-qPCR and RTPCR of CLSY3 | This study |
| AP_CLSY3_Prom_F3 | 2817 | ggccagctagtactttcagaggatcttcatcg | Amplification from genome of CLSY3 promoter | This study |
| AP_CLSY3_prom_R2 | 2818 | gtcgatgatctctagcgaggggctacgtgcac | Amplification from genome of CLSY3 promoter | This study |
| pRGEB32_Seq_F | 1435 | CCGGCTCGTATGTTGTGTGG | Amplification of CLSY3 gRNA | This study |
| AP_OsCLSY3_gRNA2_R | 2602 | tataggtctcaaaacAGGAGAGCCGGGAGGTTGTCTGCCACGGATCATCTGCACAACTC | Amplification of CLSY3 gRNA | This study |
| Hyg_F | 415 | aaagcctgaactcaccgc | RT-PCR and Southern probe | This study |
| Hyg_R | 416 | ggtttccactatcggcga | RT-PCR and Southern probe | This study |
| STN_OsRAC1_F | 2563 | GCTATGTACGTCGCCATCCAGG | RT-qPCR | This study |
| STN_OsRAC1_R | 2564 | TGAGATCACGCCCAGCAAGG | RT-qPCR | This study |
| AP_CLSY4_RT_F | 3419 | gtggtggatccagtgaatgaagagttgg | RT-qPCR | This study |
| AP_CLSY4_RT_R | 3420 | cccaaaattatcaccgtcgtcatcagc | RT-qPCR | This study |
| AP_FIE1_qpcr_F | 3602 | AAAAAGATGTGGCACCGAAT | RT-qPCR | (Chen et al., 2018) |
| AP_FIE1_qpcr_R | 3603 | GGTTCACCTTCACCAGCAGT | RT-qPCR | (Chen et al., 2018) |
| qPCR_NRPD1a&b _F | 1769 | ttgcaagtattgctcaaaggatgg | RT-qPCR | (Hari Sundar G et al., 2023) |

| | | | | |
|---|---|---|---|---|
| qPCR_NRPD1a &b _R | 1770 | ccaccagcaacttctgcgataattg | RT-qPCR | (Hari Sundar G et al., 2023) |
| VHS_GUS_qRT _PCR_Fwd | 1863 | CCCTTACGCTGAAGAGAT GC | RT-qPCR | This study |
| VHS_GUS_qRT _PCR_ rev | 1864 | TTCGTTGGCAATACTCCA CA | RT-qPCR | This study |
| AP_CHOP_PC R_1A_F | 4820 | CATCCACTGTCCAGATGA AGTCAG | RT-CHOP PCR | This study |
| AP_CHOP_PC R_1A_R | 4821 | ACTTCGATAATGGTAAAA CGCCTCC | RT-CHOP PCR | This study |
| AP_CHOP_PC R_3B_F | 4826 | AACCATTTCCATCCCATC CCAACTCG | RT-CHOP PCR | This study |
| AP_CHOP_PC R_3B_R | 4827 | CTAGCGATTCTACTATAT TTTAAAGACACATATTC | RT-CHOP PCR | This study |
| AP_CHOP_PC R_6A_F | 4836 | CCTCCGACATGTACTTGC GGTGCTCC | RT-CHOP PCR | This study |
| AP_CHOP_PC R_6A_R | 4837 | TCTCCCTCCCCATGTGAG TGATCCTGC | RT-CHOP PCR | This study |
| AP_CHOP_PC R_2B_F | 4824 | CTGCAGAAAGGCACCAA CCTCAGAG | RT-CHOP PCR | This study |
| AP_CHOP_PC R_2B_R | 4825 | AGGACAACTTTGTTCTCC ATTATGATC | RT-CHOP PCR | This study |
| AP_CHOP_PC R_5A_F | 4834 | GTCGACATGGAAAGGAG CTTGAGATACCC | RT-CHOP PCR | This study |
| AP_CHOP_PC R_5A_R | 4835 | CGACTCTTGTGTTCCCTT ACACATCATAGC | RT-CHOP PCR | This study |
| AP_CHOP_PC R_8A_F | 4842 | CAATGGTGAGAAGATGC ATTTGATGATGC | RT-CHOP PCR | This study |
| AP_CHOP_PC R_8A_R | 4843 | CCACGGCATCCATTCTCT CAGATGG | RT-CHOP PCR | This study |
| 5A_F_VHS_Os ChOP | 3997 | AAATGTACTTACCCTCAC GAGCAT | RT-CHOP PCR | (Chakraborty et al., 2022) |
| 5A_R_VHS_Os ChOP | 3998 | AATTTTACTACACTTGCG AGCCAA | RT-CHOP PCR | (Chakraborty et al., 2022) |
| SGH_OsActin_ RT_F | 2929 | atgaagtgcgacgtggatattag | RT-qPCR | This study |
| SGH_OsActin_ RT_R | 2930 | gggcgaccaccttgatcttc | RT-qPCR | This study |
| AP_BS_Os01g0 229300_F1 | 4991 | ATTTTTTGTGTTTTGGAG TTGTATT | Bisulfite-PCR | This study |
| AP_BS_Os01g0 229300_R1 | 4992 | AACCATACTTTATATATAT TACCCACACTT | Bisulfite-PCR | This study |
| AP_BS_Os05g0 327100_F1 | 4997 | TAATTTTTGTTTTAGTTTA TGAATT | Bisulfite-PCR | This study |
| AP_BS_Os05g0 327100_R1 | 4998 | CCTATAATACATCCTCAA ATATCTTAC | Bisulfite-PCR | This study |
| AP_BS_Os05g0 327100_F2 | 4999 | GAGGTAGGTTATTAATAT GTTTTTTTT | Bisulfite-PCR | This study |
| AP_BS_Os05g0 327100_R2 | 5000 | AATATCTCAAACTCCTTT CCATATC | Bisulfite-PCR | This study |
| AP_siren2_BS_ F2 | 5025 | TTTAGTTGATTTAATGGTT AATATG | Bisulfite-PCR | This study |

| | | | | |
|---|---|---|---|---|
| AP_siren2_BS_R2 | 5026 | AAAAAACACTTTCATCTAAAATATC | Bisulfite-PCR | This study |
| AP_siren2_BS_F4 | 5029 | GGAGATATTTATTTGTTATGGGGAT | Bisulfite-PCR | This study |
| AP_siren2_BS_R4 | 5030 | AAAATCAAAAATTACCCTATTTACTCTACT | Bisulfite-PCR | This study |
| AP_ARF22_CRS_F1 | 5071 | GCACCAGCCCATGACAATATCTCTTGTTTG | Imprinting study | This study |
| AP_ARF22_CRS_R1 | 5073 | TGCTCAGTAAATATAGCCTTTCCAAACAGC | Imprinting study | This study |
| AP_SHH1_CRS_F2 | 5076 | GACCTTGTCCTCTGTTTCAAGGAGAGC | Imprinting study | This study |
| AP_SHH1_CRS_R2 | 5078 | CCATCATGGTCGTATTCGACAAGGAAGACAC | Imprinting study | This study |
| AP_CLSY3_CRS_F1 | 5079 | gagaaatgtaaagattgctcagaggaggaagc | Imprinting study | This study |
| AP_CLSY3_CRS_R1 | 5081 | caggaacaggctcatattggttgctctc | Imprinting study | This study |
| AP_WP/PB1_F | 3934 | tccacaatgcgtcagaaacaatg | Imprinting study | This study |
| AP_WP/PB1_R | 3937 | cagttcatcttcatcctgtaacttgtc | Imprinting study | This study |
| miRNA168 | 32 | gtcgccgagaagatcctccatc | sRNA northern | This study |
| U6_probes | 13 and 14 | ggccatgctaatcttctctgtatcgtt and ccaattttatcggatgtccccgaagggac | sRNA northern | This study |
| amiR_PolIV probe | 1697 | atgtccaagagtaacactata | sRNA northern | (Hari Sundar G et al., 2023) |
| MITE siRNA | 3430 | ggtcccacctgtcatacacacact | sRNA northern | This study |
| CACTA siRNA | 2086 | acgttcaatacaatctcggcgcgt | sRNA northern | (Hari Sundar G et al., 2023) |
| miRNA820 | 3343 | cctggtccatccacgaggccga | sRNA northern | (Nosaka et al., 2012) |
| Tos17 siRNA | 2091 | ggcctctacatccaaccggatat | sRNA northern | (Hari Sundar G et al., 2023) |
| AP_CLSY3_amiR1_probe | 3877 | GCGAGGCCTTATTGTATCGAA | sRNA northern | This study |
| AP_CLSY3_amiR2_probe | 3878 | AGGGCGGACTTCAATTGAACA | sRNA northern | This study |
| AP_CLSY4_amiR1_probe | 3880 | GAGAGGTCATTCGTGCTTACA | sRNA northern | This study |
| AP_siren_probe2 | 4751 | CCCGTGTGCAGGTTCTAGTTCCGG | sRNA northern | This study |
| AP_siren_probe3 | 4752 | CCGTGTGCAGGTTCTAGTTCCGG | sRNA northern | This study |

### References

**Chakraborty, T., Trujillo, J.T., Kendall, T., and Mosher, R.A.** (2022). A null allele of the pol IV second subunit impacts stature and reproductive development in Oryza sativa. Plant J. **111**: 748–755.

**Chen, C., Li, T., Zhu, S.[. ]., Liu, Z., Shi, Z., Zheng, X., Chen, R., Huang, J., Shen, Y., Luo, S., and Wang, L.** (2018). Characterization of imprinted genes in rice reveals conservation of regulation and imprinting with other plant species. Plant Physiology **177**: 1754–1771.

**Hari Sundar G, V., Swetha, C., Basu, D., Pachamuthu, K., Raju, S., Chakraborty, T., Mosher, R.A., and Shivaprasad, P.V.** (2023). Plant polymerase IV sensitizes chromatin through histone modifications to preclude spread of silencing into protein-coding domains. Genome Res.

**Nosaka, M., Itoh, J.-I., Nagato, Y., Ono, A., Ishiwata, A., and Sato, Y.** (2012). Role of transposon-derived small RNAs in the interplay between genomes and parasitic DNA in rice. PLoS Genet. **8**: e1002953.

Supplemental Table S4: Sequences of DRD1 family proteins used in this study

>LOC_Os02g43460

MPRRKGKGKGVEDEVEVYEPASPPERVLIILDSSEDDLDLQEVRRSLMITGRGRARAAERV
GEEAPRGSGRRAAPVVASRRRRRRSRSRSRSPRAARPRAESSRRPTARRARARARSPSL
EIIDVDSGSDRGVVRVKEEPRSGSDSDYNGARGRARARARAPVAATAAKKKKRKRGKEAP
SRAQESREVVRVKEEPNSDGNGAGGRARARSPVAAAAKQRKRGGREAPSRAQESRVPV
QIKEEPYSGSDSDGNVAGGRAVVPAADAKQGKRGKKTPSRGKGRRVVVRETSTPAAPSN
GAPSVGRGKGRGPGRGRQRSKGAVRGRATPVNRVSTGVGSRTRSRLAEQGRAFAQEEE
EQVEEREEEEEEEEEQGRAFAQVKEEQVEEQEEDEEEGEEEMEMEVEVEVRSDDNDHGN
GGIRGEGGGTDDVAEIEEEELGTDEDETSDDSDENFSDEEGDEEELEEEEEEEEEEEDDD
DDDEEEEEEPGDAPDQPGEAGEESPPRSRIMAMPLMGKRMFEGFSFLQQVDTSTGRDIRA
RTRSNFKRKKLLDKKLLKRGTFAKPYCIDVSSSGSEEDVPQPEQSAYGGDCADDDGGSDG
NEEHRAVKRRKLNRRQSAHSDSEEDTTFVCDVKEGSGSRRVQEGAPRRQVKKEGSNKKK
DGSTPQCVRNNGPKVGRQTNGLNGQGGVSFKRNVKIAQRRKRRQATADQEKYGHLLDP
MFNEIESNQYEPVPEEQIDRRLPLVFAFGDDDKLEEKSKHDKLQDEDELWKEFDFALESINV
CSHNCEEGEKEDEQEIPADKAASCIQGKHELIIDEQIGLRCKHCNFVDLEIRFVLPSMVKSCT
ERDMRKDHELDLFFDDILTSAGYEGPRDFGGKKTGLVWDLVPGVREDMFPHQQEGFEFM
WRKLAGGTSIEQLRNNANTIEGGCVISHAPGTGKTRLAITFVQSYFAFFPECCPVIIAPRGML
ATWEQEFRKWKVKVPFHVLNSKEINWKEDRTIKQLAIMDENLAQSLARNKLDHKFRRKLKL
ASWRKGSSIIGVSYTLFRKLANQSSMDGNMVRNLLLEMPDLLVLDEGHTPRNKKSLIWKVL
EEVRTKKRIILSGTPFQNSFLELSNVLYLIRPKFARHFASKSFKKIGLEDYWTSLTLNNITEKKI
DEIRQILDPIVHIHNGDILQKSLPGLRESVVILNPLPHQKEIITAMENTVTMGTLDAEYKISLASI
HPFLVTCAKLSEKETSSVDVSLLKSLRPNPCVGVKTKFVLEIVRLCEAMKERVLVFSQYLEP
LSLIMDQLSKMFNWIEGEEILLMSGNVLVQNREALMEAFNDMKSNAKVMLASTKACCEGITL
IGASRVVLLDVVWNPSVGRQAIGRAYRIGQEKIVYTYNLITEGTKEKDKYDRQAKKDHMSKL
LFSKEPHAAGFNLSQEVIFNDKILEAMTSHRELKDMFVKILHSH

>LOC_Os05g32610

MDRAARLARRGGGVTVAEYRMVRGRRRGGDAGPVVVIDVEDDGEDAADDSAGGGGGAA
AAVKRRVVVPGAVATRTRSRRMAMAQQAPVTPPAAAEEAPSRRRKRKGAASAEAGGGGP
SKRRVRSSGSAGGRGARKRKEAEADEEEAEAEAEEEAEAEAGTPARGESMEVSQVDGG
GSSGRADDASHNGNGESRVCNADGIDQASEERPSVAGGDLIEEEHYGNGEASVAGGDRIE
EHCGNVEASVANSNRDGGEIIAGEGTEDRGNTELSVVDPVNEELASDEDDYDDEMLEEKL
VGDVIRAYSNGADLDTNGVDWEAEDEMEFADLDTNVVDWEAEDEMEFDDDNDNDADDD
GDNFGGDADEGDKSVQMHDFSKVETQDLVSHNVNVSEVRPHEDEEAIKDEMESKGKGSL
SFNEGSSYIEILDSDEEVKVVNDTGNALRRKPLVPAKLPIVPSCVAWRTRSSWGMKEERISY
NTYFEVLSDEPKEDDDDTEVELDDEEDDENDDDCNSASCDEEDEEEEEEREEEEEEEAQRR
KQKKGIDSSDDEMIDDAVDCGIDWEEDYPEVDFTRPLTFQKDGSEAPVGSEAFTEQQKRS
RFTWELERRKKLKLGMMTNHRLYERDLESDSNSSDSSQNRKNGCQGSGDHRTGRKRKN
PLSKSGKKSSRMLKRQSLMKLLMDKMCSNDDGKSTPFDQKPQIEYSFKDLHPLVFSFGDD
DPSPTDRSEQDAALDMLWADLDFTLESENIGTYYDDEGQEDSLLDHALAPITPCSRGKHEFI
IDEQIGIRCKYCSLVNLEIRFILPLLASNFAEKPAWRNSSCLKTALMCPDLYEQTGTGDGQSQ
DFHINGTVWDLIPGVITDMYQHQREAFEFMWTNLVGDIRLNEIKHGAKPDVVGGCVICHAP
GTGKTRLAIVFIQTYMKVFPDCRPVIIAPRGMLFAWEQEFKKWNVNVPFHIMNTTDYSGKED
RDICRLIKKEHRTEKLTRLVKLFSWNRGHGVLGISYGLYMKLTSEKVGCTGENKVRTILLENP
GLLVLDEGHTPRNERSVIWKTLGKVKTEKRIILSGTPFQNNFLELYNILCLVRPRFGEMFLTK
TRVGRRHCVSKKQRDKFSDKYEKGVWASLTSNVTDDNAEKVRSILKPFVHIHNGTILRTLG
RECVIVLKPLPLQKSIIRKVENVGSGNNFEHEYVISLASTHPSLVNAINMTEEEASLIDKPMLR
LRSNPYEGVKTRFVMEVVRLCEALKEKVLIFSQFIQPLELIKEHLRKIFKWREGKEILQMDGKI

LPRYRQNSIEVFNNPDSDARVLLASTRACCEGISLTGASRVVLLDVVWNPAVGRQAISRAF
RIGQKKFVYTYNLITYGTGEGDKYDRQAEKDHLSKLVFSTEDEFSNVRNMLSKAEMEHCSK
LISEDKVLEEMTSHDQLKGMFLKIHYPPTESNIVFTYNQIAPELS

>LOC_Os07g49210

MLMLMDPPARSRRCTLLTRALLLAVAALALRLIYAAFLAGMALYPPLPAAAVLGSKTYLHSA
VATPDAWRTRDWRKAVDYHATLLAPHLADGILSPTSRAVCLGAVQEALAMRELGVSTAVA
VARKRSPPLAVAGNDRRLPFQDSSVDFVFAARALDSSKRPADLAAESARILKPDGHLVVLT
TSAADAFSLRALQALLPSLRLLRSRQIKGPDDSTLRELVFQKIQDSTDDPVNKCTIGDHKLQL
LTHAEPLIQEEPRKPWITLKRNIKNIKYLPTLADISFKRNYVYVDVGARSYGSSIGSWFRKHY
PKQNHTFQVFAIEADPAFHSEYAAKKAVTLLPYAAWVKNETLNFEINADPGKEDEAKANGR
GMGRIRPMAGKKMSGEVRSVPAFDFAEWLKRTVSEQDYVVMKMDVEGTEFDLIPRLFDTG
AICLIDELFLECHYNRWQKCCPDRAEAFEMAKGVSCFYWSIQFPNFKDHLCFRNCSNASST
RHFSYRSLLIRTEKPVTTRNHAYAEVVVFVLDQNPMFFLFLRFFYPAIQRGPNCWSSANSTV
MRQAFEVFYDGSWHGVNCIRIRNGNLFVKFIYSGSTVEHNVDGDCLRLRSRRATCSDCSN
VLKPGVDVCVQSSHTPEASSQGGTNASVLLRHDARLITIKKNHQEDKCLCLFVVILYKNQCP
GNAEKVITDRRAEVVTINDIFLLQKLQPEVHEGSMKWSFSKDRLSLNKGRLISARFSSEITHLI
VLSILRGMEFNIKLVEGQIVYQIIKGDQAQWNLDSMAIPPGFGNTMEIISFQLRDEALRPTITNI
PITHVKKNNITEDMRFTVKSEMDSELDRALDVEILYEHVDLRRSKRLKTQPDRFTSYDTPRF
LSGYKKKEASSSPTKHVRGAVHCDSPVDDSKKEVESCCVEIPGNVTQKQTGVHSPMVDEK
SNSPEGQHKNTTKRTTCSLVKEKASSPEGQHEKTTKRTTCALPVKEKASSPEGQHKNTIKR
TTCSLPVKEEPSSVEIEEKSSKEQSAPEFHIPRTPAQNKEKHNRPPFSCKPKLFTSSGTLGV
NCEPAFCQKVGGKRKRHMCEREYKQMIDQCIGNIESEMERDSMFNFDANMMNYVQHSYR
EEDFTWPPSADNQEVEEDELEELWKEMDYSLTTLALLEQKQVMAQSRINMLVDNFDGLRL
DCLTLTDDYRCYYQKKEKFAESGSVNESTDYFGKVGGIPCHHECILDEELGLACRLCNVVC
TEAKDIFPEMFNGNDYKDRPGCSNICLDDDILDPSLLANLAPELSELKNSGSVWSAISDLDP
KLLPHQRKALDFLWKNLAGSIQVEGMDNSNVSTGGCVIAHTPGSGKTLLLISFLVSYMKAHP
RSRPLVLTPKAAIHTWKREFEKWGISLPLHVFHHANRSGKPLGAMDSKLRSLLNNFHRPTW
TNMRLMDSLDKLFKWHAHPSVLLMTYSSFLGMTKQDSKVRNRYREFIAEVLMNNPGLLILD
EGHNPRSNKSKLRKLLMKVKTEFRILLSGTAFQNNFEEYFNTLCLARPRFIGDIMSELVPER
KRETVGRRAKHQEAVARRAFVEKVGQKIESDNKHIRSDGISLLNKLTRGFIDSFEGAKLINLP
GIHVYTVFMKPTDIQEEMLAKVTMPKLGSSRFPLEVELLITIGSIHPWLIKTTKAVSTFFSPAE
VKKVERYKRDFAAGCKAKFVIDLLHKSSFRGERVLIFCHNVSPITFLVKLIEMVFGWRLGEEV
LVLQGDQELPVRSDVMDKFNGDSAGKRKVLIASTTACAEGISLTGASRLVMLDSEWNHSKT
RQAIARAFRRGQERTVYVYLLVASGTWEEEKYNSNRRKAWMSKMVFLGRYVDDSSQNRV
TDIDDEVLKELADEDHTGTFHMIVKQD

>LOC_Os03g06920

MARYPAPTSSRAIGAPIQPTEPHAPLPNTGGEGAPPPARTMPPPSSQAATSTPPAAATPLQ
RPPAQATAQPSTQRYYVGVQRDKGTGKWAACVVDPSNPTKHRLVGAFPDEHAAALAHDR
LDLAFRGGGHRGAGDNFRPAFHAVELEFLRLCAATSSPGSHCGLVAGGDKYDEKYSEFLR
KIYHGVMDNSPSYKKFFDVILDFFIARAREIGREALEDGGDMLVERFVAMHKNKAVTPRWR
AWYRSDSRKVLQIPLSLRGGGGEIDHSTQKEARMDSDSCKRRKHESGHDSSSRVQSQSSI
LSRNRILCHQLLEQCDDLKYGSSTNDYKAISMKRLELISILQKLQEVPIQLPYASPLKSSETNR
LVQDGRNSSCRNIIDLDSDNDEDYTFANVDNIGANTTVVLVDSDDGDSVASFVDEKSSDSK
QNANYIEESVLPEQHAQQQEISMLDNENISSEAQAVKKGKDSMDINDVIYNKSGHEEIGEEE
AQAENVQIKGNLKKEIISVASDELACEVMRSQSPTNGNFDQYDNSSPVDELEGLWMDMYL
AMACSKTVGSDHNIVPSENSCEQAEDECQHDFLMKDDLGIVCRVCGLIQQRIENIFEYQWK
KRKQSYRARPSEHRNSSDADAIDKTSGAILEVVPDALCLHPQHSQHMKPHQVEGFNFLVK
NLADENNPGGCILAHAPGSGKTFLIISFVHSFLAKYPAGRPLIILPKGILSTWRTEFLHWQVD

DIPLYDFYSSKADKRSEQLKVLNLWEESRSILLLGYQQFACIVSDHTSDTEAIMCQEKLLKVP
SLVILDEGHTPRNEETDLLTSLENIRTPRKVVLSGTLFQNHVREVFNILKLVRSKFLKMDKSR
AIVNCILSKVDLMGKSARSKNISDKDFFDLVQEHLQKDGNDKMRAVIIQNLRELTADVLHYY
QGKLLDELPGIVDFTVFLNMSSKQEHIIKGLDGINKFAKRSRCNAVSLHPCLKNANKADADD
GNVTNRKIGSIISGIDINDGVKAKFVHNLLSLSEATGGKVLVFSQYVRSLIFLEKLVSRMKGW
KSEVHIFRVTGGSTQDQREQAVHRFNNSPDARVFFGSIKACGEGISLVGASRIVILDVHENP
SVMRQAIGRAYRPGQSKMVYCYRLVAADSPEEDDHHTAFKKERVSKLWFEWNELCSSDD
FELATVDVSDSEDRFLESSALKQDIKALLKR

>LOC_Os08g14610

MSGSGNSLDTVALIVGGGSDSSGIVGRKRRRCDLIRERWCCLCPVWCKEAQEVVVPGRG
RNGARQRDGGGCALGTTEVLGRICNSSVEKAEERETVIPAISNTEKMGEKQQKSIPRDRKR
KGELDPAADYVKDLWDAFYVTAESTHLDTSEVNNKKQLDNCNHDIHVYEDLGHVCHECGL
VVRKADSLFHYQWKKASRKRTNVNEVCLKKVGSDAISLSEDFIFSDIAIHPRHAKNIRPHQLE
GFKFLVNNLVTDEPGGCILVHAPGSGEIFMLISFIQGFMARHFTARPLVVLPEGILGTWKREF
QQWQVEDIPLYDFD

SIKADNRVEQLEVLKSWSSKRSILFVGSKHFTQIVCDDRDENAVAECRDTLLMVPSLLILDE
GHTPSIDETDMLQSARKVQTPCKVVMSGTLFHNHVKEVFNTLDLVRPGFLKTETFWPIVTR
MMGQLEISSARSITEISESMEDTLLNDDNFTRKVNVIRSLGELTKDVLHYCKGEDLNEFPVLL
DFSVFLELSPKQKDILCKLEEDHGMLKTSAVGAALYVHPCLSEISEANDVDRDDRVDSLVNS
INLGDGVKARFFLNILALANSAGEKLVAFSQYTLPMKFLERLLVKEMGWHVGKEIFVINGDTS
MEDGQLAMDQFNGSADAKVLFGSIKAFGEGISLVGASRIVILDVHLNPSVTRQAIGSTFRPG
QKKKVFVYRLVAADSPEEKAHETAFNKEVIPKLWFQWSGRCTTEDFKLNQVCIDGSRDELL
ETDVIRQDIKALYQSIDMGLVSEATVCFNNVSSSGLSVHDTGGNVIGQGDQDSEKNRYLSIA
SETMLVHFVHVFSFVCVPVNTTCSLSICREILWFLIGTREVPFSSVKFGISSWMGKITLIRIFRI
LYDEKGQIALLASKGSIKDKQEACKSWRYPSIHPWLTEAVVLAVTAATPPHLLAVAGPLPLP
SLCSAVPFALLYYFGKGRKVVHTLKNWLQQVSVENKICGWGYNTTEVLGRICNCSVEKAEE
RETIILASGNMEKMEEKHQKSDQDFHFSDSTMAIPRERKQKGEVDPAADCLKDRWGAFYV
AVESTQLDTSEVNNKKQLNNYNHDIHVYEDLGRVCHECGSGEIFMLISFIQGFMTRHSTARP
LGTWKREFQQWQVEHCSGFIYL

>LOC_Os08g19250

MAGFADALRPDKFTSVHFKRWQIRVNLWLTAMKCFWVSTGKPMGVLNADQQKEFDEATT
LFVGCILSVLGDRLVEVYMHMTDATELWDALNTKFGATDASNDLYIMEQFHVYKMADNRSV
VEQAHKIQTMAKELKLLKCVLPDKFVAGCIIAKLSPSWRGFGTALKHKRQEYFVEGLIASLDV
EEKAREKDVASKGDGGQSSTNVVHKAQNKSKGKYKAQQTTNFNKKKKNNNPNQDERTCF
VCGQPGHLARKCPQRKGMKAPAGQTSKSANVTIGNTGDGSGYGRTGFHRPNGEWVTCF
CSWCWHGRSEVYFGKDRAAEERVAYYWKEDVRSEMDSIIANGTWEVTERPYGCKPVGCK
WVFKKKLRPDGTIEKYKARLVAKGYTQKEGEDFFDTYSPVARLTTIRVLLSLAASHGLLIHQ
MDVKTTFLNEELDEEIYMDQPDGFVVEGQEDKVCKLLKSLYGLKQAPKQWHEKFDKTLTSA
GFAVNEADKYVYYRHGGGEGVILCLYVDDILIFRTNLEVINEVKSFLSQNFDMKDLGVADVIL
NIKLIRGENGITLLQSHYVEKILNRFDYIDSTMELGLHYSGYPAVLEEYSGSNWISDVDEIKAT
SGYVFTLGGGAVSWRSCKQTILMRSIMEAELTALDTATVEAEWLRDLLMDLPIVEKPVPAIL
MNCDNQTIIVKMNSSKDNMKSSRHVKRRLKFVRKLRNSGVITLDYIQTARNLANPFTKGLSR
NVIDNASKEMGLRPISNDTLMLLYICNVMIYLWNILNHYMPIFSTFKRHEIEAYEKFKRSAVGT
ALYIHPCLSEISEGDAADRASNLTDATVDSLIESIIIKDGVKAKFFFNIMSLANSAGEKLLAFSQ
YILPMKFLERLLVKRLGWHVGKEIFMISGDTSADDREVAMDQFNNSADAKVLFGSIKACGE
GISLVGASRVIILDVHLNPSVTRQAIGRAFRPGQQKKVFVYRLVAADSPEVKFHETAFKKEVP
KLWFEWSELCTTEDFKLNQVDIDDSEDELLEANAIRQDIKALYRRMLQ

>LOC_Os06g14440

MYYRRQRKASSEANANVFMPGGPNDISFPASNRDHDWGYGGVGKEWEASYARKLQLMN
FLSSLHQRTANPLVTTRMDANMDTPLEQKQKDSSAIIVLDSDDEDGYTEGCEQLTSENNKQ
QAPSGLTSPYTTWIVSSAKDQVNGTLHVDGVQSTQIVPYYGQNAPLINQFPLQTSWQPSIQ
YERVILQKRPEEQRVQDLVAASHAEKIAETQVLLTLPTLPNERKRRKTEPTTLVDVDGGTNL
GKRKRKNHQNQAAVDSNLDLQQNDVPSQSYRTMIEEEKPVKESDGLEDLWKDFSLAAECT
KLDTNEDMSNEKDVDDENEMDDDCNHDIRIHEDLGHVCRICGMIVRKAETIIDYQWKKASR
TRTNYYESRSKDADEIDTGAVKVSEDFIVSDIAIHPRHAKQMRPHQLEGFSFLVKNLVGDKP
GGCILAHAPGSGKTFMLISFIQSFLAKYPSARPLVVLPKGILGTWKREFQRWQVEDIPLYDFY
SVKADKRVEQLEVLKSWEAQMSILFLGYKQFSRIICGDGDGNIAAACRDRLLMVPNLLILDE
GHTPRNRETDVLASLKRVQTPRKVVLSGTLFQNHVSEVFNILDLVRPKFLKMESSRPIARRI
MSQVAISGIRSLKGVHDSAFTESVEDTLLNDDNFTRKAHVIRSLRELTKDVLHYYKGDILDEL
PGLVDFSVFLKLSTKQKEIVHKIEAYEKFKRSAVGTALYIHPCLSEISEGDAADRASNLTDAT
VDSLIESIIIKDGVKAKFFFNILSLANSAGEKLLAFSQYILPMKFLERLLVKRLGWHVGKEIFMIS
GDTSADDREVAMDQFNNSADAKVLFGSIKACGEGISLVGASRVIILDVHLNPSVTRQAIGRA
FRPGQQKKVFVYRLVAADSPEVKFHETAFKKEVIPKLWFEWSELCTTEDFKLNQVDIDDSE
DELLEANAIRQDIKALYRR

>AtCLSY3(AT1G05490)

MECIGKRVKSRSWQRLQAVNKRKKMETVAPVTSPPKKRRQKKPKNYDSDIEDITPTCNDS
VPPPQVSNMYSVPNNSVKESFSRIMRDLNVEKKSGPSSSRLTDGSEQNPCLKERSFRVSD
LGVEKKCSPEITDLDVGIPVPRFSKLKDVSEQKNTCLMQKSSPEIADLDLVISVPSSSVLKDV
SEEIRFLKDKCSPEIRGLVLEKSVPGEIEILSDSESETEARRRASAKKKLFEESSRIVESISDG
EDSSSETDEEEEENQDSEDNNTKDNVTVESLSSEDPSSSSSSSSSSSSSSSSSSSSDDESY
VKEVVGDNRDDDDLRKASSPIKRVSLVERKALVRYKRSGSSLTKPRERDNKIQKLNHREEE
KKERQREVVRVVTKQPSNVVYTCAHCGKENTGNPESHSSFIRPHSIRDEIEDVNNFASTNV
SKYEDSVSINSGKTTGAPSRPEVENPETGKELNTPEKPSISRPEIFTTEKAIDVQVPEEPSRP
EIYSSEKAKEVQAPEMPSRPEVFSSEKAKEIQVPEMPSIPEIQNSEKAKEVQANNRMGLTTP
AVAEGLNKSVVTNEHIEDDSDSSISSGDGYESDPTLKDKEVKINNHSDWRILNGNNKEVDLF
RLLVNSVWEKGQLGEEDEADELVSSAEDQSQEQAREDHRKYDDAGLLIIRPPPLIEKFGVE
EPQSPPVVSEIDSEEDRLWEELAFFTKSNDIGGNELFSNVEKNISANETPAAQCKKGKHDLC
IDLEVGLKCMHCGFVEREIRSMDVSEWGEKTTRERRKFDRFEEEEGSSFIGKLGFDAPNNS
LNEGCVSSEGTVWDKIPGVKSQMYPHQQEGFEFIWKNLAGTIMLNELKDFENSDETGGCI
MSHAPGTGKTRLTIIFLQAYLQCFPDCKPVIIAPASLLLTWAEEFKKWNISIPFHNLSSLDFTG
KENSAALGLLMQKNATARSNNEIRMVKIYSWIKSKSILGISYNLYEKLAGVKDEDKKTKMVR
EVKPDKELDDIREILMGRPGLLVLDEAHTPRNQRSCIWKTLSKVETQKRILLSGTPFQNNFLE
LCNVLGLARPKYLERLTSTLKKSGMTVTKRGKKNLGNEINNRGIEELKAVMLPFVHVHKGSI
LQSSLPGLRECVVVLNPPELQRRVLESIEVTHNRKTKNVFETEHKLSLVSVHPSLVSRCKIS
EKERLSIDEALLAQLKKVRLDPNQSVKTRFLMEFVELCEVIKEKVLVFSQYIDPLKLIMKHLVS
RFKWNPGEEVLYMHGKLEQKQRQTLINEFNDPKSKAKVFLASTKACSEGISLVGASRVILLD
VVWNPAVERQAISRAYRIGQKRIVYTYHLVAKGTPEGPKYCKQAQKDRISELVFACSSRHD
KGKEKIAEAVTEDKVLDTMVEHSKLGDMFDNLIVQPKEADLVEGFSILMP

>AtCLSY4(AT3G24340)

MDMTSCVARRTRSRTESYLNSILNKSKGISGEEEDQSLGCVNSRTEKRRVNMRDACSPSP
RKKKRRRRKDDDDDVVFVRTEYPEGKRDDENVGSTSGNLQSKSFDFGDRVCDFDADDRN
LGCEEKASNFNPIDDDDDVVFVGTVQRENDHVEDDDNVGSASVISPRVCDFDEDDAKVSG
KENPLSPDDDDDVVFLGTIAGENQHVEDVNAGSEVCDILLDDANLRGEEKTYVSDEVVSLS
SSSDDEEDPLEELGTDSREEVSGEDRDSGESDMDEDANDSDSSDYVGESSDSSDVESSD
SDFVCSEDEEGGTRDDATCEKNPSEKVYHHKKSRTFRRKHNFDVINLLAKSMLESKDVFKE

DIFSWDKIAEVDSREDPVVRESSSEKVNEHGKPRERRSFHRVREKNHLNGESFYGGEKLC
DGEETINYSTEDSPPLNLRFGCEEPVLIEKTEEEKELDSLWEDMNVALTLEGMHSSTPDKN
GDMLCSKGTHDFVLDDEIGLKCVHCAYVAVEIKDISPAMDKYRPSVNDNKKCSDRKGDPLP
NRLEFDASDPSSFVAPLDNIEGTVWQYVPGIKDTLYPHQQEGFEFIWKNLAGTTKINELNSV
GVKGSGGCIISHKAGTGKTRLTVVFLQSYLKRFPNSHPMVIAPATLMRTWEDEVRKWNVNI
PFYNMNSLQLSGYEDAEAVSRLEGNRHHNSIRMVKLVSWWKQKSILGISYPLYEKLAANKN
TEGMQVFRRMLVELPGLLVLDEGHTPRNQSSLIWKVLTEVRTEKRIFLSGTLFQNNFKELSN
VLCLARPADKDTISSRIHELSKCSQEGEHGRVNEENRIVDLKAMIAHFVHVHEGTILQESLPG
LRDCVVVLNPPFQQKKILDRIDTSQNTFEFEHKLSAVSVHPSLYLCCNPTKKEDLVIGPATLG
TLKRLRLKYEEGVKTKFLIDFIRISGTVKEKVLVYSQYIDTLKLIMEQLIAECDWTEGEQILLMH
GKVEQRDRQHMIDNFNKPDSGSKVLLASTKACSEGISLVGASRVVILDVVWNPSVESQAIS
RAFRIGQKRAVFIYHLMVKDTSEWNKYCKQSEKHRISELVFSSTNEKDKPINNEVVSKDRIL
DEMVRHEKLKHIFEKILYHPKKSDMNTSFF

>AtCLSY2(AT5G20420)

MKKRGFYNLKHPFDPCPFEFFCSGTWKPVEYMRIEDGMMTIRLLENGYVLEDIRPFQRLRL
RSRKAALSDCICFLRPDIDVCVLYRIHEDDLEPVWVDARIVSIERKPHESECSCKINVRIYIDQ
GCIGSEKQRINRDSVVIGLNQISILQKFYKEQSTDQFYRWRFSEDCTSLMKTRLSLGKFLPD
LSWLTVTSTLKSIVFQIRTVQTKMVYQIVTDEEGSSSTLSSMNITLEDGVSLSKVVKFNPADIL
DDSQDLEIKQETDYYQEEDEVVELRRSKRRNVRPDIYTGCDYEPDTIDGWVRMMPYQFGK
CAVNVESDEDEDDNNEDGDTNDDLYIPLSRLFIKKKKTNSREAKPKSRKGEIVVIDKRRVHG
FGRKERKSELSVIPFTPVFEPIPLEQFGLNANSFGGGGSFSRSQYFDETEKYRSKGMKYGK
KMTEMEEMMEADLCWKGPNQVKSFQKRTSRSSRSVAPKTEDSDEPRVYKKVTLSAGAYN
KLIDTYMNNIESTIAAKDEPTSVVDQWEELKKTNFAFKLHGDMEKNLSEDGEGETSENEML
WREMELCLASSYILDDNEVRVDNEAFEKARSGCEHDYRLEEEIGMCCRLCGHVGSEIKDVS
APFAEHKKWTIETKHIEEDDIKTKLSHKEAQTKDFSMISDSSEMLAAEESDNVWALIPKLKRK
LHVHQRRAFEFLWRNVAGSVEPSLMDPTSGNIGGCVISHSPGAGKTFLIIAFLTSYLKLFPG
KRPLVLAPKTTLYTWYKEFIKWEIPVPVHLIHGRRTYCTFKQNKTVQFNGVPKPSRDVMHVL
DCLEKIQKWHAHPSVLVMGYTSFTTLMREDSKFAHRKYMAKVLRESPGLLVLDEGHNPRS
TKSRLRKALMKVGTDLRILLSGTLFQNNFCEYFNTLCLARPKFIHEVLMELDQKFKTNHGVN
KAPHLLENRARKLFLDIIAKKIDASVGDERLQGLNMLKNMTNGFIDNYEGSGSGSGDALPGL
QIYTLVMNSTDIQHKILTKLQDVIKTYFGYPLEVELQITLAAIHPWLVTSSNCCTKFFNPQELS
EIGKLKHDAKKGSKVMFVLNLIFRVVKREKILIFCHNIAPIRMFTELFENIFRWQRGREILTLTG
DLELFERGRVIDKFEEPGNPSRVLLASITACAEGISLTAASRVIMLDSEWNPSKTKQAIARAF
RPGQQKVVYVYQLLSRGTLEEDKYRRTTWKEWVSCMIFSEEFVADPSLWQAEKIEDDILRE
IVGEDKVKSFHMIMKNEKASTG

>AtCLSY1(AT3G42670)

MKRKHYFEFNHPFNPCPFEVFCWGTWKAVEYLRIENGTMTMRLLENGQVLDDIKPFQRLRI
RSRKATLIDCTSFLRPGIDVCVLYQRDEETPEPVWVDARVLSIERKPHESECLCTFHVSVYID
QGCIGLEKHRMNKVPVLVGLNEIAILQKFCKEQSLDRYYRWRYSEDCSSLVKTRLNLGKFLP
DLTWLLVTSVLKNIVFQIRTVHEKMVYQIVTDEDCEGSSSSLSAMNITVEDGVVMSKVVLFN
PAEDTCQDSDVKEEIEEEVMELRRSKRRSGRPERYGDSEIQPDSKDGWVRMMPYRYNIW
NVSSDDDDEEEDCEDDKDTDDDLYLPLSHLLRKKGSKKGFSKDKQREIVLVDKTERKKRKK
TEGFSRSCELSVIPFTPVFEPIPLEQFGLNANSLCGGVSGNLMDEIDKYRSKAAKYGKKKKK
KIEMEEMESDLGWNGPIGNVVHKRNGPHSRIRSVSRETGVSEEPQIYKKRTLSAGAYNKLI
DSYMSRIDSTIAAKDKATNVVEQWQGLKNPASFSIEAEERLSEEEEDDGETSENEILWREM
ELCLASSYILDDHEVRVDNEAFHKATCDCEHDYELNEEIGMCCRLCGHVGTEIKHVSAPFA
RHKKWTTETKQINEDDINTTIVNQDGVESHTFTIPVASSDMPSAEESDNVWSLIPQLKRKLH
LHQKKAFEFLWKNLAGSVVPAMMDPSSDKIGGCVVSHTPGAGKTFLIIAFLASYLKIFPGKR

PLVLAPKTTLYTWYKEFIKWEIPVPVHLLHGRRTYCMSKEKTIQFEGIPKPSQDVMHVLDCL
DKIQKWHAQPSVLVMGYTSFLTLMREDSKFAHRKYMAKVLRESPGLLVLDEGHNPRSTKS
RLRKALMKVDTDLRILLSGTLFQNNFCEYFNTLCLARPKFVHEVLVELDKKFQTNQAEQKAP
HLLENRARKFFLDIIAKKIDTKVGDERLQGLNMLRNMTSGFIDNYEGSGSGSGDVLPGLQIY
TLLMNSTDVQHKSLTKLQNIMSTYHGYPLELELLITLAAIHPWLVKTTTCCAKFFNPQELLEIE
KLKHDAKKGSKVMFVLNLVFRVVKREKILIFCHNIAPIRLFLELFENVFRWKRGRELLTLTGDL
ELFERGRVIDKFEEPGGQSRVLLASITACAEGISLTAASRVIMLDSEWNPSKTKQAIARAFRP
GQQKVVYVYQLLSRGTLEEDKYRRTTWKEWVSSMIFSEEFVEDPSQWQAEKIEDDVLREIV
EEDKVKSFHMIMKNEKASTGG

>AtDRD1(AT2G16390)

MGFVYIVMTGYYKNVHKRKQNQVDDGPEAKRVKSSAKVIDYSNPFAVSNMLEALDSGKFG
SVSKELEEIADMRMDLVKRSIWLYPSLAYTVFEAEKTMDNQQVVEGVINLDDDDDDDTDVE
KKALCVVPSSSEIVLLDSDDEDNERQRPMYQFQSTLVQHQKNQGDVTPLIPQCSFEEVDLG
RGKEMPSAIKAIVEGQTSRGKVLPIENGVVNEKGVYVGVEEDDSDNESEAADEDLGNIWNE
MALSIECSKDVARETSHKEKADVVEDCEHSFILKDDMGYVCRVCGVIEKSILEIIDVQFTKAK
RNTRTYASETRTKRFGESDNELKFSEEGLMIGGLAAHPTHAAEMKPHQIEGFQFLCSNLVA
DDPGGCIMAHAPGSGKTFMIISFMQSFLAKYPQAKPLVVLPKGILPTWKKEFVRWQVEDIPL
LDFYSAKAENRAQQLSILKQWMEKKSILFLGYQQFSTIVCDDTTDSLSCQEILLKVPSILILDE
GHTPRNEDTNLLQSLAQVQTPRKVVLSGTLYQNHVKEVFNILNLVRPKFLKLDTSKSAVKRI
LAYTPCDVRGRLTGSNSDMASMFNETVEHTLQKSEDFTVKIKVIQDLREMTKKVLHYYKGD
FLDELPGLADFTVVLNLSPKQLNEVKKLRREKRKFKVSAVGSAIYLHPKLKVFSDKSDDVSD
TTMDEMVEKLDLNEGVKAKFFLNLINLCDSAGEKLLVFSQYLIPLKFLERLAALAKGWKLGK
EVFVLTGNTSSEQREWSMETFNSSPDAKIFFGSIKACGEGISLVGASRILILDVPLNPSVTRQ
AIGRAFRPGQKKMVHAYRLIAGSSPEEEDHNTCFKKEVISKMWFEWNEYCGYQNFEVETID
VDEAGDTFLESPALREDIRVLYKR

>GRMZM2G178435(RML1)

MPAPPSTEAGRSRTMTRVIILLDSDKEDDGTGRQAGRELGGAAIASAGEASKLVKPEVVDD
VGSNPVRPGALPTSLRVQGHRAPSSPSPVPAAVRKQPEIIAISDEDNDGSRFRRVRRVKDE
ASDWVLSAKAKRAMVSGVPPGSSDVKRKRKRGSSGAGDFHALDRNLSASGAGRRTSWM
AEDAGSSRNVSSSELSRGGVGDRSGSTKKARGAPGKTRRGGGTRRERSTSAAPANLVGG
SATVGSRIRLRSRQQGRVQCATYSARVSSEDTGEDEKHMQEQTRVEDVEFMEVDDDYDD
VNVAGNVIDQESEQDEALEGRSSQDSHGYSEDKEGKDSAALSDNEEDVGGKELLEEEEEG
ADQEESHIIYDGEGEQEEDASEEETQELDETGEAQPFNPSNTMAGSTMRSGGDGKQVFR
RRVFEGIYLPENPHRTVGKGIQGRTRSQRKCKDKKLLKRGTFSKPYNIDIPDSTSDSEEEIEP
PAPQQGLLSSSEEDNMTFGKRKRRRAAINKRWDKRLSASSDEEDYGASAMDAKERPFRR
LKKGLSNLQAAKEGCRNYEGSNPGHARYSGPNGGNLENMSSAQDDISFKRNVHMIRIKKR
GRAAKAVYDELLDSLFSGWENHIGNPVHAEAGNSLPLVFSFGDEDAEENTENDKYQEQED
LWMECGIAFQSMNIGSNGCEEDGKEIPPVKVTSCNIGQHEFIIDEQIGVRCKHCHVVDLEIR
DVLPTLGKCSAERGSAINPEFDRMLKEMLNVFEQNDVLVSNGHELPCNFGDHKAGSVWNL
IPGVKETMFPHQQDAFEFMWTKLAGGTTIEQLKHTIKSDAGGGCVISHAPGTGKTRLAITFV
QSYLEVFPHCSPVIIAPRGMLATWEKEFRKWKATGEARVLDERKLANHEGMDGDKVRKLLL
EKPNLLVLDEGHTPRNKKSLIWKVLKRVHTEKRIILSGTLFQNNFEELYNTLRLVRPKDADAL
HLETDESKDFWSSLRLNDITKANINEVRKKLDPIVHIHSGRFLQKSLPGLRESVVILNPLLYQK
EVIASMEKTVAMGLDAEYKISLASIHPSLLASAKLSMKEESILDKPKLESLRSNPSGGVKTRF
VLEIVRLCEALNERVLVFSQYLEPLSLIMEQLKERFSWAEGEEILLMSGKVLVKKRQTMMEV
FNNMKSKAKVMLASTKACCEGITLVGASRVVLLDVVWNPSVGRQAIGRAYRIGQRKIVYTY
NLIAEGTTEKRKYDRQAKKEHMSKLLFSNELERGGCNLPPELTFNDRVLEELTAREDLKNLF
V

>GRMZM2G154946(RMR1)

MDRATPRVCGRRGVSQAAVEAAPSSSRARRRDKAPAVVMDLGDDDCGGGGARKTVGGA
AGRCEGSTKAPLPLLPPMMVPAGAVALRTRSRRRAMLAAAVVEEAPTKKKKKEGAIPDAAE
APRGHGSKAAATSMATSSHKRRAGTSRSTSRDKRRARSGRASEPARVGRARKRKRNELE
APARRERVKAPCVSESDDNSGRGDDASHDGDAEPRVGVAIGTDLVNGDHPAAKEVVEGA
GDEDTGDGGNSGLASTADVFAEEMAPFEDDYDDEMLEEQLVGDVIRAYSNGRNFDSDGV
DWEAEDEMEFNDDADNSDFMDDADDSDFMDDAYEGGNSKPIQNHAKLEIQDWVNQKVVL
SGGRCEARGEGDLEEELDVGKEADEEDVEPKSEAAPGSDKRVLQLEILGSDEEIKVLENMS
SAPSRKASVQSKLPTIPSCVAWRTRSSWGVNQDRLSYDTYFEELSDEPKEDDDDTEVELD
EVEDDNNDDDSSDAYDKDDEEKEEEEEEEAERRKLNNRICTSDEDMINITVPTSRYDMFKKK
NSSRYDIEWVEDEDASVDMLQPVSFKKDSSWKPVAVGNDTFTEQQKRSRFTWELERRKK
LKLEMKTNPLHERDLDSDPNSSGSDQIRKYGFKSDGSHKVDRKKKHTSPKSGKKPSSAIILK
RQSLLKLLVDKMSGDKSLASFPFDQNPQLQFIFKEMHPLVFSFGDEDLVAADRPEQDVGLD
MLWADFDFALESENIGTYYDDECQEGNQLDFSLAPVTPCSRGKHEFVIDDQIGIRCKYCSLV
NLEIKFMFPSLVSVFAEKSAWPNDKGVKNTLMFHDLYEQGVNDTEQSQDIHQYGTVWNLIP
GVISTMYEHQREAFEFMWTNLVGDIRLDEIKHGAKPDVVGGCVICHAPGTGKTRLAIVFIQT
YMKVFPDCRPVIIAPRGMLFAWDEEFKKWNVDVPFHILNTTDYTGKEDREICKLIKKEHRTE
KLTRLVKLLSWNKGHGILGISYGLYTKLTSEKPGCTEENKVRSILLDNPGLLVLDEGHTPRNE
RSVMWKTLGNVKTEKRIILSGTPFQNNFLELYNILCLVRPRFGEMFLTKSRVGRRHYVSKKQ
KDKFSDKYEKGVWASLTSNVTDDNAEKVRSILKPFVHIHNGNILRTLPGLRESVIILKPLPLQK
SIIKKVENIGSGNNFEHEYVISLASTHPSLVTAINMSEEEASLIDKPMLAKVRSNPYEGVKTRF
VIEVVRLSEALREKVLIFSQFIQPLELIKEHLRKFFKWREGKEILQMDGKILPRYRQASIEAFN
NPNNDSRVLLASTRACCEGISLTGASRIVLLDVVWNPAVGRQAISRAFRIGQKKFVYTYNLIT
YGTGEGDKYDRQAEKDHLSKLVFSTEDEFNNVRNMLSKAEMEHCSKFISEDKVLEEMTSH
DQLKGMFLKIHYPPTESNIVYSYNQIATE

>GRMZM2G108166

MVKGSTGHHSNPIAPVLQHDIDGSYLRVRSRKATCSDCSHVLKPGADVCVWQAVYRGETK
DSVLLCCRDARLIKIKRNHQSDRCLCLFAVIFYKDQCPGSKEKVISGTIADVVTIDDICILQNLQ
PEELQDGSVRWNSAVDCFHHNRSKLLSARFSLEVAYLIVLSSLRRMEFNIKMVDGNIIYQIIK
GDQARDSIDSMSIPPGFGKNMDIISFKPRGEALRPITRTVPVTQVEEGNLTEDGCIAVKGES
DSAQDVEILYAHVDIRRSKRMKTQPDRFTSYDARNFNRTYNKKEADGPSTKYEDSESGLSC
DSSEQRESSDEEALENPRSMAAEHKYPVKRNQCSLPVKEKQISMEIKKNTTDQGCSDSYIP
HTPAKNTERPRFRLKPFASSRSLDGNSEPAFCQKRGRKRKKHMCQIEYKRMIDQCIGNIQC
EVERDSDFKFGDQILDGCVRAYQEVDFTWPSSADSQEEKDELDELWKEMDYALATVAILE
QKQMTDSEVVHESNTDLGKGGEHCHHDCMLDEQLGLTCRLCNVVCIEAKDIFPPMFTGKD
HERPERNHFGQDGHVLDLSFFEICAPEFSKIKESGNVWASITDLEPKLLAHQRKAFEFIWKN
LAGSLQLEEMDGSTSRGGCVVAHTPGAGKTLLLISFLVSYLKVHPRSRPLVLTPKAAIHTWR
TEFQKWGILLPLHVLHHSNRTSKLMGGLSSKLQAVLKSFHQPSWKTMRIMHCLDKLCKWH
EEPSILLMTYSSFLSLTKEDSKLRHQAFITKVLMNNPGLLILDEGHNPRSNKSKLRKLLMKVK
TEFRILLSGTVFQNNFEEYFNTLSLARPRFVNDVMTTLVTESEKRTRSRTGKHQEALARHVF
VERVGHKIESSSKHDRMDGISLLNELTQGFIDSFEGTKLNILPGIRVYTLFMKPTDVQEEVLA
KLLMPLSGNARYPLEYELLITIASIHPWLINTTKCASTYFTPAEVASVDKYKRNFAAGCKAKF
VIDLLHKSSFRGERVLVFCHNVAPIAFLVTLIEIVFGWRLGQEVLVLQGDQELHVRSDVMDK
FNSDRRGKRKVLIASTTACAEGISLTGASRLVMLDSEWNHSKTRQAIARAFRPGQERMVFV
YLLVASGTWEEDKYNSNRRKAWIAKMVFFGRHFDDPLQNRVTEIDDEVLKELADEDETNTF
HMIVKQD

>GRMZM2G393742(RML3)

MSQSPGGREGIYYSRQRKPSERNGSVFTPIAAMYPSGHALPDANRNHSLVFGGTSKDWD
NIRQFIASLERASENSSAIASKTGGGKSTNHSVEPAEQKGKGDIIVLDSDDEDGDGNSPEHN
KLASEMNKELGTSVLASNIAERMATNGSQTFETVHAYGGSKNTQIVPYGQGSALVNQFPLQ
TSWQPSIQFERVVLTKRPEEQRMQDLVAATIAEKRAETQMFLSLPTERKRRRTDHSLLMLD
SFVPKQRRRKGDTGLAPADLSLDLHQTATSQEPDIAIEEEEKRKNDGDGLEDYWKDFALAV
ESTKLDDVDEAAANEKEDNGKMEDIDCNHDIRIHEDLGHVCRVCGMIVRRADSIIDYQWKK
ASRRRMNGYGGNSKDADEIDCGTVKLSEDFIVADIAIHPRHAQAMKPHQVEGFNFLVKNLIG
DKPGGCILAHAPGSGKTFLLISFIQSFMARYPSARPLVVLPKGILVIWKKEIQRWQVQDIPVY
DFYSVKAEKRVEQLQILKSWEDKMGILFLGYKQFSTIVTDDGGSKVTAACRDRLLKVPNLLIL
DEGHTPRNKETDVLESLSRVETPRKVVLSGTLFQNHVEEVFNILNLVRPKFLRMESSRPIAR
RIMSQVEIFGRSSKGLADGAFTEAVEGTLLNDENFKRKVHVIRGLRELTRDVLHYYKGAILD
ELPGLVDFSVFLKLTPKQKDIVHKLEMHDRFKRSAVGSALYIHPCLSGLSEVNAENRAHTLR
DDSVDSLMDSINVRDGVKANFFMNILSLANSAGEKVLAFSQYILPMTFFERLLVKKKGWHV
GREIFMISGDTSQEDREAAVDRFNSSADAKVLFGSIRACGEGISIVGASRVVILDVHLNPSVT
RQAIGRAFRPGQQKKVFVYRLVAADSDEVKVHETAFKKEVIQKLWFEWSEQCTTENFKLG
QVDIDDSGDELLDTRAIRQDIKALYRR
